# HCR spectral imaging: 10-plex, quantitative, high-resolution RNA and protein imaging in highly autofluorescent samples

**DOI:** 10.1101/2023.08.30.555626

**Authors:** Samuel J. Schulte, Mark E. Fornace, John K. Hall, Niles A. Pierce

## Abstract

Signal amplification based on the mechanism of hybridization chain reaction (HCR) provides a unified framework for multiplex, quantitative, high-resolution imaging of RNA and protein targets in highly autofluorescent samples. With conventional bandpass imaging, multiplexing is typically limited to four or five targets due to the difficulty in separating signals generated by fluorophores with overlapping spectra. Spectral imaging has offered the conceptual promise of higher levels of multiplexing, but it has been challenging to realize this potential in highly autofluorescent samples including whole-mount vertebrate embryos. Here, we demonstrate robust HCR spectral imaging with linear unmixing, enabling simultaneous imaging of 10 RNA and/or protein targets in whole-mount zebrafish embryos and mouse brain sections. Further, we demonstrate that the amplified and unmixed signal in each of 10 channels is quantitative, enabling accurate and precise relative quantitation of RNA and/or protein targets with subcellular resolution, and RNA absolute quantitation with single-molecule resolution, in the anatomical context of highly autofluorescent samples.

**SUMMARY:** Spectral imaging with signal amplification based on the mechanism of hybridization chain reaction enables robust 10-plex, quantitative, high-resolution imaging of RNA and protein targets in whole-mount vertebrate embryos and brain sections.

## INTRODUCTION

RNA in situ hybridization methods (Harrison *et al*., 1973; Tautz & Pfeifle, 1989; Qian *et al*., 2004) and immunohistochemistry methods (Coons *et al*., 1941; Ramos-Vara, 2005; Kim *et al*., 2016) provide biologists with essential tools for elucidating the spatial organization of biological circuitry, enabling imaging of RNA and protein expression in an anatomical context. When developing these technologies for use in a given sample type, the first priority is to achieve a high signal-to-background ratio when mapping the expression pattern of a single target RNA or protein. Once this prerequisite has been met, other important priorities include the need for multiplex experiments to map relationships between circuit elements within a single specimen, the preference for the signal to be quantitative rather than qualitative, and the desire for the staining to be high-resolution to enable examination of spatial relationships at a subcellular level. In thin samples with low autofluorescence, it is straightforward to achieve all of these goals using probes direct-labeled with fluorophores (Kislauskis *et al*., 1993; Femino *et al*., 1998; Kosman *et al*., 2004; Chan *et al*., 2005; Raj *et al*., 2008). However, in whole-mount vertebrate embryos and other challenging imaging settings including thick brain slices, autofluorescence greatly increases the technical challenge of achieving high signal-to-background, motivating the development of in situ amplification methods (Qian *et al*., 2004; Ramos-Vara & Miller, 2014). For decades, the challenge of achieving high signal-to-background in thick autofluorescent samples proved sufficiently daunting that it was predominantly met by using enzyme-mediated catalytic reporter deposition (CARD) for both in situ hybridization (Tautz & Pfeifle, 1989; Harland, 1991; Lehmann & Tautz, 1994; Kerstens *et al*., 1995; Nieto *et al*., 1996; Thisse *et al*., 2004; Piette *et al*., 2008; Thisse & Thisse, 2008; Wang *et al*., 2012) and immunohistochemistry (Takakura *et al*., 1997; Sillitoe & Hawkes, 2002; Ahnfelt-Ronne *et al*., 2007; Fujisawa *et al*., 2015; Staudt *et al*., 2015), which came with unfortunate consequences. Using CARD, multiplexing is cumbersome due to the lack of orthogonal deposition chemistries, necessitating serial amplification for one target after another (Denkers *et al*., 2004; Kosman *et al*., 2004; Clay & Ramakrishnan, 2005; Barroso-Chinea *et al*., 2007; Tóth & Mezey, 2007; Glass *et al*., 2009; Stack *et al*., 2014; Mitchell *et al*., 2014; Tsujikawa *et al*., 2017), staining is qualitative rather than quantitative, and spatial resolution is routinely compromised by diffusion of reporter molecules prior to deposition (Tautz & Pfeifle, 1989; Takakura *et al*., 1997; Sillitoe & Hawkes, 2002; Thisse *et al*., 2004; Acloque *et al*., 2008; Weiszmann *et al*., 2009). Notably, cumbersome serial multiplexing leads to progressive sample degradation and lengthy protocols. For example, it takes four days for 2-plex imaging in whole-mount zebrafish embryos (Thisse *et al*., 2004; Clay & Ramakrishnan, 2005) and five days for 3-plex imaging in whole-mount chicken embryos (Denkers *et al*., 2004; Acloque *et al*., 2008).

To overcome these longstanding shortcomings of CARD in the context of RNA imaging, in situ amplification based on the mechanism of hybridization chain reaction (HCR) (Dirks & Pierce, 2004) draws on principles from the emerging discipline of dynamic nucleic acid nanotechnology to enable multiplex, quantitative, high-resolution RNA fluorescence in situ hybridization (RNA-FISH) with high signal-to-background in highly autofluorescent samples (Choi *et al*., 2010; Choi *et al*., 2014; Choi *et al*., 2016; Shah *et al*., 2016b; Trivedi *et al*., 2018; Choi *et al*., 2018). Using HCR RNA-FISH, targets are detected using probes that trigger isothermal enzyme-free chain reactions in which fluorophore-labeled HCR hairpins self-assemble into tethered fluorescent amplification polymers, boosting the signal above autofuorescence. The programmability of HCR allows orthogonal amplifiers to operate independently within the sample so that the experimental timeline for multiplex experiments is independent of the number of target RNAs (Choi *et al*., 2010; Choi *et al*., 2014). The amplified HCR signal scales approximately linearly with the number of target molecules, enabling accurate and precise RNA relative quantitation with subcellular resolution in an anatomical context (Trivedi *et al*., 2018; Choi *et al*., 2018). Amplification polymers remain tethered to their initiating probes, preventing the signal from diffusing away from the target, enabling imaging of RNA expression with subcellular or single-molecule resolution as desired (Choi *et al*., 2014; Shah *et al*., 2016b; Choi *et al*., 2016; Choi *et al*., 2018). Building on these advances, HCR immunofluorescence (IF) extends the benefits of HCR signal amplification to protein imaging (Schwarzkopf *et al*., 2021), enabling accurate and precise protein relative quantitation with subcellular resolution in highly autofluorescent samples including formalin-fixed paraffin-embedded (FFPE) tissue sections. Moreover, simultaneous HCR RNA-FISH/IF provides a unified framework for multiplex, quantitative, high-resolution RNA and protein imaging in highly autofluorescent samples with 1-step HCR signal amplification performed for all target RNAs and proteins simultaneously (Schwarzkopf *et al*., 2021).

With conventional bandpass fluorescence imaging, fluorescence is excited with a light source for each channel (e.g., a laser) and collected as a single intensity using a different band-pass filter for each channel. Fluorescence microscopes are typically able to distinguish at most 4 or 5 fluorophores using bandpass imaging due to overlapping spectra for different fluorophores (e.g., see Fig. 1A). Dramatically higher levels of multiplexing can be achieved using temporal barcoding methods (Lubeck *et al*., 2014; Chen *et al*., 2015; Moffitt *et al*., 2016; Shah *et al*., 2016a), but these approaches require low target expression levels in order to spatially separate the signal for each target molecule as a distinct dot, and require repeated imaging that is unfavorable for whole-mount embryos and delicate specimens. Spectral imaging (Garini *et al*., 2006; Mansfield *et al*., 2008; Valm *et al*., 2016) offers a strategy for exceeding 5-plex without the need for repeated staining, stripping, and imaging and without constraints on target expression level or pattern. However, in practice, it has proven difficult to realize the conceptual promise to move beyond 5 targets in the challenging imaging environment of whole-mount vertebrate embryos (Cutrale *et al*., 2017; Cutrale *et al*., 2019).

**Fig. 1:**
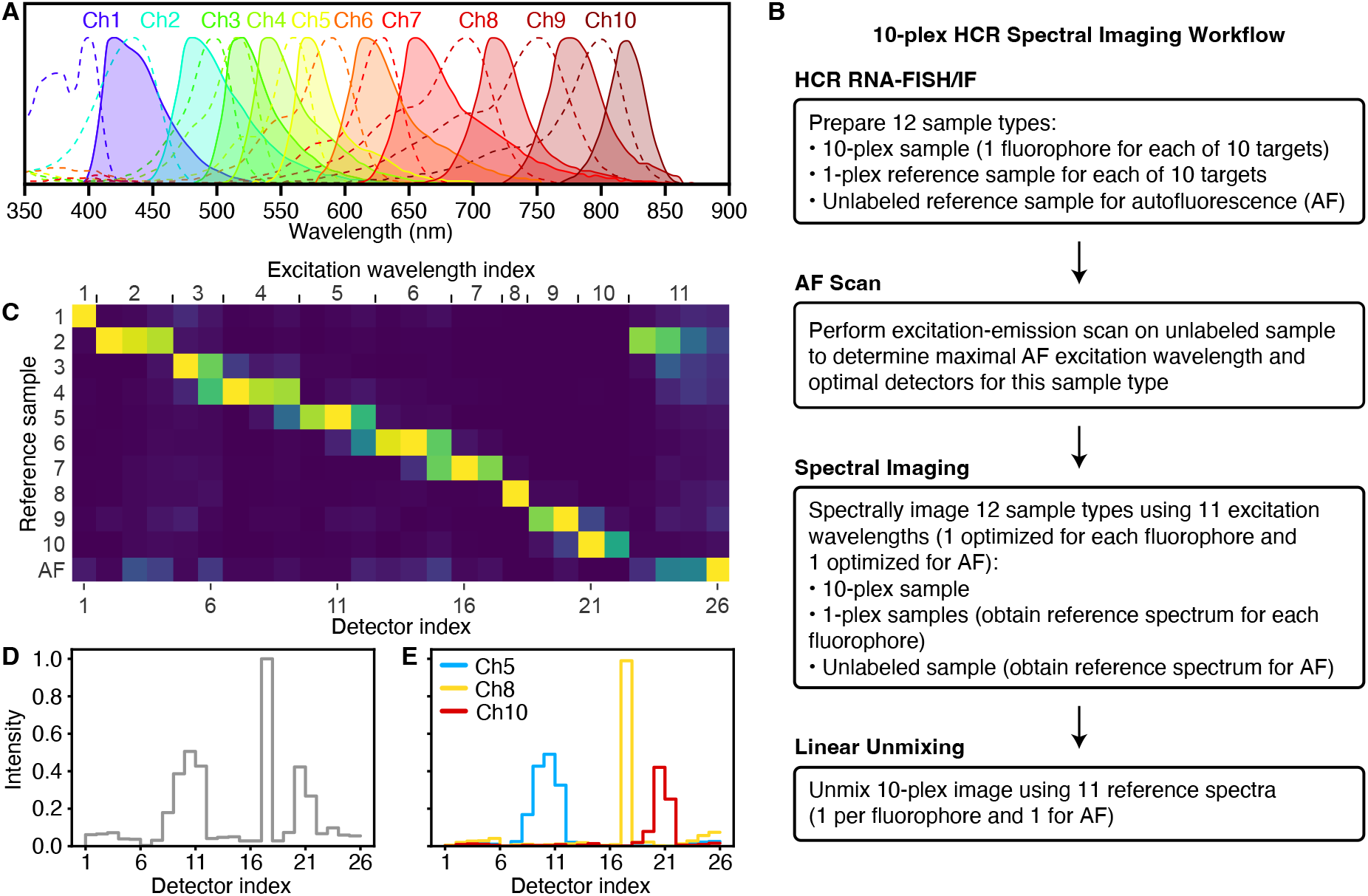
Overview of 10-plex HCR spectral imaging and linear unmixing. (A) Excitation spectra (dashed lines) and emission spectra (solid lines) for the 10 fluorophores selected for HCR spectral imaging (left to right: Alexa405, Atto425, Alexa488, Alexa514, Alexa546, Alexa594, Atto633, Alexa700, Alexa750, iFluor800; spectra from FluoroFinder). (B) Workflow for performing a 10-plex HCR spectral imaging and linear unmixing experiment. (C) Linear unmixing matrix of normalized reference spectrum intensities gathered via spectral imaging of 10 fluorophores in 1-plex samples and autofluorescence (AF) in an unlabeled sample for 27 hpf zebrafish embryos. (D) Spectral fluorescence intensities for a single pixel in the notochord of a 27 hpf zebrafish embryo. (E) Linear unmixing of the pixel fluorescence of panel D using the reference spectra from panel C to determine the fluorescence contribution of each fluorophore and AF, revealing that the fluorescence in this notochord pixel is predominantly a combination of Ch5, Ch8, and Ch10 fluorophores.

With spectral imaging, each fluorophore is excited by one or more lasers and fluorescence is collected with one or more detectors, generating a signal for each pixel that is a superposition of fluorescence from multiple fluorophores, characterized for multiple excitation wavelengths using multiple detectors at different emissions wavelengths (Garini *et al*., 2006). Spectrally imaging a set of reference samples that each contain only one type of fluorophore provides a reference spectrum for each channel. Each pixel in a spectral image of the sample of interest may then be interpreted as a linear superposition of these reference spectra with an unknown coefficient for each channel. Linear unmixing of the fluorescence in that pixel then determines the non-negative coefficients that best approximate the spectrum of the sample of interest, revealing the fluorescence contribution of each channel (Mansfield *et al*., 2008). The output of linear unmixing is a set of unmixed fluorescence channels for all pixels. The mathematics of spectral imaging are enticing, hypothetically enabling separation of an arbitrary number of fluorophore channels, but the imperfections and complexities of real-world samples and experiments have so far undermined the practicality of spectral imaging in whole-mount vertebrate embryos. Here, we establish standardized ingredients and workflows for HCR spectral imaging to enable biologists to perform robust 10-plex imaging of RNA and protein targets in whole-mount vertebrate embryos and other highly autofluorescent samples in a plug-and-play manner. Moreover, we demonstrate that HCR spectral imaging and linear unmixing generate high signal-to-background, quantitative signal, and high resolution in all 10 channels.

## RESULTS

### Ingredients and workflow for robust 10-plex HCR spectral imaging

Using 10-plex HCR spectral imaging, each of 10 targets is detected using a different probe set comprising one or more probes capable of triggering a different HCR amplifier carrying a different fluorophore. Our approach combines several ingredients into a standardized workflow. First, an optimized set of 10 orthogonal HCR amplifiers (one per target) that operate independently in the same sample at the same time. The role of these amplifiers is to boost the signal strength relative to autofluorescence in each channel, increasing the ease and robustness of linear unmixing, and ultimately the signal-to-background of the unmixed channels. Second, an optimized set of 10 fluorophores (one per HCR amplifier) with overlapping excitation and emissions spectra selected to facilitate subsequent linear unmixing (spectra depicted in Fig. 1A). Third, an optimized set of excitation wavelengths (one per fluorophore; see Table S5) and an optimized set of detection wavelengths (using between 1 and 3 detectors per fluorophore; see Table S5). Fourth, an optimized excitation wavelength and set of detection wavelengths for autofluorescence, which is treated as an 11th channel; the optimized excitation and emission values for autofluorescence will need to be determined for a given sample type by the user in a preliminary experiment as described below. Fifth, spectral imaging hardware that provides the flexibility to use optimal excitation wavelengths for each fluorophore and for autofluorescence; we use the Leica Stellaris 8 confocal microscope, which is equipped with a tunable white light laser capable of generating laser lines in 1 nm increments between 440 nm to 790 nm (in addition to a fixed 405 nm laser). Sixth, a linear unmixing algorithm that takes as input a 10-plex spectral image and 11 reference spectra (one per fluorophore and one for autofluorescence) and returns as output 11 unmixed channels (one per fluorophore and one for autofluorescence); we use either the Leica LAS X software or our own Unmix 1.0 software package.

These ingredients are employed in a standardized workflow (see Fig. 1B) to perform robust 10-plex HCR spectral imaging of 10 RNA and/or protein targets. First, the user prepares 12 sample types: a 10-plex sample (or multiple replicate samples as desired), a 1-plex reference sample for each of 10 targets, and two unlabeled autofluorescence (AF) samples (one for an excitation-emission scan and one AF reference sample). Second, the user employs one AF sample to perform an excitation-emission scan to determine the excitation wavelength that maximizes autofluorescence, which in turn determines a set of optimized detection wavelengths (using 4 detectors); this preliminary step need not be repeated for future experiments in the same sample type. Third, the user spectrally images the 10-plex sample using the standardized set of 11 excitation wavelengths (1 optimized for each fluorophore and 1 optimized for AF). Likewise, the user spectrally images each 1-plex reference sample using the 11 standardized excitation wavelengths to obtain a reference spectrum for each fluorophore in a region of maximum expression for a given target. Further, the user spectrally images the AF reference sample using the 11 standardized excitation wavelengths to obtain a reference spectrum for AF in a region of maximum autofluorescence. Fourth, the 11 reference spectra (one per fluorophore and one for AF) are used to linearly unmix the 10-plex image and produce 11 unmixed channels (one per fluorophore and one for AF).

As an example, Fig. 1C depicts 11 reference spectra (one per fluorophore and one for autofluorescence) collected in a whole-mount zebrafish embryo. Each reference sample was excited with 11 different excitation wavelengths (standardized ingredients of the method) and for each excitation wavelength, emissions were collected with between 1 and 4 detectors (a total of 26 detectors spanning the range of emissions wavelengths; also standardized ingredients of the method). From a spectral image of the 10-plex sample, Fig. 1D depicts the raw spectral fluorescence intensities for a single pixel in the notochord region of a whole-mount zebrafish embryo collected using 11 excitation wavelengths and a total of 26 detectors. Linear unmixing determines the (non-negative) contributions from each of the 10 fluorophores and AF (treated as an 11th channel) that combine to create the spectral fluorescence intensity curve. Fig. 1E displays that the fluorescence in this notochord pixel is predominantly a combination of Ch5, Ch8, and Ch10 fluorophores that together generate the three peaks seen in the spectral fluorescence intensity curve of Fig. 1D.

### 10-plex RNA imaging using spectral HCR RNA-FISH in a whole-mount vertebrate embryo

We first validated 10-plex HCR spectral imaging for HCR RNA-FISH, which is performed using the two-stage protocol summarized in Fig. 2A. During the detection stage, an RNA target is detected using a probe set comprising one or more pairs of split-initiator DNA probes, each carrying a fraction of HCR initiator i1 (Choi *et al*., 2018). Probe pairs that hybridize specifically to proximal binding sites on the target RNA colocalize a full HCR initiator i1 capable of triggering HCR signal amplification. Meanwhile, any individual probes that bind nonspecifically in the sample do not colocalize full HCR initiator i1 and do not trigger HCR. During the amplification stage, each colocalized full HCR initiator i1 triggers self-assembly of metastable fluorophore-labeled HCR hairpins (h1 and h2) into a tethered fluorescent HCR amplification polymer to generate an amplified signal at the site of the target RNA. For 10-plex HCR spectral imaging, the same 2-stage protocol is used as for a 1-plex experiment, with all 10 targets detected in parallel during the detection stage, and signal amplification performed for all 10 targets in parallel during the amplification stage (Fig. 2B). To validate 10-plex HCR spectral imaging, we performed HCR RNA-FISH in whole-mount zebrafish embryos for 10 target mRNAs with known expression patterns and a range of expression levels (Fig. 2C). Following the workflow of Fig. 1B, we prepared samples for 10-plex zebrafish replicates, as well as 10 1-plex reference samples, and two AF samples. One AF sample was used to determine the optimal excitation wavelength and detector wavelengths for autofluorescence. We spectrally imaged three replicate 10-plex samples (using probes and amplifiers for all 10 targets) as well as 11 reference samples (a 1-plex sample for each of 10 targets using a probe set and amplifier for only that target and an AF reference sample using no probes and amplifiers). Linear unmixing returned 11 un-mixed channels (Fig. 2C; one channel per target plus an 11th AF channel). Visual inspection of the expression patterns reveals robust separation of all 10 targets, including those with overlapping expression patterns (e.g., *shha, ntla*, and *col2a1a* in the notochord (Moreno-Ayala *et al*., 2015)). High signal-to-background is achieved for all 10 targets using 10 fluorophores with overlapping spectra that span the visible and near-IR spectrum, including wavelengths with high autofluorescence. We estimate signal-to-background for each channel by characterizing signal plus background in a region of high expression and background in a region of no or low expression. This approach yields a conservative estimate of performance, as characterizing background in a region of little or no expression places an upper bound on background and hence a lower bound on signal-to-background. Across the 10 target mRNAs, the estimated signal-to-background ratio for each target ranged from 17 to 100 with a median of 46.5 (see Table S7 for additional details).

**Fig. 2:**
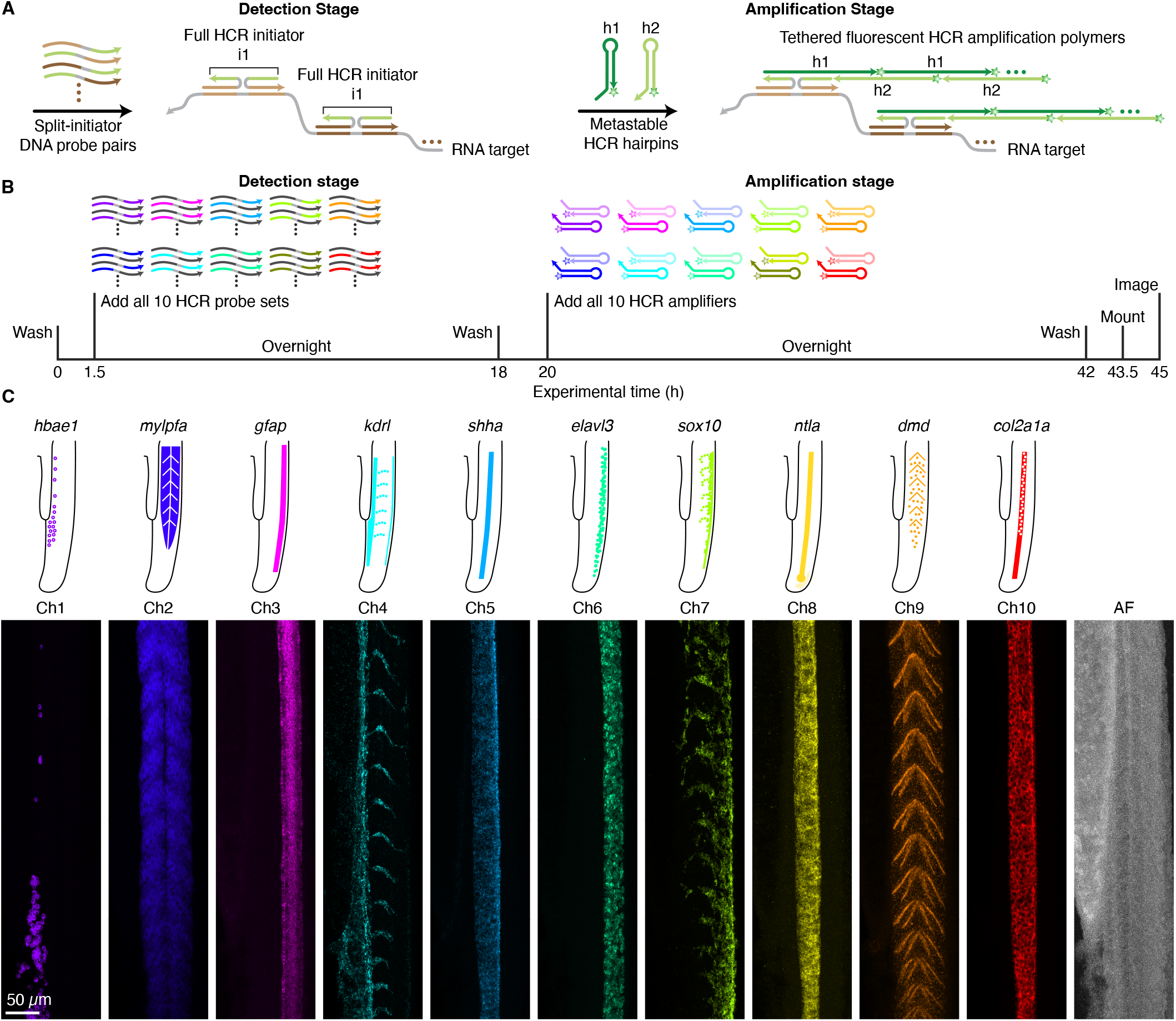
10-plex RNA imaging in a whole-mount zebrafish embryo. (A) 2-stage HCR RNA-FISH protocol. Detection stage: split-initiator DNA probe pairs bind to RNA targets to colocalize full HCR initiator i1; wash. Amplification stage: colocalized full initiator i1 triggers self-assembly of fluorophore-labeled HCR hairpins into tethered fluorescent amplification polymers; wash. (B) 10-plex HCR RNA-FISH timeline. During the detection stage, all 10 targets are detected in parallel. During the amplification stage, signal amplification is performed for all 10 targets in parallel. (C) Top: expression atlas for 10 RNA targets in the tail of a zebrafish embryo (lateral view). Bottom: Linearly unmixed channels from a 10-plex confocal image including autofluorescence (AF) as an 11th channel; maximum intensity z-projection; 0.57*×*0.57*×*4.0 μm pixels. Embryo fixed 27 hpf. See Section S3.1 and Supplementary Movie 1 for additional data.

### 10-plex qHCR imaging: mRNA relative quantitation with subcellular resolution in an anatomical context

We have previously demonstrated that HCR RNA-FISH and HCR IF enable accurate and precise relative quantitation of RNA and protein targets with subcellular resolution in an anatomical context (qHCR imaging mode), generating an amplified signal that scales approximately linearly with the number of target molecules per imaging voxel (Trivedi *et al*., 2018; Choi *et al*., 2018; Schwarzkopf *et al*., 2021). Here, we validate that spectral imaging and linear unmixing preserve the quantitative nature of HCR imaging. To test relative quantitation for all 10 channels in a whole-mount zebrafish embryo, we redundantly detected each of five target mRNAs with two probe sets that each trigger a different HCR amplifier carrying a different fluorophore (Fig. 3A), leading to a 10-channel image with two channels for each of the 5 target mRNAs (Fig. 3B). If HCR signal scales approximately linearly with the number of target mRNAs per voxel, a two-channel scatter plot of normalized voxel intensities for each target will yield a tight linear distribution with zero intercept (Trivedi *et al*., 2018). Consistent with expectation, we observe high accuracy (linearity with zero intercept) and precision (scatter around the line) for subcellular voxels for all five target mRNAs (Fig. 3C) and all 10 channels. Moreover, all 10 channels provide high signal-to-background ranging from 27 to 130 with a median of 60 (see Table S10 for additional details).

**Fig. 3:**
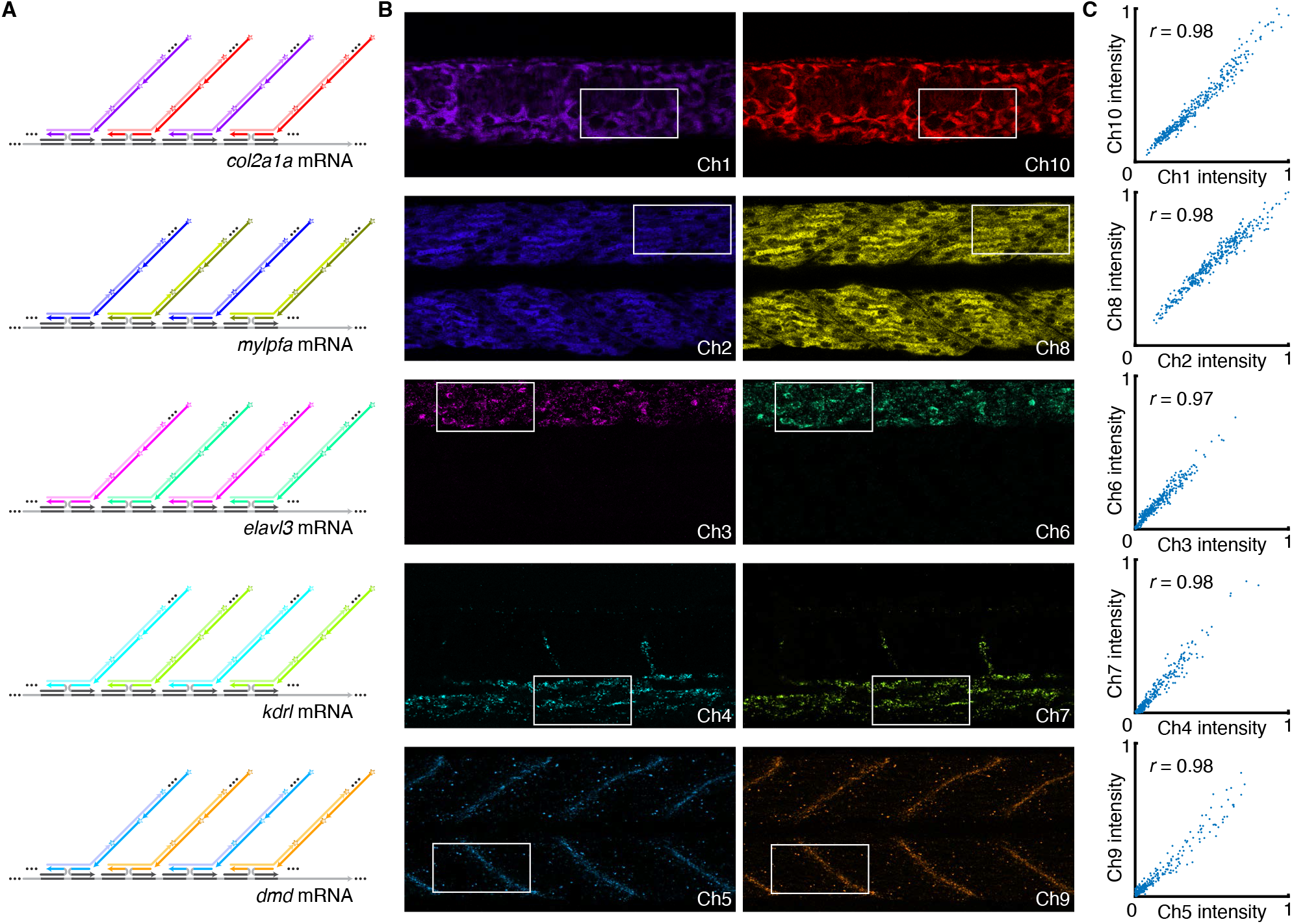
10-plex qHCR imaging: mRNA relative quantitation with subcellular resolution in an anatomical context using spectral imaging with linear unmixing. (A) Simultaneous 2-channel redundant detection of each of 5 target mRNAs using a total of 10 channels in a whole-mount zebrafish embryo. Each target mRNA is detected using two split-initiator DNA probe sets, each initiating an orthogonal HCR amplifier labeled with a different fluorophore. (B) Linearly unmixed channels from a 10-plex confocal image; single optical sections; 0.180*×*0.180*×*1.2 μm pixels. Embryo fixed 27 hpf. (C) High accuracy (linearity with zero intercept) and precision (scatter around the line) for mRNA relative quantitation in an anatomical context. Highly correlated normalized signal (Pearson correlation coefficient, *r*) for subcellular voxels (2.0*×*2.0*×*1.2 μm) in the depicted region of panel B. See Section S3.2 for additional data.

### dHCR imaging: digital mRNA absolute quantitation in a 10-plex sample

We have previously demonstrated that HCR enables digital mRNA absolute quantitation with single-molecule resolution for low-expression targets imaged at high magnification (dHCR imaging mode), enabling counting of individual target molecules as dots in highly autofluorescent samples including whole-mount vertebrate embryos and thick brain slices (Shah *et al*., 2016b; Choi *et al*., 2018). We were curious whether spectral imaging and linear unmixing would preserve the single-molecule imaging capabilities of HCR imaging. To examine this question, we spectrally imaged the 10-plex redundant detection samples of Fig. 3 again at high magnification in the dorsal posterior region of the zebrafish tail, where the target mRNA *kdrl* is expressed as single-molecule punctae. In those samples, *kdrl* is redundantly detected in conjunction with redundant detection of four other target mRNAs.

To determine the fidelity with which single-molecule targets can be detected with spectral HCR imaging, we performed dot detection on the redundant *kdrl* channels (Ch4 and Ch7) using dot detection methods drawn from the computer vision community (Fig. 4). As false-positive and false-negative rates for each channel go to zero, the colocalization fraction for each channel (fraction of dots in a given channel that are in both channels) will approach one from below. We observe colocalization rates of *≈*0.8 for both channels, roughly comparable to the performance observed in 2-plex experiments using band-pass HCR imaging (Choi *et al*., 2018), indicating that spectral imaging and linear unmixing are compatible with single-molecule imaging in 10-plex experiments.

**Fig. 4:**
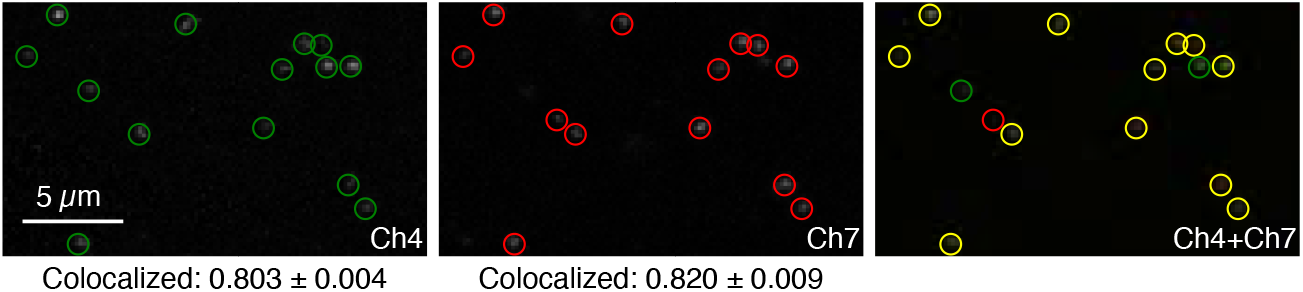
dHCR imaging: digital mRNA absolute quantitation in an anatomical context using 10-plex spectral imaging with linear unmixing. 2-channel redundant detection of target mRNA *kdrl* alongside 2-channel redundant detection of 4 other target mRNAs as part of a 10-plex experiment in a whole-mount zebrafish embryo (see Fig. 3A). Linearly unmixed channels from a 10-plex confocal image in the dorsal posterior tail, where *kdrl* is expressed as single-molecule punctae. Representative field of view; maximum intensity z-projection; 0.18*×*0.18 *×*1.2 μm pixels. Left: Ch4 (*kdrl*). Middle: Ch7 (*kdrl*). Right: Ch4+Ch7 merge. Green circles: dots detected in Ch4. Red circles: dots detected in Ch7. Yellow circles: dots detected in both channels. Colocalization represents the fraction of dots in one channel that are detected in both channels (mean*±*SEM, *N* = 3 replicate embryos). Embryo fixed 27 hpf. See Section S3.3 for additional data.

### 10-plex RNA and protein imaging using spectral HCR RNA-FISH/IF in a mouse brain section

We have previously shown that HCR RNA-FISH/IF enables simultaneous multiplex, quantitative, high-resolution imaging of RNA and protein targets in highly autofluorescent samples (Schwarzkopf *et al*., 2021). To demonstrate the versatility of 10-plex HCR spectral imaging, here we image a total of 7 mRNA targets and 3 protein targets in a coronal mouse brain section. HCR RNA-FISH/IF is performed using the three-stage protocol summarized in Fig. 5A. During the protein detection stage, protein targets are detected with unlabeled primary antibody probes that are subsequently detected by initiator-labeled secondary antibody probes. During the RNA detection stage, RNA targets are detected using split-initiator DNA probe sets. During the amplification stage, 1-step HCR signal amplification is performed for all 10 RNA and protein targets simultaneously. The same three-stage protocol is used to image any combination of up to 10 RNA and protein targets simultaneously (Fig. 5B).

**Fig. 5:**
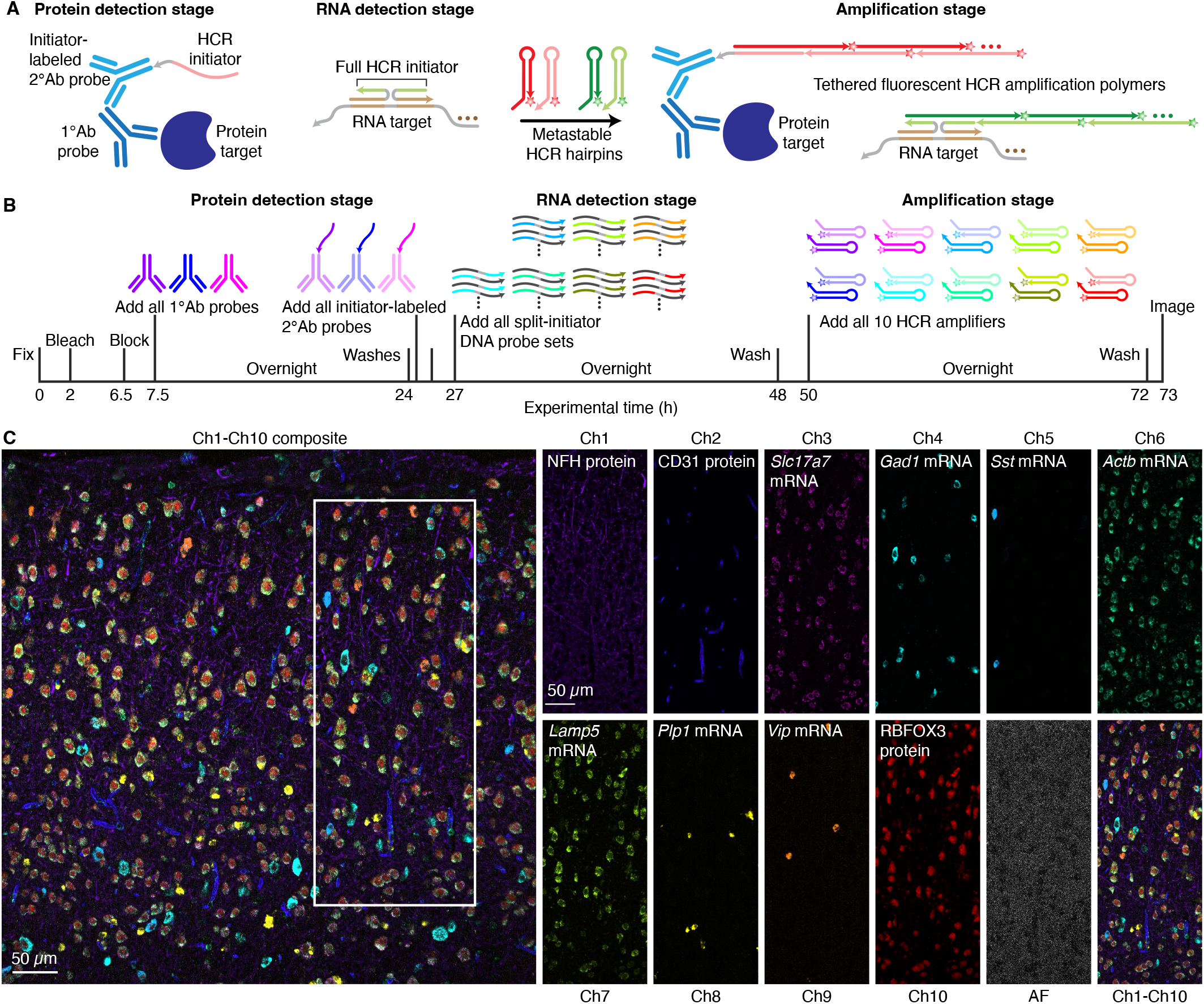
10-plex RNA and protein imaging in a fresh-frozen mouse brain section. (A) 3-stage HCR RNA-FISH/IF protocol. Protein detection stage: unlabeled primary antibody probes bind to protein targets; wash; initiator-labeled secondary antibody probes bind to primary antibody probes; wash. RNA detection stage: split-initiator DNA probes bind to RNA targets; wash. Amplification stage: initiators trigger self-assembly of fluorophore-labeled HCR hairpins into tethered fluorescent amplification polymers; wash. (B) 10-plex HCR RNA-FISH/IF timeline. During the protein detection stage, all protein targets are detected in parallel. During the RNA detection stage, all RNA targets are detected in parallel. During the amplification stage, signal amplification is performed for all 10 targets in parallel. (C) 10-plex confocal image of 3 protein targets and 7 mRNA targets in the cerebral cortex of a fresh-frozen mouse brain section; single optical section; 0.57*×*0.57*×*4.0 μm pixels. Left: composite image of linearly unmixed Ch1-Ch10. Right: single fluorescence channels for the depicted region in the composite image (including autofluorescence (AF) as an 11th channel). Sample: fresh-frozen mouse brain section (coronal); 5 μm thickness; interaural region 0.88 mm*±* 0.20 mm; 8 weeks old. See Section S3.4 for additional data.

We used the workflow of Fig. 1B to perform HCR spectral imaging for a 10-plex sample as well as for 11 reference samples (1 per fluorophore and 1 for autofluorescence). Linear unmixing returned 11 unmixed channels (Fig. 5C), one for each of 7 target mRNAs, one for each of 3 target proteins, and one for autofluorescence. Each target RNA and protein has a characteristic expression pattern corresponding to certain cell types and/or cellular compartments within the cerebral cortex, enabling verification that fluorophores were successfully separated for all 10 targets. For the protein targets, NFH labels the intermediate filaments in large myelinated axons (Giasson & Mushynski, 1996), CD31 labels endothelial cells (Zhang *et al*., 2021), and RBFOX3 labels neuronal nuclei (Lucas *et al*., 2014). Among the mRNA targets, *Actb* is expressed in several cell types (Wang *et al*., 2020), while *Slc17a7* labels excitatory neurons and *Gad1* labels inhibitory neurons (Tasic *et al*., 2018). Subtypes of inhibitory neurons are labeled by *Sst* and *Vip*, and expression of these two mRNA targets overlaps as expected with *Gad1* -expressing neurons (Tasic *et al*., 2018); these results further confirm the ability to distinguish targets with overlapping expression patterns using HCR spectral imaging with linear unmixing. Lastly, as expected, *Lamp5* expression is most pronounced in the upper layers of the cortex (Tiveron *et al*., 2016), and *Plp1* labels oligodendrocytes, which are non-overlapping with the neuronal cells (Zeisel *et al*., 2015). Across 10 RNA and protein targets, high signal-to-background was achieved for all targets, ranging from 25 to 140 with a median of 67.5 (see Table S12 for additional details).

## DISCUSSION

HCR spectral imaging with linear unmixing enables researchers to simultaneously image any combination of 10 RNA and protein targets without placing limitations on expression level (high/low) or pattern (overlapping/non-overlapping). The approach is well-suited for whole-mount vertebrate embryos and delicate specimens because the method does not employ repeated staining, stripping, and imaging to achieve multiplexing, so iterative sample degradation is avoided and there is no requirement for the sample to be immobilized on a slide during staining. 10-plex HCR RNA-FISH (RNA targets only) or HCR IF (protein targets only) is achieved using a two-stage protocol involving two overnight incubations (detection stage, amplification stage). 10-plex HCR RNA-FISH/IF (combination of RNA and protein targets) is achieved using a three-stage protocol involving three overnight incubations (protein detection stage, RNA detection stage, amplification stage). All protocols are compatible with a normal sleep schedule.

Spectral imaging and linear unmixing preserve two complementary quantitative HCR imaging modes in the context of 10-plex experiments: 1) qHCR imaging – analog RNA or protein relative quantitation with subcellular resolution in an anatomical context, 2) dHCR imaging – digital mRNA absolute quantitation with single-molecule resolution in an anatomical context. Multiplex HCR imaging enables bi-directional quantitative discovery (Trivedi *et al*., 2018): *read-out* from anatomical space to expression space to discover co-expression relationships in selected regions of the sample; *read-in* from expression space to anatomical space to discover those anatomical locations in which selected gene co-expression relationships occur. Here, by using HCR spectral imaging and linear unmixing to achieve 10-plex, high-accuracy, high-precision, high-resolution qHCR imaging, we enable 10-dimensional quantitative read-out/read-in analyses for any combination of 10 RNA and/or protein circuit elements in the anatomical context of highly autofluorescent samples. Note that in performing these quantitative co-expression analyses, a 4-plex experiment enables exploration of 6 target pairs and a 5-plex experiment enables exploration of 10 target pairs, but a 10-plex experiment enables exploration of 45 target pairs, all in the anatomical context of a single sample.

To detect protein targets, the present work performs HCR IF using unlabeled primary antibody probes and initiator-labeled secondary antibody probes, thereby requiring that each primary antibody be of a different isotype or raised in a different host species, which is sometimes feasible given the large libraries of commercially available antibodies. For example, we demonstrate imaging of three protein targets simultaneously using unlabeled chicken, rat, and rabbit primary antibody probes (Fig. 5). However, in situations where it is desirable to use multiple primary antibodies raised in the same host species or of the same isotype, HCR IF can instead be performed using initiator-labeled primary antibody probes (Schwarzkopf *et al*., 2021). Antibody-oligo conjugation can sometimes interfere with target recognition, necessitating post-conjugation probe validation. By contrast, using a small library of validated initiator-labeled secondary antibodies as we do here, primary antibodies can be exploited without modification, eliminating the need to validate antibody-oligo conjugation for each new target protein.

For RNA targets, HCR imaging provides automatic background suppression throughout the protocol, ensuring that reagents will not generate amplified background even if they bind non-specifically in the sample (Choi *et al*., 2018). During the detection stage, split-initiator DNA probes that bind non-specifically in the sample do not colocalize a full HCR initiator and do not trigger HCR. During the amplification stage, metastable HCR hairpins that bind non-specifically do not trigger formation of an amplification polymer. For protein targets, HCR hairpins continue to provide automatic background suppression during the amplification stage, but the detection stage employs antibody probes that carry a full HCR initiator so it is important to use antibodies that are highly selective for their targets and to wash unused antibody probes from the sample. High signal-to-background is achieved across all 10 channels for both RNA and protein targets.

To enable plug-and-play 10-plex HCR spectral imaging, we employ a standardized set of ingredients: 10 orthogonal HCR amplifiers (one per target), an optimized set of 10 fluorophores (one per amplifier), 10 optimized excitation wavelengths (one per fluorophore), 10 sets of optimized detection wavelengths (one to three detectors per fluorophore), an optimized excitation wavelength and set of four detection wavelengths for autofluorescence (measured by the user for the sample type of interest; treated as an 11th channel), spectral imaging hardware that provides the flexibility to use optimal excitation wavelengths for each fluorophore and for autofluorescence (Leica Stellaris 8 confocal microscope), and linear unmixing software (Leica LAS X or Unmix 1.0) that returns 11 unmixed channels (one per fluorophore and one for autofluorescence). For users that do not have access to a Leica Stellaris 8 microscope, our robust ingredients and workflow provide a starting point for adaptation to locally available hardware and our Unmix 1.0 software provides the flexibility to unmix spectral imaging data without use of Leica LAS X software. If a user encounters a sample where different tissues produce widely differing autofluorescence spectra, the number of auxiliary autofluorescence channels can be increased, collecting one AF reference spectrum per AF channel.

These standardized ingredients are combined in a straight-forward workflow. To perform 10-plex HCR spectral imaging in a highly autofluorescent sample, a user need only image a reference spectrum for each fluorophore in a 1-plex sample and an autofluorescence reference spectrum in an unlabeled sample in order to linearly unmix the spectral fluorescence of a 10-plex sample.

## MATERIALS AND METHODS

### Probes, amplifiers, and buffers

Details regarding the probes, amplifiers, and buffers for each experiment are displayed in Table S1.1 for HCR RNA-FISH and Table S1.2 for HCR IF.

### 10-plex HCR spectral imaging and linear unmixing

10-plex HCR spectral imaging and linear unmixing were performed using the protocol detailed in Section S2.1. Linear unmixing was performed using Leica LAS X software for main text figures. Alternatively, linear unmixing can be performed with our Unmix 1.0 software package (as demonstrated in Figs S2 and S20–S22).

### HCR RNA-FISH in whole-mount zebrafish embryos

Within the 10-plex workflow, HCR RNA-FISH in whole-mount zebrafish embryos was performed using the protocols detailed in Section S2.2. Experiments were performed in AB wild-type whole-mount zebrafish embryos (fixed 27 hpf) obtained from the Zebrafish Facility within the Beckman Institute at Caltech. Procedures for the care and use of zebrafish embryos were approved by the Caltech IACUC.

### HCR RNA-FISH/IF in fresh-frozen mouse brain sections

Within the 10-plex workflow, HCR RNA-FISH/IF in fresh-frozen mouse brain sections was performed using the protocols detailed in Section S2.3. Experiments were performed in C57BL/6 fresh-frozen coronal mouse brain sections (thickness: 5 μm; region: interaural 0.88 mm *±* 0.2 mm; age: 8 weeks old; sex: male) from Acepix Biosciences (Cat. # A2203-0561).

### Confocal microscopy

Microscopy was performed using a Leica Stellaris 8 inverted confocal microscope with an HC PL APO 20*×*/0.75 IMM CORR CS2 (Cat. # 11506343) objective or HC PL APO 63*×*/1.40 OIL CS2 (Cat. # 11506350) objective. Details on the objectives, excitation wavelengths, detectors, and detection wavelengths used for each experiment is displayed in Table S3.

### Image analysis

Image analysis was performed as detailed in Section S1.4, including: definition of raw pixel intensities; measurement of signal, background, and signal-to-background; calculation of normalized subcellular voxel intensities for qHCR imaging; and dot detection and colocalization for dHCR imaging. Dot detection and colocalization were performed with our Dot Detection 2.0 software package. For qHCR redundant detection experiments (Fig. 3; see Section S3.2), Huygens Software (Scientific Volume Imaging) was used to correct for chromatic aberration across the 10 channels. All images are displayed with 0.1% of pixels saturated across three replicates, with the exception of the single-molecule images of Figs 4 and S15, which are displayed with no saturated pixels. All images are displayed without background subtraction.

## Acknowledgments

We thank A. Kahan of the V. Gradinaru Lab for the gift of mouse brain sections during initial validation, M. E. Bronner for reading a draft of the manuscript, and the following resources within the Beckman Institute at Caltech: G. Shin of Molecular Technologies for providing HCR reagents, G. Spigolon of the Biological Imaging Facility for assistance with imaging, and the Zebrafish Facility for providing zebrafish embryos.

## Competing interests

The authors declare competing financial interests in the form of patents, pending patent applications, and the startup company Molecular Instruments, Inc.

## Author contributions

Conceptualization: N.A.P.; Methodology: S.J.S., M.E.F., N.A.P.; Software: M.E.F; Validation: S.J.S., M.E.F., J.H.; Investigation: S.J.S., J.H.; Writing -original draft: S.J.S., N.A.P.; Writing -review & editing: all; Visualization: S.J.S. and M.E.F.; Supervision: N.A.P.; Project administration: N.A.P.; Funding acquisition: N.A.P.

## Funding

This work was funded by the National Institutes of Health (NIBIB R01EB006192 and NIGMS training grant GM008042 to S.J.S.) and by the Beckman Institute at Caltech (PMTC). The Leica Stellaris 8 confocal microscope in the Biological Imaging Facility within the Beckman Institute at Caltech was purchased with support from Caltech and the following Caltech entities: the Beckman Institute, the Resnick Sustainability Institute, the Division of Biology & Biological Engineering, and the Merkin Institute for Translational Research.

## Supplementary information

Supplementary information available online.

