## Supplementary Information for "HCR spectral imaging: 10-plex, quantitative, high-resolution RNA and protein imaging in highly autofluorescent samples"

**S3 Additional studies** **27**

**References** **66**

### S1 Additional materials and methods

#### S1.1 Probe and amplifier details for RNA targets using HCR RNA-FISH

| Species | Sample | RNA target | Split-initiator probe pairs | Probe set catalog # | HCR amplifier catalog # | Figures |
|---|---|---|---|---|---|---|
| *D. rerio* | whole-mount embryo | *hbae1* | 3 | 4503/E784 | B1-Alexa405 | 2C, S2–S3 |
| | | *mylpfa* | 13 | 4644/E992 | B2-Atto425 | 2C, S2–S3 |
| | | *gfap* | 17 | 3974/E108 | B6-Alexa488 | 2C, S2–S3 |
| | | *kdrl* | 60 | 4503/E786 | B9-Alexa514 | 2C, S2–S3 |
| | | *shha* | 20 | 3699/D741 | B7-Alexa546 | 2C, S2–S3 |
| | | *elavl3* | 20 | 4362/E588 | B10-Alexa594 | 2C, S2–S3 |
| | | *sox10* | 34 | 4644/E990 | B8-Atto633 | 2C, S2–S3 |
| | | *ntla* | 20 | 2196/A430 | B3-Alexa700 | 2C, S2–S3 |
| | | *dmd* | 20 | 4607/E930 | B5-Alexa750 | 2C, S2–S3 |
| | | *col2a1a* | 15 | 4560/E856 | B4-iFluor800 | 2C, S2–S3 |
| *D. rerio* | whole-mount embryo | *col2a1a* | 10 | 4607/E918 | B1-Alexa405 | 3B, S6, S11, S16 |
| | | *col2a1a* | 10 | 4607/E920 | B4-iFluor800 | 3B, S6, S11, S16 |
| | | *mylpfa* | 6 | 4607/E924 | B2-Atto425 | 3B, S7, S12, S16 |
| | | *mylpfa* | 6 | 4607/E922 | B3-Alexa700 | 3B, S7, S12, S16 |
| | | *elavl3* | 20 | 4607/E908 | B6-Alexa488 | 3B, S8, S13, S16 |
| | | *elavl3* | 20 | 4607/E910 | B10-Alexa594 | 3B, S8, S13, S16 |
| | | *kdrl* | 30 | 3665/D679 | B9-Alexa514 | 3B, S9, S14, S16 |
| | | *kdrl* | 30 | 4644/E994 | B8-Atto633 | 3B, S9, S14, S16 |
| | | *dmd* | 20 | 4607/E930 | B5-Alexa750 | 3B, S10, S15, S16 |
| | | *dmd* | 20 | 4607/E932 | B7-Alexa546 | 3B, S10, S15, S16 |
| *D. rerio* | whole-mount embryo | *kdrl* | 30 | 3665/D679 | B9-Alexa514 | 4, S17 |
| | | *kdrl* | 30 | 4644/E994 | B8-Atto633 | 4, S17 |
| *M. musculus* | brain section | *Slc17a7* | 20 | 4671/F037-P | B6-Alexa488 | 5C, S20–S23 |
| | | *Gad1* | 33 | 4631/E968-P | B9-Alexa514 | 5C, S20–S23 |
| | | *Sst* | 11 | 4797/F111-P | B7-Alexa546 | 5C, S20–S23 |
| | | *Actb* | 20 | 2306/A758-P | B5-Alexa594 | 5C, S20–S23 |
| | | *Lamp5* | 27 | 4761/F033-P | B8-Atto633 | 5C, S20–S23 |
| | | *Plp1* | 24 | 4787/F081-P | B3-Alexa700 | 5C, S20–S23 |
| | | *Vip* | 24 | 4611/E972-P | B10-Alexa750 | 5C, S20–S23 |

**Table S1. Organism, sample type, target RNA, probe set details, HCR amplifier details, and figure numbers for HCR RNA-FISH.** For HCR RNA-FISH, HCR probe sets, amplifiers, and buffers (probe hybridization buffer, probe wash buffer, amplification buffer) were obtained from Molecular Technologies (MT) within the Beckman Institute at Caltech.

#### S1.2 Probe and amplifier details for protein targets using HCR IF

| Species | Sample | Protein target | 1°Ab probe (unlabeled)<br>2°Ab probe (initiator-labeled) | Dilution factor<br>working concentration ($\mu$g/mL) | Probe supplier (catalog #) | HCR amplifier supplier (catalog #) | Figures |
|---|---|---|---|---|---|---|---|
| *M. musculus* | brain section | NFH | 1°pAb chicken IgY anti-NFH | 1:2,000 | Inv (PA1-10002) | | |
| | | | 2°pAb donkey anti-chicken IgY-B1 | 1 | MI | MT (B1-Alexa405) | 5C, S20–S23 |
| | | CD31 | 1°mAb rat IgG anti-CD31 | 1:10 | BDB (550274) | | |
| | | | 2°pAb donkey anti-rat IgG-B2 | 1 | MI | MT (B2-Atto425) | 5C, S20–S23 |
| | | RBFOX3 | 1°mAb rabbit IgG anti-RBFOX3 | 1:5,000 | CST (24307) | | |
| | | | 2°pAb donkey anti-rabbit IgG-B4 | 1 | MI | MT (B4-iFluor800) | 5C, S20–S23 |

**Table S2. Organism, sample type, target protein, 1°Ab probe details, 2°Ab probe details, HCR amplifier details, and figure numbers for HCR IF.** For HCR IF, initiator-labeled secondary antibody probes and antibody buffer were obtained from Molecular Instruments (MI) and amplifiers and amplification buffer were obtained from Molecular Technologies (MT) within the Beckman Institute at Caltech. Inv: Invitrogen. BDB: BD Biosciences. CST: Cell Signaling Technology.

##### S1.3 Confocal microscope settings

| Sample | Target | Objective | Fluorophore | Laser (nm) | Detector(s) | Detection wavelengths (nm) | Pixel size $(x \times y \times z\ \mu\text{m})$ | Figures |
|---|---|---|---|---|---|---|---|---|
| Whole-mount embryo | *hbae1* | 20× | Alexa405 | 405 | HyD S 1 | 410–450 | 0.568 × 0.568 × 4.0 | 2C, S2–S3 |
| | *mylpfa* | 20× | Atto425 | 440 | HyD S 1, S 2, S 3 | 450–475, 475–495, 495–520 | 0.568 × 0.568 × 4.0 | 2C, S2–S3 |
| | *gfap* | 20× | Alexa488 | 488 | HyD S 1, S 2 | 493–513, 513–533 | 0.568 × 0.568 × 4.0 | 2C, S2–S3 |
| | *kdrl* | 20× | Alexa514 | 518 | HyD S 1, S 2, S 3 | 523–543, 543–563, 563–583 | 0.568 × 0.568 × 4.0 | 2C, S2–S3 |
| | *shha* | 20× | Alexa546 | 557 | HyD S 1, S 2, S 3 | 566–580, 580–600, 600–620 | 0.568 × 0.568 × 4.0 | 2C, S2–S3 |
| | *elavl3* | 20× | Alexa594 | 590 | HyD S 1, S 2, S 3 | 600–620, 620–640, 640–660 | 0.568 × 0.568 × 4.0 | 2C, S2–S3 |
| | *sox10* | 20× | Atto633 | 629 | HyD S 2, S 3 | 640–660, 660–680 | 0.568 × 0.568 × 4.0 | 2C, S2–S3 |
| | *ntla* | 20× | Alexa700 | 686 | HyD S 2 | 696–723 | 0.568 × 0.568 × 4.0 | 2C, S2–S3 |
| | *dmd* | 20× | Alexa750 | 755 | HyD S 2, S 3 | 765–780, 780–795 | 0.568 × 0.568 × 4.0 | 2C, S2–S3 |
| | *col2a1a* | 20× | iFluor800 | 790 | HyD S 3, R 5 | 815–830, 835–850 | 0.568 × 0.568 × 4.0 | 2C, S2–S3 |
| | Autofluorescence | 20× | – | 459 | HyD S 1, S 2, S 3, X 4 | 465–500, 500–535, 535–570, 570–630 | 0.568 × 0.568 × 4.0 | 2C, S2–S3 |
| Whole-mount embryo | *col2a1a* | 63× | iFluor800 | 790 | HyD S 3 , R 5 | 815–830, 835–850 | 0.180 × 0.180 × 1.2 | 3B, S6, S11, S16 |
| | *mylpfa* | 63× | Atto425 | 440 | HyD S 1, S 2, S 3 | 450–475, 475–495, 495–520 | 0.180 × 0.180 × 1.2 | 3B, S7, S12, S16 |
| | *elavl3* | 63× | Alexa594 | 590 | HyD S 1, S 2, S 3 | 600–620, 620–640, 640–660 | 0.180 × 0.180 × 1.2 | 3B, S8, S13, S16 |
| | *kdrl* | 63× | Alexa514 | 518 | HyD S 1, S 2, S 3 | 523–543, 543–563, 563–583 | 0.180 × 0.180 × 1.2 | 3B, S9, S14, S16 |
| | *dmd* | 63× | Alexa750 | 755 | HyD S 2, S 3 | 765–780, 780–795 | 0.180 × 0.180 × 1.2 | 3B, S10, S15, S16 |
| | Autofluorescence | 63× | – | 459 | HyD S 1, S 2, S 3, X 4 | 465–500, 500–535, 535–570, 570–630 | 0.180 × 0.180 × 1.2 | 3B, S6–S16 |
| Whole-mount embryo | *kdrl* | 63× | Alexa514 | 518 | HyD S 1, S 2, S 3 | 523–543, 543–563, 563–583 | 0.180 × 0.180 × 1.2 | 4, S17 |
| Brain section | NFH | 20× | Alexa405 | 405 | HyD S 1 | 410–450 | 0.568 × 0.568 × 4.0 | 5C, S20–S23 |
| | CD31 | 20× | Atto425 | 440 | HyD S 1, S 2, S 3 | 450–475, 475–495, 495–520 | 0.568 × 0.568 × 4.0 | 5C, S20–S23 |
| | *Slc17a7* | 20× | Alexa488 | 488 | HyD S 1, S 2 | 493–513, 513–533 | 0.568 × 0.568 × 4.0 | 5C, S20–S23 |
| | *Gad1* | 20× | Alexa514 | 518 | HyD S 1, S 2, S 3 | 523–543, 543–563, 563–583 | 0.568 × 0.568 × 4.0 | 5C, S20–S23 |
| | *Sst* | 20× | Alexa546 | 557 | HyD S 1, S 2, S 3 | 566–580, 580–600, 600–620 | 0.568 × 0.568 × 4.0 | 5C, S20–S23 |
| | *Actb* | 20× | Alexa594 | 590 | HyD S 1, S 2, S 3 | 600–620, 620–640, 640–660 | 0.568 × 0.568 × 4.0 | 5C, S20–S23 |
| | *Lamp5* | 20× | Atto633 | 629 | HyD S 2, S 3 | 640–660, 660–680 | 0.568 × 0.568 × 4.0 | 5C, S20–S23 |
| | *Plp1* | 20× | Alexa700 | 686 | HyD S 2 | 696–723 | 0.568 × 0.568 × 4.0 | 5C, S20–S23 |
| | *Vip* | 20× | Alexa750 | 755 | HyD S 2, S 3 | 765–780, 780–795 | 0.568 × 0.568 × 4.0 | 5C, S20–S23 |
| | RBFOX3 | 20× | iFluor800 | 790 | HyD S 3, R 5 | 815–830, 835–850 | 0.568 × 0.568 × 4.0 | 5C, S20–S23 |
| | Autofluorescence | 20× | – | 459 | HyD S 1, S 2, S 3, X 4 | 465–500, 500–535, 535–570, 570–630 | 0.568 × 0.568 × 4.0 | 5C, S20–S23 |

**Table S3. Microscope settings.** Confocal microscopy was performed with a Leica Stellaris 8 inverted confocal microscope. Objectives were as follows: HC PL APO 20×/0.75 IMM CORR CS2 (catalog # 11506343), HC PL APO 63×/1.40 OIL CS2 (catalog # 11506350); both utilized with oil immersion.

#### S1.4 Image analysis

We build on an image analysis framework developed over a series of publications (Choi *et al.*, 2010; Choi *et al.*, 2014; Choi *et al.*, 2016; Trivedi *et al.*, 2018; Schwarzkopf *et al.*, 2021). For convenience, here we provide a self-contained description of the details relevant to the present work.

##### S1.4.1 Raw pixel intensities

The total fluorescence within a pixel is a combination of signal and background. Fluorescent background (BACK) arises from three sources in each channel:

- autofluorescence (AF): fluorescence inherent to the sample.
- non-specific detection (NSD): probes that bind non-specifically in the sample and subsequently trigger HCR amplification. For HCR IF experiments using both primary antibody probes and secondary antibody probes, $NSD_{1^\circ}$ arises from non-specific binding of primary antibody probes and $NSD_{2^\circ}$ arises from non-specific binding of secondary antibody probes, with NSD = $NSD_{1^\circ}$ + $NSD_{2^\circ}$.
- non-specific amplification (NSA): HCR hairpins that bind non-specifically in the sample.

Fluorescent signal (SIG) in each channel corresponds to:

- signal (SIG): probes that bind specifically to the target and subsequently trigger HCR amplification.

For pixel $i$ of replicate sample $n$, we denote the background

$$X_{n,i}^{\text{BACK}} = X_{n,i}^{\text{NSD}} + X_{n,i}^{\text{NSA}} + X_{n,i}^{\text{AF}}, \tag{S1}$$

the signal:

$$X_{n,i}^{\text{SIG}}, \tag{S2}$$

and the total fluorescence (SIG+BACK):

$$X_{n,i}^{\text{SIG+BACK}} = X_{n,i}^{\text{SIG}} + X_{n,i}^{\text{BACK}}. \tag{S3}$$

##### S1.4.2 Measurement of signal, background, and signal-to-background

For each of 10 linearly unmixed channels, background (BACK) is characterized for pixels in one or more representative rectangular regions of no- or low-expression for the corresponding target and the combination of signal plus background (SIG+BACK) is characterized for pixels in one or more representative rectangular regions of high expression for the corresponding target (e.g., Figures S3, S16, S23). For the pixels in these regions, we characterize the distribution by plotting an intensity histogram and characterize average performance by calculating the mean pixel intensities

$$\bar{X}_n^{\text{BACK}}, \qquad \bar{X}_n^{\text{SIG+BACK}} \tag{S4}$$

for replicate $n$. Performance across replicates is characterized by calculating the sample means

$$\bar{X}^{\text{BACK}}, \qquad \bar{X}^{\text{SIG+BACK}} \tag{S5}$$

and standard error of the mean

$$s_{\bar{X}^{\text{BACK}}}, \qquad s_{\bar{X}^{\text{SIG+BACK}}}. \tag{S6}$$

The mean signal is then estimated as

$$\bar{X}^{\text{SIG}} = \bar{X}^{\text{SIG+BACK}} - \bar{X}^{\text{BACK}} \tag{S7}$$

with the standard error of the mean estimated via uncertainty propagation as

$$s_{\bar{X}^{\text{SIG}}} \leq \sqrt{(s_{\bar{X}^{\text{SIG+BACK}}})^2 + (s_{\bar{X}^{\text{BACK}}})^2}. \tag{S8}$$

The signal-to-background ratio is estimated as:

$$\bar{X}^{\text{SIG/BACK}} = \bar{X}^{\text{SIG}} / \bar{X}^{\text{BACK}} \tag{S9}$$

with standard error of the mean estimated via uncertainty propagation as

$$s^{\text{SIG/BACK}} \leq \bar{X}^{\text{SIG/BACK}} \sqrt{\left(\frac{s_{\bar{X}^{\text{SIG}}}}{\bar{X}^{\text{SIG}}}\right)^2 + \left(\frac{s_{\bar{X}^{\text{BACK}}}}{\bar{X}^{\text{BACK}}}\right)^2}. \tag{S10}$$

These upper bounds on estimated standard errors hold under the assumption that the correlation between SIG and BACK is non-negative.

###### S1.4.3 Normalized voxel intensities for qHCR imaging: analog RNA relative quantitation with subcellular resolution in an anatomical context

For quantitative imaging using HCR, precision increases with voxel size as long as the imaging voxels remain smaller than the features in the expression pattern (see Section S2.2 of (Trivedi *et al.*, 2018)). To increase precision, we calculate raw voxel intensities by averaging neighboring pixel intensities while still maintaining a subcellcular voxel size. To facilitate relative quantitation between voxels, we estimate the normalized HCR signal of voxel $j$ in replicate $n$ as:

$$x_{n,j} \equiv \frac{X_{n,j}^{\text{SIG+BACK}} - X^{\text{BOT}}}{X^{\text{TOP}} - X^{\text{BOT}}}, \tag{S11}$$

which translates and rescales the data so that the voxel intensities in each channel fall in the interval [0,1]. Here,

$$X^{\text{BOT}} \equiv \bar{X}^{\text{BACK}} \tag{S12}$$

is the mean background across replicates (see Section S1.4.2) and

$$X^{\text{TOP}} \equiv \max_{n,j} X_{n,j}^{\text{SIG+BACK}} \tag{S13}$$

is the maximum total fluorescence for a voxel across replicates.

Pairwise expression scatter plots that each display normalized voxel intensities for two channels (e.g., Figures 4 and 5 of (Trivedi *et al.*, 2018)) provide a powerful quantitative framework for performing multidimensional read-out/read-in analyses (Figure 6 of (Trivedi *et al.*, 2018)). Read-out from anatomical space to expression space enables discovery of expression clusters of voxels with quantitatively related expression levels and ratios (amplitudes and slopes in the expression scatter plots), while read-in from expression space to anatomical space enables discovery of the corresponding anatomical locations of these expression clusters within the embryo. The simple and practical normalization approach of (S11)–(S13) translates and rescales all voxels identically within a given channel (enabling comparison of amplitudes and slopes in scatter plots between replicates), and does not attempt to remove scatter in the normalized signal estimate that is caused by scatter in the background.

To validate qHCR spectral imaging with subcellular resolution ($2.0\times2.0\times1.2$ $\mu$m voxels) in whole-mount zebrafish embryos, Figures 3B and S6–S10 display highly correlated normalized voxel intensities for 2-channel redundant detection of five target RNAs. In this setting, accuracy corresponds to linearity with zero intercept, and precision corresponds to scatter around the line (Trivedi *et al.*, 2018). To address chromatic aberration resulting from the wide range of wavelengths used in spectral imaging, Huygens Software (Scientific Volume Imaging) was used to apply a chromatic aberration correction across the 10 channels. As a point of reference, Figures S11–S15 display normalized voxel intensities for 2-channel redundant detection of five target RNAs without chromatic aberration correction (cf. Figures S6–S10).

##### S1.4.4 Dot detection and colocalization for dHCR imaging: digital RNA absolute quantitation in an anatomical context

To validate the performance of spectral imaging with linear unmixing for single-molecule imaging, we perform a 2-channel redundant detection experiment in which target RNA *kdrl* is detected using two independent probe sets and HCR amplifiers in Ch4 and Ch7 of a 10-plex experiment. Let $N_4$ denote the number of dots detected in Ch4, $N_7$ the number of dots detected in Ch7, and $N_{47}$ the number of colocalized dots appearing in both channels. We define the colocalization fraction for each channel:

$$C_4 = N_{47}/N_4 \tag{S14}$$

$$C_7 = N_{47}/N_7 \tag{S15}$$

As the false-positive and false-negative rates for single-molecule detection go to zero, $C_4$ and $C_7$ will both approach 1 from below, providing a quantitative basis for evaluating performance.

Single molecules were identified in each channel using the Dot Detection 2.0 software package. The Dot Detection 2.0 software package employs the following dot detection algorithm applied to a three-dimensional confocal image stack $I$ with pixel intensity $I(\vec{r})$ at 3D location $\vec{r}$:

- **Step 1: Remove smoothly varying features.** To eliminate variations in pixel intensity arising from background variations that occur on a length scale much larger than the dots of interest, local background subtraction is performed by convolving $I$ with an isotropic Gaussian (standard deviation $\sigma_{\text{smooth}}$) to obtain a blurred image. Then for each pixel $i$ at a 3D location $\vec{r}_i$, the blurred value is subtracted from $I(\vec{r}_i)$, and any resulting negative values are removed:

$$I'(\vec{r}_i) = \max(0, I(\vec{r}_i) - [\mathcal{N}_{\vec{0},\sigma_{\text{smooth}}} * I](\vec{r}_i)) \tag{S16}$$

- **Step 2: Identify candidate dots.** Candidate dots are accentuated by application of a determinant-of-Hessian filter to the image, with typical dot size characterized by standard deviation $\sigma$. The determinant-of-Hessian filter is a rotation-invariant filter used for blob detection (Beaudet, 1978):

$$I_{\text{DoH}}(\vec{r}_i) = -\begin{vmatrix} [\frac{\partial^2}{\partial x^2}\mathcal{N}_{\vec{0},\sigma} * I'](\vec{r}_i) & [\frac{\partial^2}{\partial x \partial y}\mathcal{N}_{\vec{0},\sigma} * I'](\vec{r}_i) & [\frac{\partial^2}{\partial x \partial z}\mathcal{N}_{\vec{0},\sigma} * I'](\vec{r}_i) \\ [\frac{\partial^2}{\partial y \partial x}\mathcal{N}_{\vec{0},\sigma} * I'](\vec{r}_i) & [\frac{\partial^2}{\partial y^2}\mathcal{N}_{\vec{0},\sigma} * I'](\vec{r}_i) & [\frac{\partial^2}{\partial y \partial z}\mathcal{N}_{\vec{0},\sigma} * I'](\vec{r}_i) \\ [\frac{\partial^2}{\partial z \partial x}\mathcal{N}_{\vec{0},\sigma} * I'](\vec{r}_i) & [\frac{\partial^2}{\partial z \partial y}\mathcal{N}_{\vec{0},\sigma} * I'](\vec{r}_i) & [\frac{\partial^2}{\partial z^2}\mathcal{N}_{\vec{0},\sigma} * I'](\vec{r}_i) \end{vmatrix} \tag{S17}$$

  All local maxima are then found in the resulting image $I_{\text{DoH}}$. A local maximum is defined as a pixel $i$ at 3D location $\vec{r}_i$ whose intensity $I_{\text{DoH}}(\vec{r}_i)$ is greater than or equal to the intensities of all face-neighboring pixels (e.g., an interior pixel in a 3D image will have 6 face neighbors):

$$R^{\max'} = \left\{\vec{r}_i \mid I_{\text{DoH}}(\vec{r}_i) \geq I_{\text{DoH}}(\vec{s}) \;\forall\; \vec{s} \in \textsc{Neighbors}(\vec{r}_i)\right\} \tag{S18}$$

  For computational expediency, only the highest $n_{\text{maxima}}$ local maxima are retained. Subsequently, if a local maximum is within $r_{\min}$ of another local maximum with a higher value, the lower local maximum is discarded, forming the resultant set of dot centroids $R^{\max} \subseteq R^{\max'}$.

- **Step 3: Estimate dot weights.** We approximate the image $I'$ using a weighted Gaussian for each dot, with the weight of each dot constrained to be non-negative. Given the set of dot locations $R^{\max}$, the set of optimal weights is obtained by solving the minimization problem:

$$\arg\min_{w} \sum_i \Big(\sum_j w_j \cdot \mathcal{N}_{R_j^{\max},\sigma}(\vec{r}_i) - I'(\vec{r}_i)\Big)^2 \text{ such that } w_j \geq 0 \;\forall\; j \tag{S19}$$

  where $\mathcal{N}_{R_i^{\max},\sigma}(\vec{r}_i)$ is the value of a unit 3D Gaussian with centroid $R_j^{\max}$ and standard deviation $\sigma$ evaluated at location $\vec{r}_i$, and $I'(\vec{r}_i)$ is intensity of pixel $i$ located at $\vec{r}_i$. This optimization takes the form of a non-negative

| Parameter | Value | Description |
|---|---|---|
| $\sigma$ | 0.3 $\mu$m | Approximate expected dot radius |
| $\sigma_{\mathrm{smooth}}$ | 40 $\mu$m | Lengthscale of smoothly varying features |
| $n_{\mathrm{candidates}}$ | 2,000 | Upper bound on the number of dots |
| $n_{\mathrm{maxima}}$ | 100,000 | Upper bound on the number of local maxima considered |
| $r_{\mathrm{min}}$ | 0.94 $\mu$m | Minimum separation between dots in the same channel |
| $r_{xy}$ | 0.5 $\mu$m | Maximum separation in $xy$ for dots to be colocalized between channels |
| $r_z$ | 1.0 $\mu$m | Maximum separation in $z$ for dots to be colocalized between channels |
| $f_{\mathrm{tol}}$ | $10^{-8}$ | Tolerance for NNLS convergence |
| $g_{\mathrm{tol}}$ | $10^{-8}$ | Tolerance for L-BFGS-B convergence |
| $f_{\mathrm{trunc}}$ | 5.0 | Number of standard deviations at which Gaussians are truncated |
| $t_{\mathrm{dot}}$ | see below | Threshold on dot weight |

**Table S4.** Summary of parameters used for dot detection and colocalization

least squares (NNLS) problem, which can be efficiently solved using Algorithm 1 by calling $\mathrm{NNLS}(A, b, f_{\mathrm{tol}})$ with input matrix $A$ and vector $b$:

$$A_{j,k} = \sum_i \mathcal{N}_{R_j^{\mathrm{max}},\sigma}(\vec{r}_i) \cdot \mathcal{N}_{R_k^{\mathrm{max}},\sigma}(\vec{r}_i) \tag{S20}$$

$$b_j = \sum_i \mathcal{N}_{R_j^{\mathrm{max}},\sigma}(\vec{r}_i) \cdot I'(\vec{r}_i) \tag{S21}$$

A sparse matrix representation is used for $A$ to enable consideration of large numbers of local maxima. For computational expediency, only the $n_{\mathrm{candidates}}$ dots with the largest optimized weights are retained.

- **Step 4: Refine dot centroids and weights.** We next optimize the dot centroids to have sub-pixel resolution and update the corresponding weights. Specifically, we minimize the sum of squared residuals between the image $I'$ and the combined dot Gaussians:

$$\min_{R,w} \sum_i \Big( \sum_j w_j \cdot \mathcal{N}_{R_j,\sigma}(\vec{r}_i) - I'(\vec{r}_i) \Big)^2 \text{ such that } w_j \geq 0 \; \forall \; j \tag{S22}$$

This minimization is performed as an outer optimization over dot centroids $R$:

$$R^{\mathrm{opt}} = \arg\min_R \sum_i \Big( \sum_j w_j^{\mathrm{opt}}(R) \cdot \mathcal{N}_{R_j,\sigma}(\vec{r}_i) - I'(\vec{r}_i) \Big)^2 \tag{S23}$$

with all dot centroids constrained to remain within the volume of the image stack, and an inner optimization over dot weights $w$:

$$w^{\mathrm{opt}}(R) = \arg\min_w \Big( \sum_j w_j \cdot \mathcal{N}_{R_j,\sigma}(\vec{r}_i) - I'(\vec{r}_i) \Big)^2 \text{ such that } w_j \geq 0 \; \forall \; j \tag{S24}$$

The inner NNLS problem is solved efficiently using Algorithm 1 with inputs analogous to (S20) and (S21). The outer nonlinear optimization problem is solved using the L-BFGS-B algorithm (Zhu *et al.*, 1997) (convergence criterion: $L^\infty$ norm of the gradient less than or equal to $g_{\mathrm{tol}}$). After optimization, if a dot centroid is within $r_{\mathrm{min}}$ of another dot centroid corresponding to a higher weight, the dot with the lower weight is discarded.

- **Step 5: Eliminate dim dots.** To eliminate dim dots, any dot with a weight $w^{\mathrm{opt}}(R^{\mathrm{opt}})$ less than or equal to $t_{\mathrm{dot}}$ is discarded. The resulting number of dots $N_c$ is recorded for channel $c$.

In all steps, Gaussians are truncated at $f_{\text{trunc}}$ standard deviations (whether $\sigma$ or $\sigma_{\text{smooth}}$).

After identifying the dots in each channel of a 2-channel redundant detection image, a dot $j$ in Channel A and a dot $k$ in Channel B are considered colocalized if all of the following four statements were true:

- **Test 1:** The $xy$ centroids (taken from $R^{\text{opt}}$) differed by less than a lateral distance threshold ($r_{xy}$).
- **Test 2:** Dot $j$ is the closest dot in Channel A to dot $k$ in Channel B.
- **Test 3:** Dot $k$ is the closest dot in Channel B to dot $j$ in Channel A.
- **Test 4:** The $z$ centroids (taken from $R^{\text{opt}}$) differed by less than the axial distance threshold ($r_z$).

Note that due to the lower axial resolution, dot colocalization was tested separately for $xy$ and $z$. The dot weight threshold $t_{\text{dot}}$ was set as follows for (Ch4, Ch7) for Replicates 1-3:

- Replicate 1: $(250, 400)$
- Replicate 2: $(250, 300)$
- Replicate 3: $(290, 350)$

###### S1.4.5 Linear unmixing

For convenience, we perform linear unmixing using the Leica LAS X software available with the Leica Stellaris 8 microscope. Alternatively, for generality, our Unmix 1.0 software package can be used to perform the linear unmixing step.

Consider a 10-plex sample of interest containing a total of 10 different target RNAs and/or proteins, each stained by a different fluorophore. Spectrally imaging a set of 10 reference samples that each contain only one type of fluorophore provides a reference spectrum for each channel. Spectrally imaging an unlabeled sample provides a reference spectrum for autofluorescence (treated as an 11th channel). Each pixel in a spectral image of a 10-plex sample of interest may then be interpreted as a linear superposition of these 11 reference spectra with an unknown coefficient for each channel. Linear unmixing of the fluorescence in that pixel then determines the non-negative coefficients that best approximate the spectrum of the pixel, revealing the fluorescence contribution of each channel (Mansfield *et al.*, 2008). For each pixel in a 10-plex image, the algorithm uses 11 reference spectra (e.g., Figures 1C, S1, S19; one per fluorophore and one for AF) to linearly unmix the fluorescence in that pixel by solving a non-negative least squares (NNLS) optimization problem (Franc *et al.*, 2005; Garini *et al.*, 2006) to identify the coefficient for each of 11 fluorescence channels (one per fluorophore and one for AF). The output of linear unmixing is a set of 11 unmixed fluorescence channels for all pixels.

The Unmix 1.0 software package solves the NNLS optimization problem following the approach of (Franc *et al.*, 2005). Given a reference spectrum matrix $F$ (with $n_{\text{detector}}$ rows and $n_{\text{channel}}$ columns) and a set of fluorescence measurements for a single pixel arranged as a column vector $y$ (with $n_{\text{detector}}$ elements) we solve a constrained minimization over the channel coefficients arranged as a column vector $x$ (with $n_{\text{channel}}$ elements):

$$\min_x ||Fx - y||^2 \quad \text{such that} \quad x_i \geq 0 \;\forall\, i. \tag{S25}$$

Expanding the norm yields the equivalent minimization problem:

$$\min_x x^T F^T F x - 2y^T F x \quad \text{such that} \quad x_i \geq 0 \;\forall\, i. \tag{S26}$$

where $F^T F$ is a positive semi-definite symmetric matrix. For a single pixel in a spectral image, the vector $x$ of $n_{\text{channel}}$ coefficients is obtained by calling NNLS($F^T F, F^T y, f_{\text{tol}}$) using Algorithm 1 below, which proceeds via coordinate descent. This algorithm is called once for each pixel in a spectral image, yielding $n_{\text{channel}}$ coefficients for each pixel. The tolerance was set to $f_{\text{tol}} = 10^{-8}$.

**Algorithm 1** Non-negative least squares algorithm via coordinate descent

```
function NNLS(A, b, f_tol)
    x ← 0⃗
    m ← b − A^T x
    y_1 ← −(m + b)^T x
    while true do
        for k = 1, …, n_cols(A) do
            t ← max(0, x_k + m_k / A_{k,k})
            m ← m + (x_k − t) A_{:,k}
            x_k ← t
        y_0 ← y_1
        y_1 ← −(m + b)^T x
        if (y_0 − y_1) ≤ |y_1| · f_tol^2 then
            return x
```

### S2 Protocols

#### S2.1 Protocol for 10-plex HCR spectral imaging and linear unmixing

##### S2.1.1 Spectral imaging and linear unmixing workflow overview

1. **HCR RNA-FISH/IF (see the protocols of Sections S2.2 and S2.3).** Prepare 12 sample types (HCR RNA-FISH and/or HCR IF for a total of 10 RNA and/or protein targets):
   (a) 10-plex sample or multiple replicate samples (1 fluorophore for each of 10 targets): use all 10 HCR probe sets and amplifiers.
   (b) 1-plex reference sample for each of 10 targets: use the corresponding HCR probe set and amplifier for a given target.
   (c) Two unlabeled autofluorescence (AF) samples: omit all HCR probe sets and amplifiers.
2. **AF scan (see Section S2.1.2).** Use one unlabeled sample to perform an excitation-emission scan to determine the maximal AF excitation wavelength, which in turn determines a set of optimized detection wavelengths using 4 detectors.
3. **Spectral imaging (see Section S2.1.3).** Spectrally image 12 sample types using 11 excitation wavelengths (one optimized for each fluoropohore and one optimized for AF):
   (a) 10-plex sample.
   (b) 1-plex samples (obtain reference sample for each fluorophore).
   (c) Unlabeled sample (obtain reference spectrum for AF).
4. **Linear unmixing (see Section S2.1.4).** Use the 11 reference spectra (one per fluorophore and one for AF) to linearly unmix the 10-plex image to obtain 11 unmixed channels (one per fluorophore and one for AF).

##### S2.1.2 Protocol for performing autofluorescence excitation-emission scan using Leica Stellaris 8

1. Open the LAS X software (this protocol was developed using version 4.5).

2. At the top of the window, click “Configuration” > “Hardware” > change “Bit Depth” to 16.

3. Click “Acquire” at the top of the window to return to the image acquisition screen.

4. Place one of the autofluorescence samples on the microscope to conduct an excitation-emission scan (also known as a $\Lambda\lambda$ scan) to determine the optimal autofluorescence excitation wavelength.

5. In the left panel, under “Acquisition Mode”, change “xyz” to “xy$\Lambda\lambda$”.

6. In the left panel, in the “$\Lambda\lambda$: Excitation Emission Scan Settings” sub-panel, click the plus sign in the upper lefthand corner.

7. In the pop-up menu, click “Reset Values to Default”.

8. In the pop-up menu, in the following order, use the mouse scroll wheel to set “Excitation Steps” to 18 (this will automatically also set “Excitation Stepsize” to 20 nm), “Detection Steps” to 14, “Detection Bandwidth” to 20 nm, and “Detection Stepsize” to 24 nm. Close the pop-up menu.

9. Under the eyepiece, navigate to the region of the autofluorescence sample with the most intense autofluorescence.

10. Enter “Live” imaging mode, and adjust the laser intensity and/or detector gain so that the highest pixel intensity is approximately 25% of the maximum possible pixel intensity. The laser line and HyD S 1 detector may need to be moved to different wavelengths to see the autofluorescence.
    NOTE: *Keeping the pixel intensities low ensures pixel saturation will not occur during the $\Lambda\lambda$ scan, as pixel saturation would obscure the spectral information.*

11. Click “Start” to begin the excitation-emission scan.

12. When the scan is finished, click on the “LambdaLambda 001” file under “Open projects” in the left panel.

13. Click “Process” at the top of the window.

14. Click “Excitation / Emission Contour Plot” in the left panel.

15. In the right panel, drag the “t” slider so that the sample is visible. The “$\Lambda$” slider and pixel intensity slider may also need to be adjusted to make the sample visible.

16. In the right panel, click the Rectangle, Oval, or Polygon button at the top of the screen, and draw a region in the image around the brightest autofluorescence.

17. In the middle panel, click “Apply” at the bottom of the screen. This will display a contour plot.

18. Reposition the crosshairs to the maximum of the contour plot, and make note of the Excitation wavelength displayed at the bottom right of the plot. This wavelength, henceforth denoted as $\lambda_{\mathrm{AF}}$, will serve as the optimal autofluorescence excitation wavelength.
    NOTE: *Going forward, if the sample preparation protocol remains the same, the optimal autofluorescence excitation wavelength ($\lambda_{AF}$) determined here can be used for future batches of experiments with this sample type, and this excitation-emission scan does not need to be repeated for each batch.*

##### S2.1.3 Protocol for spectral imaging using Leica Stellaris 8

1. Open the LAS X software (this protocol was developed using version 4.5).

2. At the top of the window, click “Configuration” > “Hardware” > and make sure “Bit Depth” is set to 16.

3. Click “Acquire” at the top of the window to return to the image acquisition screen.

4. Click “Acquisition” in the left panel, and make sure “Acquisition Mode” is set to “xyz”.

5. In the middle panel of the software, create 11 Settings. Settings 1-10 are used to image the target fluorophores, while Setting 11 is used to image autofluorescence.

6. Configure Settings 1-10 as follows:

| Setting | Laser line (nm) | Detector(s) | Detector wavelengths (nm) | Corresponding fluorophore |
|---|---|---|---|---|
| 1 | 405 | HyD S 1 | 410–450 | Alexa405 |
| 2 | 440 | HyD S 1, S 2, S 3 | 450–475, 475–495, 495–520 | Atto425 |
| 3 | 488 | HyD S 1, S 2 | 493–513, 513–533 | Alexa488 |
| 4 | 518 | HyD S 1, S 2, S 3 | 523–543, 543–563, 563–583 | Alexa514 |
| 5 | 557 | HyD S 1, S 2, S 3 | 566–580, 580–600, 600–620 | Alexa546 |
| 6 | 590 | HyD S 1, S 2, S 3 | 600–620, 620–640, 640–660 | Alexa594 |
| 7 | 629 | HyD S 2, S 3 | 640–660, 660–680 | Atto633 |
| 8 | 686 | HyD S 2 | 696–723 | Alexa700 |
| 9 | 755 | HyD S 2, S 3 | 765–780, 780–795 | Alexa750 |
| 10 | 790 | HyD S 3, R 5 | 815–830, 835–850 | iFluor800 |

**Table S5. Settings 1-10 configurations for spectral imaging.**

7. For Setting 11 (autofluorescence), set the laser line to the optimal autofluorescence excitation wavelength ($\lambda_{AF}$) determined via the AF excitation-emission scan above.

8. For Setting 11, activate the HyD S 1, S 2, S 3, and X 4 detectors. The detector wavelengths will be determined by $\lambda_{AF}$. Configure the detectors for Setting 11 as follows:

| Detector | Lower wavelength (nm) | Upper wavelength (nm) |
|---|---|---|
| HyD S 1 | $\lambda_{AF}$ + 6 | $\lambda_{AF}$ + 41 |
| HyD S 2 | $\lambda_{AF}$ + 41 | $\lambda_{AF}$ + 76 |
| HyD S 3 | $\lambda_{AF}$ + 76 | $\lambda_{AF}$ + 111 |
| HyD X 4 | $\lambda_{AF}$ + 111 | $\lambda_{AF}$ + 171 |

**Table S6. Setting 11 configuration for spectral imaging.** $\lambda_{AF}$: optimal autofluorescence excitation wavelength.

For example, if the optimal autofluorescence excitation wavelength ($\lambda_{AF}$) was measured to be 459 nm, the Setting 11 detectors would be configured as follows:

- HyD S 1: 465–500 nm
- HyD S 2: 500–535 nm
- HyD S 3: 535–570 nm
- HyD X 4: 570–630 nm

9. Place the 10-plex sample on the microscope.

10. For each of the 11 Settings:

(a) Navigate to the position in the sample that has the maximum intensity for the fluorophore corresponding to that Setting.

(b) For the linear unmixing to perform properly, it is important that no pixels are saturated. Therefore, while using the "Live" mode, set the laser intensity and detector gain(s) for that Setting so that the maximum pixel intensity for each detector is no more than 50% of the maximum possible value. Do not change the laser wavelength or detector wavelengths; the Format, Speed, Zoom, Averaging, and Accumulation settings may be adjusted as needed.
NOTE: *Each detector collects emissions spectra over a range of wavelengths. For some Settings, multiple detectors are utilized to collect a broader range of emissions spectra for a given fluorophore.*

(c) Click "Capture Image", and double-check that the captured image reaches no more than 50% of saturation for all pixels in all detectors. The captured image may then be deleted.

11. As a final check, to ensure that no pixels will become saturated, while in "Live" mode, traverse the entire sample and verify that no pixel intensities exceed 50% of saturation in any of the detectors. Because it is possible that the laser for one Setting can cross-excite a fluorophore corresponding to a neighboring Setting, be sure to check that the chosen laser intensities and detector gains do not result in pixel intensities above 50% of saturation in the detectors of neighboring Settings. Decrease the laser intensities and/or detector gains if any pixel intensities are too high.

12. Now that the laser intensity and detector gain settings are determined for all 11 Settings, do not change the laser intensity or detector gain settings again.

13. Collect a Z-stack for the 10-plex sample by setting the "Begin" and "End" locations for a Z-stack and clicking "Start".

14. One by one, place each of the 10 reference spectrum samples on the microscope. Find the area of the sample with the brightest fluorescence, and collect a single Z-section at that location. Rename each image file to indicate the target name and fluorophore number.
NOTE: *All 11 Settings should still be active when collecting the reference spectrum sample images.*

15. Place the other autofluorescence sample on the microscope. Find the area of the sample with the brightest autofluorescence, and collect a single Z-section at that location. Rename the image file to "autofluorescence".
NOTE: *All 11 Settings should still be active when collecting the autofluorescence sample image.*

###### S2.1.4 Protocol for linear unmixing using the LAS X software

1. Click "Process" along the top of the window.

2. Within the "ProcessTools" menu in the left panel, click "Channel Dye Separation" (under "Dye Separation").

3. Click "Open Projects" at the top of the left panel.

4. One by one, for each reference spectrum file and the autofluorescence image file:

   (a) Click on the image file in the left panel.

   (b) In the right panel, look at the image(s) corresponding to the Setting for that sample, and reposition and resize the circular region selector so that it covers the brightest region of the target (or the brightest region of the autofluorescence for the autofluorescence sample). Avoid including pixels that are outside the brightest region to prevent corruption of the fluorophore spectrum.

   (c) Click "Add" near the bottom of the middle panel. This records a reference spectrum for the fluorophore.

5. Click "Save Matrix" in the middle panel, give the matrix a descriptive name, and click "Save".

6. Click on the 10-plex image file in the left panel.

7. Click “ProcessTools” at the top of the left panel.

8. Click “Automatic Dye Separation” in the left panel.

9. Within the middle panel, under “Method”, click the “Manual” circle, which allows the matrix to be loaded.

10. Enter “11” for “Fluorescent Dyes” in the middle panel.
    NOTE: *Rescale should be left as “All Channels”.*

11. Click on the dividing line between the right and middle panels of the software and drag it all the way to the right (thereby making the panel with the 26 channels of images as small as possible). This reveals a button at the bottom of the middle panel labeled “Load”. Click the “Load” button.

12. Navigate to the saved matrix file, click on the matrix file, and click “Open”.

13. Click the “Apply” button located to the right of the “Load” button.

14. Click the “Apply” button at the bottom of the screen to unmix the 10-plex image.

15. To view the unmixed image, click “Acquire” at the top of the window. In the left panel, the 11-channel unmixed image (one channel per target plus one channel for autofluorescence) will have appeared with “DyeSep” added near the end of the file name.

16. Save the project.

#### S2.2 Protocols for HCR RNA-FISH in whole-mount zebrafish embryos

Protocols for HCR RNA-FISH in whole-mount zebrafish embryos are adapted from (Choi *et al.*, 2016; Choi *et al.*, 2018). This protocol has been optimized for embryos at 27 hpf; other developmental stages may require additional optimization.

##### S2.2.1 Preparation of fixed whole-mount zebrafish embryos

1. Collect zebrafish embryos and incubate at 28 °C in a Petri dish with egg $H_2O$.
   NOTE: *Collect no more than 100 embryos per Petri dish.*

2. Replace the egg $H_2O$ with fresh egg $H_2O$ 6 hours after collecting the embryos.

3. At 27 hours post-fertilization (hpf), dechorionate the embryos by replacing the egg $H_2O$ with 1 mg/mL pronase solution. After 5 min, gently pipet up and down with a glass pipette to dechorionate the embryos.

4. Remove dechorionated embryos to a Petri dish with fresh egg $H_2O$.

5. Gently wash the dechorionated embryos twice with egg $H_2O$.

6. Transfer up to 80 embryos to a 2 mL Eppendorf tube and remove excess egg $H_2O$.

7. Fix embryos in 2 mL of 4% paraformaldehyde (PFA) for 24 h at 4 °C.
   CAUTION: *Use PFA with extreme care, as it is a hazardous material.*
   NOTE: *Cool PFA to 4 °C before use.*

8. Wash embryos 3 × 5 min with 1 mL of 1× phosphate-buffered saline (PBS) to stop the fixation.
   NOTE: *Avoid using calcium chloride and magnesium chloride in PBS, as this leads to increased autofluorescence in the samples.*

9. Dehydrate and permeabilize with a series of methanol (MeOH) washes (1 mL each):
   (a) 100% MeOH for 4 × 10 min
   (b) 100% MeOH for 1 × 50 min.

10. Remove the final 100% MeOH wash.

11. Add 1 mL 100% MeOH and store embryos overnight at −20 °C before use.
    NOTE: *Embryos can be stored for at least one year at −20 °C.*

##### S2.2.2 Buffer recipes for sample preparation

| | |
|---|---|
| **10 mg/mL pronase stock solution** | For 10 mL of solution |
| 10 mg/mL pronase | 100 mg of pronase powder |
| | Fill up to 10 mL with ultrapure $H_2O$ |
| **1 mg/mL pronase solution** | For 25 mL of solution |
| 1 mg/mL pronase | 2.5 mL of 10 mg/mL pronase stock solution |
| egg $H_2O$ | Fill up to 25 mL with egg $H_2O$ |
| **4% paraformaldehyde (PFA)** | For 25 mL of solution |
| 4% PFA | 1 g of PFA powder |
| 1× PBS | 25 mL of 1× PBS |
| | Heat solution at 50–60 °C to dissolve powder |
| | Aliquot and store at −20 °C |

NOTE: *Avoid using calcium chloride and magnesium chloride in PBS, as this leads to increased autofluorescence in the samples.*

NOTE: *Handle pronase powder and PFA powder with extreme care, as they are hazardous materials.*

##### S2.2.3 HCR RNA-FISH

**Detection stage**

1. Transfer the required number of embryos for an experiment to a 2 mL Eppendorf tube.
2. Rehydrate with a series of graded 1 mL MeOH/PBST washes for 5 min each at room temperature:
   (a) 75% MeOH / 25% PBST
   (b) 50% MeOH / 50% PBST
   (c) 25% MeOH / 75% PBST
   (d) 5 × 100% PBST.
3. For each sample, move 5–8 embryos to a 1.5 mL Eppendorf tube and remove excess liquid.
4. Add 350 $\mu$L of pre-heated 30% probe hybridization buffer and incubate for 30 min at 37 °C.
   CAUTION: *Probe hybridization buffer contains formamide, a hazardous material.*
   NOTE: *Probe hybridization buffer should be pre-heated to 37 °C before use.*
5. Prepare probe solution by adding 4 $\mu$L of each 2 $\mu$M odd and even probe set to probe hybridization buffer and mixing well. Use a volume of probe hybridization buffer such that the final volume is 500 $\mu$L.
6. Remove the probe hybridization buffer from the samples and add the probe solution.
7. Incubate embryos overnight (>12 h) at 37 °C.
8. Remove excess probes by washing embryos 4 × 15 min with 500 $\mu$L of 30% probe wash buffer at 37 °C.
   CAUTION: *Probe wash buffer contains formamide, a hazardous material.*
   NOTE: *Probe wash buffer should be pre-heated to 37 °C before use.*
9. Wash embryos 3 × 5 min with 500 $\mu$L of 5× SSCT at room temperature.

**Amplification stage**

1. Add 350 $\mu$L of amplification buffer and incubate for 30 min at room temperature.
   NOTE: *Bring amplification buffer to room temperature before use.*
2. Separately prepare 30 pmol of hairpin h1 and 30 pmol of hairpin h2 by snap cooling 10 $\mu$L of 3 $\mu$M hairpin stock solution (heat at 95 °C for 90 seconds and cool to room temperature in a dark drawer for 30 min).
   NOTE: *HCR hairpins h1 and h2 are provided in hairpin storage buffer and are ready for snap cooling. HCR hairpins h1 and h2 should be snap cooled in separate tubes.*
3. Prepare hairpin solution by adding 10 $\mu$L of each snap-cooled hairpin to amplification buffer and mixing well. Use a volume of amplification buffer such that the final volume is 500 $\mu$L.
4. Remove the amplification buffer from the samples and add the hairpin solution.
5. Incubate embryos overnight (>12 h) in the dark at room temperature.
6. Remove excess hairpins by washing with 500 $\mu$L of 5× SSCT at room temperature:
   (a) 2 × 5 min
   (b) 2 × 30 min
   (c) 1 × 5 min

**Sample mounting for microscopy**

1. Transfer embryos onto a No. 1.5 coverslip and remove excess liquid.
2. Add 60 $\mu$L ProLong Glass Antifade Mountant on top of the embryos.
3. Use an eyelash tool to gently position the embryos in a lateral position for imaging.
4. Place the coverslip on a 37 °C surface (such as a slide moat) for 1 h to set the mountant.

##### S2.2.4 Buffer recipes for HCR RNA-FISH

| **PBST** | For 500 mL of solution |
|---|---|
| 1× PBS | 50 mL of 10× PBS |
| 0.1% Tween-20 | 5 mL of 10% Tween-20 |
| | Fill up to 500 mL with ultrapure $H_2O$ |
| | Filter with a 0.2 $\mu$m Nalgene Rapid-Flow filter |

| **5× SSCT** | For 500 mL of solution |
|---|---|
| 5× saline sodium citrate (SSC) | 125 mL of 20× SSC |
| 0.1% Tween-20 | 5 mL of 10% Tween-20 |
| | Fill up to 500 mL with ultrapure $H_2O$ |
| | Filter with a 0.2 $\mu$m Nalgene Rapid-Flow filter |

NOTE: *Avoid using calcium chloride and magnesium chloride in PBS, as this leads to increased autofluorescence in the samples.*

##### S2.2.5 Reagents and supplies

Pronase (Roche, 10165921001)
Paraformaldehyde (PFA) (Sigma-Aldrich, P6148)
10× Phosphate-buffered saline (PBS) (Invitrogen, AM9625)
Methanol (Mallinckrodt Chemicals, 3016-16)
20× Saline sodium citrate (SSC) (Life Technologies, 15557-044)
10% Tween-20 solution (Teknova, T0025)
ProLong Glass Antifade Mountant (Invitrogen, P36984)

#### S2.3 Protocols for HCR RNA-FISH/IF in fresh-frozen mouse brain sections

Protocols for HCR RNA-FISH/IF in fresh-frozen mouse brain sections are adapted from (Schwarzkopf *et al.*, 2021).

##### S2.3.1 Preparation of fresh-frozen mouse brain tissue sections

1. Remove slide-mounted sections from the $-80$ °C freezer and place on dry ice.

2. One by one, remove a slide from the slide storage box, draw a hydrophobic barrier around the tissue section, and add 100 $\mu$L of ice-cold 4% formaldehyde.
   CAUTION: *Use formaldehyde with extreme care, as it is a hazardous material.*
   NOTE: *The formaldehyde solution should be pre-cooled on ice before use.*

3. Fix for 2 h at 4 °C in a humidified chamber.
   NOTE: *To prevent evaportation, a humidified chamber should be used for all future steps other than autofluorescence bleaching step.*

4. Remove the 4% formaldehyde solution and add 2 mL of 1$\times$ of phosphate-buffered saline (PBS) across the entire slide to remove the optimal cutting temperature (OCT) compound and excess fixative. Incubate for 5 min at room temperature.

5. Wash 2 $\times$ 5 min with 1 mL of 1$\times$ PBS.

6. Remove 1$\times$ PBS and dry around the tissue section with a Kimwipe, taking care to not allow the tissue section to dry.

7. Re-apply the hydrophobic barrier around the tissue section if needed.

8. Wash 1 $\times$ 5 min with 200 $\mu$L of 1$\times$ PBS.

9. Proceed to autofluorescence bleaching protocol to reduce autofluorescence.

##### S2.3.2 Autofluorescence bleaching protocol

1. Prepare bleaching solution fresh before use.
   CAUTION: *Keep bleaching solution uncapped inside a fume hood, as it produces gas.*

2. Add 200 $\mu$L of bleaching solution on top of the tissue.

3. Place slides under a 180 W LED light at 4 °C. Keep slides 15 cm away from the light source.
   CAUTION: *The LED light is extremely bright. Use the LED light in a covered area to avoid eye exposure.*

4. Expose the tissue to the LED light for 3 h.
   NOTE: *Turn off the light and check the slide every hour, re-applying additional fresh bleaching solution if necessary. Do not allow the slide to dry.*

5. Wash slide 4 × 10 min with 100 $\mu$L of PBST.

6. Proceed to HCR RNA-FISH/IF assay.

##### S2.3.3 Buffer recipes for autofluorescence bleaching protocol

| **Bleaching solution** | For 3 mL of solution |
|---|---|
| 1× PBS | 300 $\mu$L of 10× PBS |
| 24 mM sodium hydroxide (NaOH) | 72 $\mu$L of 1 N NaOH |
| 4.5% hydrogen peroxide ($H_2O_2$) | 450 $\mu$L of 30% $H_2O_2$ |
| | 2178 $\mu$L of nanopure $H_2O$ |

| **PBST** | For 500 mL of solution |
|---|---|
| 1× PBS | 50 mL of 10× PBS |
| 0.1% Tween-20 | 5 mL of 10% Tween-20 |
| | Fill up to 500 mL with ultrapure $H_2O$ |
| | Filter with a 0.2 $\mu$m Nalgene Rapid-Flow filter |

NOTE: *Avoid using calcium chloride and magnesium chloride in PBS, as this leads to increased autofluorescence in the samples.*

##### S2.3.4 Multiplexed HCR RNA-FISH/IF using unlabeled primary antibody probes and initiator-labeled secondary antibody probes for protein targets, split-initiator DNA probes for RNA targets, and simultaneous HCR signal amplification for all targets

**Protein detection stage**

1. Block tissue by applying 100 $\mu$L of antibody buffer on top of the tissue. Incubate for 1 h at room temperature.
2. Prepare the primary antibody solution by adding all primary antibodies to antibody buffer and mixing well. Use a volume of antibody buffer such that the final volume is 100 $\mu$L per section.
   NOTE: *Follow manufacturer's guidelines for primary antibody working concentration.*
3. Remove the blocking solution. Add the primary antibody solution and incubate overnight (>12 h) at 4 °C.
   NOTE: *Incubation may be optimized (e.g., 1–2 h at room temperature) depending on the antibodies used.*
4. Remove excess antibodies by washing 3 × 5 min with 100 $\mu$L of PBST at room temperature.
5. Prepare the initiator-labeled secondary antibody solution by adding all secondary antibodies to antibody buffer and mixing well. Use a volume of antibody buffer such that the final volume is 100 $\mu$L per section.
   NOTE: *Use a working concentration of 1 $\mu$g/mL for all initiator-labeled secondary antibodies.*
6. Add the secondary antibody solution and incubate for 1 h at room temperature.
7. Remove excess antibodies by washing 3 × 5 min with 100 $\mu$L of PBST at room temperature.

**RNA detection stage**

1. Post-fix sample by adding 100 $\mu$L of 4% formaldehyde on the tissue section and incubating for 10 min at room temperature.
   CAUTION: *Use formaldehyde with extreme care, as it is a hazardous material.*
2. Wash 3 × 5 min with 100 $\mu$L of PBST.
3. Wash 1 × 5 min with 100 $\mu$L of 5× SSCT.
4. Add 100 $\mu$L of pre-heated 30% probe hybridization buffer and incubate for 30 min at 37 °C.
   CAUTION: *Probe hybridization buffer contains formamide, a hazardous material.*
   NOTE: *Probe hybridization buffer should be pre-heated to 37 °C before use.*
5. Prepare probe solution by adding 0.8 $\mu$L of each 2 $\mu$M odd and even probe set to probe hybridization buffer and mixing well. Use a volume of probe hybridization buffer such that the final volume is 100 $\mu$L.
6. Remove the probe hybridization buffer and add the probe solution on top of the samples.
7. Incubate overnight (>12 h) at 37 °C.
8. Remove excess probes by washing 4 × 15 min with 100 $\mu$L of 30% probe wash buffer at 37 °C.
   CAUTION: *Probe wash buffer contains formamide, a hazardous material.*
   NOTE: *Probe wash buffer should be pre-heated to 37 °C before use.*
9. Wash 3 × 5 min with 100 $\mu$L of 5× SSCT at room temperature.

**Amplification stage**

1. Add 100 $\mu$L of amplification buffer and incubate for 30 min at room temperature.
   NOTE: *Bring amplification buffer to room temperature before use.*

2. Separately prepare 6 pmol of hairpin h1 and 6 pmol of hairpin h2 by snap cooling 2 $\mu$L of each 3 $\mu$M hairpin stock solution (heat at 95 °C for 90 seconds and cool to room temperature in a dark drawer for 30 min).
   NOTE: *HCR hairpins h1 and h2 are provided in hairpin storage buffer and are ready for snap cooling. HCR hairpins h1 and h2 should be snap cooled in separate tubes.*

3. Prepare hairpin solution by adding 2 $\mu$L of each snap-cooled hairpin to amplification buffer and mixing well. Use a volume of amplification buffer such that the final volume is 100 $\mu$L.

4. Remove the amplification buffer and add the hairpin solution on top of the samples.

5. Incubate overnight (>12 h) in the dark at room temperature.

6. Remove excess hairpins by washing with 100 $\mu$L of 5× SSCT at room temperature:

   (a) 2 × 5 min

   (b) 2 × 15 min

   (c) 1 × 5 min

**Sample mounting for microscopy**

1. Aspirate 5× SSCT and carefully dry around the tissue section with a Kimwipe.
   NOTE: *Do not let the tissue section dry.*

2. Apply 60 $\mu$L of Fluoromount-G mountant on top of the tissue.

3. Slowly lower a 22 × 30 mm No. 1.5 coverslip on top of the mountant.

4. Store at 4 °C protected from light prior to imaging.

##### S2.3.5 Buffers for HCR IF with HCR RNA-FISH

| | |
|---|---|
| **<u>PBST</u>** | <u>For 500 mL of solution</u> |
| 1× PBS | 50 mL of 10× PBS |
| 0.1% Tween-20 | 5 mL of 10% Tween-20 |
| | Fill up to 500 mL with ultrapure $H_2O$ |
| | Filter with a 0.2 $\mu$m Nalgene Rapid-Flow filter |

| | |
|---|---|
| **<u>5× SSCT</u>** | <u>For 500 mL of solution</u> |
| 5× saline sodium citrate (SSC) | 125 mL of 20× SSC |
| 0.1% Tween-20 | 5 mL of 10% Tween-20 |
| | Fill up to 500 mL with ultrapure $H_2O$ |
| | Filter with a 0.2 $\mu$m Nalgene Rapid-Flow filter |

NOTE: *Avoid using calcium chloride and magnesium chloride in PBS, as this leads to increased autofluorescence in the samples.*

##### S2.3.6 Reagents and supplies

Image-iT 4% formaldehyde fixative solution in phosphate-buffered saline (PBS) (Invitrogen, FB002)
10× phosphate-buffered saline (PBS) (Invitrogen, AM9624)
30% hydrogen peroxide ($H_2O_2$) (Sigma-Aldrich, H1009)
1 N sodium hydroxide (NaOH) (Sigma-Aldrich, S2770)
7-band 2.1 180 Watt LED Grow Light (HTG Supply, LED-7BV2.1180)
10% Tween-20 solution (Teknova, T0025)
20× saline sodium citrate (SSC) (Life Technologies, 15557-044)
Fluoromount-G (SouthernBiotech, 0100-01)

### S3 Additional studies

#### S3.1 Summary of signal-to-background estimates for HCR RNA-FISH and HCR IF

| Method | Sample | Target | Type | Probes | Amplifier | SIG/BACK | Figures | Table |
|---|---|---|---|---|---|---|---|---|
| HCR RNA-FISH | whole-mount zebrafish embryo | *hbae1* | RNA | 3 DNA split-initiator pairs | B1-Alexa405 | $54 \pm 8$ | 2C, S2–S3 | S8 |
| HCR RNA-FISH | whole-mount zebrafish embryo | *mylpfa* | RNA | 13 DNA split-initiator pairs | B2-Atto425 | $62 \pm 5$ | 2C, S2–S3 | S8 |
| HCR RNA-FISH | whole-mount zebrafish embryo | *gfap* | RNA | 17 DNA split-initiator pairs | B6-Alexa488 | $19 \pm 2$ | 2C, S2–S3 | S8 |
| HCR RNA-FISH | whole-mount zebrafish embryo | *kdrl* | RNA | 60 DNA split-initiator pairs | B9-Alexa514 | $58 \pm 7$ | 2C, S2–S3 | S8 |
| HCR RNA-FISH | whole-mount zebrafish embryo | *shha* | RNA | 20 DNA split-initiator pairs | B7-Alexa546 | $39 \pm 6$ | 2C, S2–S3 | S8 |
| HCR RNA-FISH | whole-mount zebrafish embryo | *elavl3* | RNA | 20 DNA split-initiator pairs | B10-Alexa594 | $33 \pm 5$ | 2C, S2–S3 | S8 |
| HCR RNA-FISH | whole-mount zebrafish embryo | *sox10* | RNA | 34 DNA split-initiator pairs | B8-Atto633 | $70 \pm 20$ | 2C, S2–S3 | S8 |
| HCR RNA-FISH | whole-mount zebrafish embryo | *ntla* | RNA | 20 DNA split-initiator pairs | B3-Alexa700 | $17 \pm 3$ | 2C, S2–S3 | S8 |
| HCR RNA-FISH | whole-mount zebrafish embryo | *dmd* | RNA | 20 DNA split-initiator pairs | B5-Alexa750 | $18 \pm 3$ | 2C, S2–S3 | S8 |
| HCR RNA-FISH | whole-mount zebrafish embryo | *col2a1a* | RNA | 15 DNA split-initiator pairs | B4-iFluor800 | $100 \pm 20$ | 2C, S2–S3 | S8 |
| HCR RNA-FISH | whole-mount zebrafish embryo | *col2a1a* | RNA | 10 DNA split-initiator pairs | B1-Alexa405 | $80 \pm 20$ | 3B, S6, S11, S16 | S11 |
| HCR RNA-FISH | whole-mount zebrafish embryo | *col2a1a* | RNA | 10 DNA split-initiator pairs | B4-iFluor800 | $100 \pm 20$ | 3B, S6, S11, S16 | S11 |
| HCR RNA-FISH | whole-mount zebrafish embryo | *mylpfa* | RNA | 6 DNA split-initiator pairs | B2-Atto425 | $35 \pm 7$ | 3B, S7, S12, S16 | S11 |
| HCR RNA-FISH | whole-mount zebrafish embryo | *mylpfa* | RNA | 6 DNA split-initiator pairs | B3-Alexa700 | $130 \pm 20$ | 3B, S7, S12, S16 | S11 |
| HCR RNA-FISH | whole-mount zebrafish embryo | *elavl3* | RNA | 20 DNA split-initiator pairs | B6-Alexa488 | $47 \pm 9$ | 3B, S8, S13, S16 | S11 |
| HCR RNA-FISH | whole-mount zebrafish embryo | *elavl3* | RNA | 20 DNA split-initiator pairs | B10-Alexa594 | $70 \pm 20$ | 3B, S8, S13, S16 | S11 |
| HCR RNA-FISH | whole-mount zebrafish embryo | *kdrl* | RNA | 30 DNA split-initiator pairs | B9-Alexa514 | $50 \pm 8$ | 3B, S9, S14, S16 | S11 |
| HCR RNA-FISH | whole-mount zebrafish embryo | *kdrl* | RNA | 30 DNA split-initiator pairs | B8-Atto633 | $110 \pm 20$ | 3B, S9, S14, S16 | S11 |
| HCR RNA-FISH | whole-mount zebrafish embryo | *dmd* | RNA | 20 DNA split-initiator pairs | B7-Alexa546 | $40 \pm 6$ | 3B, S10, S15, S16 | S11 |
| HCR RNA-FISH | whole-mount zebrafish embryo | *dmd* | RNA | 20 DNA split-initiator pairs | B5-Alexa750 | $27 \pm 6$ | 3B, S10, S15, S16 | S11 |
| HCR IF | fresh-frozen mouse brain section | NFH | protein | $1^\circ$pAb + $2^\circ$pAb-initiator | B1-Alexa405 | $26 \pm 7$ | 5C, S20–S23 | S13 |
| HCR IF | fresh-frozen mouse brain section | CD31 | protein | $1^\circ$mAb + $2^\circ$pAb-initiator | B2-Atto425 | $55 \pm 9$ | 5C, S20–S23 | S13 |
| HCR RNA-FISH | fresh-frozen mouse brain section | *Slc17a7* | RNA | 20 DNA split-initiator pairs | B6-Alexa488 | $25 \pm 4$ | 5C, S20–S23 | S13 |
| HCR RNA-FISH | fresh-frozen mouse brain section | *Gad1* | RNA | 33 DNA split-initiator pairs | B9-Alexa514 | $110 \pm 20$ | 5C, S20–S23 | S13 |
| HCR RNA-FISH | fresh-frozen mouse brain section | *Sst* | RNA | 11 DNA split-initiator pairs | B7-Alexa546 | $140 \pm 20$ | 5C, S20–S23 | S13 |
| HCR RNA-FISH | fresh-frozen mouse brain section | *Actb* | RNA | 20 DNA split-initiator pairs | B5-Alexa594 | $34 \pm 4$ | 5C, S20–S23 | S13 |
| HCR RNA-FISH | fresh-frozen mouse brain section | *Lamp5* | RNA | 27 DNA split-initiator pairs | B8-Atto633 | $30 \pm 9$ | 5C, S20–S23 | S13 |
| HCR RNA-FISH | fresh-frozen mouse brain section | *Plp1* | RNA | 24 DNA split-initiator pairs | B3-Alexa700 | $100 \pm 20$ | 5C, S20–S23 | S13 |
| HCR RNA-FISH | fresh-frozen mouse brain section | *Vip* | RNA | 24 DNA split-initiator pairs | B10-Alexa750 | $80 \pm 20$ | 5C, S20–S23 | S13 |
| HCR IF | fresh-frozen mouse brain section | RBFOX3 | protein | $1^\circ$mAb + $2^\circ$pAb-initiator | B4-iFluor800 | $96 \pm 8$ | 5C, S20–S23 | S13 |

**Table S7. Signal-to-background summary for HCR RNA-FISH and HCR IF.** Mean $\pm$ SEM for representative regions within $N = 3$ embryos (27 hpf whole-mount zebrafish embryos) or $N = 3$ replicate sections (5 $\mu$m fresh-frozen coronal mouse brain sections).

#### S3.2 Reference spectra, replicates, signal, and background for 10-plex RNA imaging with high signal-to-background in whole-mount zebrafish embryos (cf. Figure 2)

Additional studies are presented as follows:

- Figure S1 displays 11 raw and normalized reference spectra (1 per fluorophore and 1 for autofluorescence).
- Figure S2 displays 10-plex images for $N = 3$ replicate embryos linearly unmixed using the Leica LAS X software (cf. Figure 2C).
- Figure S3 displays representative regions of individual channels used for measurement of signal and background for each target.
- Table S8 displays estimated values for signal, background, and signal-to-background for each target.
- Figure S4 displays 10-plex images for $N = 3$ replicate embryos linearly unmixed using the Unmix 1.0 software package (cf. Figure S2), providing an alternative for researchers that do not have access to the LAS X software.
- Figure S5 compares the pixel intensities for linear unmixing using the Leica LAS X software vs the Unmix 1.0 software package. For each channel and replicate, the pixel intensities are highly correlated between the two methods (Pearson correlation coefficient $r > 0.996$ for each fluorophore and for autofluorescence).

**Protocol:** spectral imaging and linear unmixing protocol of Section S2.1 using the HCR RNA-FISH protocol of Section S2.2.
**Sample:** whole-mount zebrafish embryos; fixed 27 hpf.
**Reagents:** Table S1.
**Microscope settings:** Table S3.

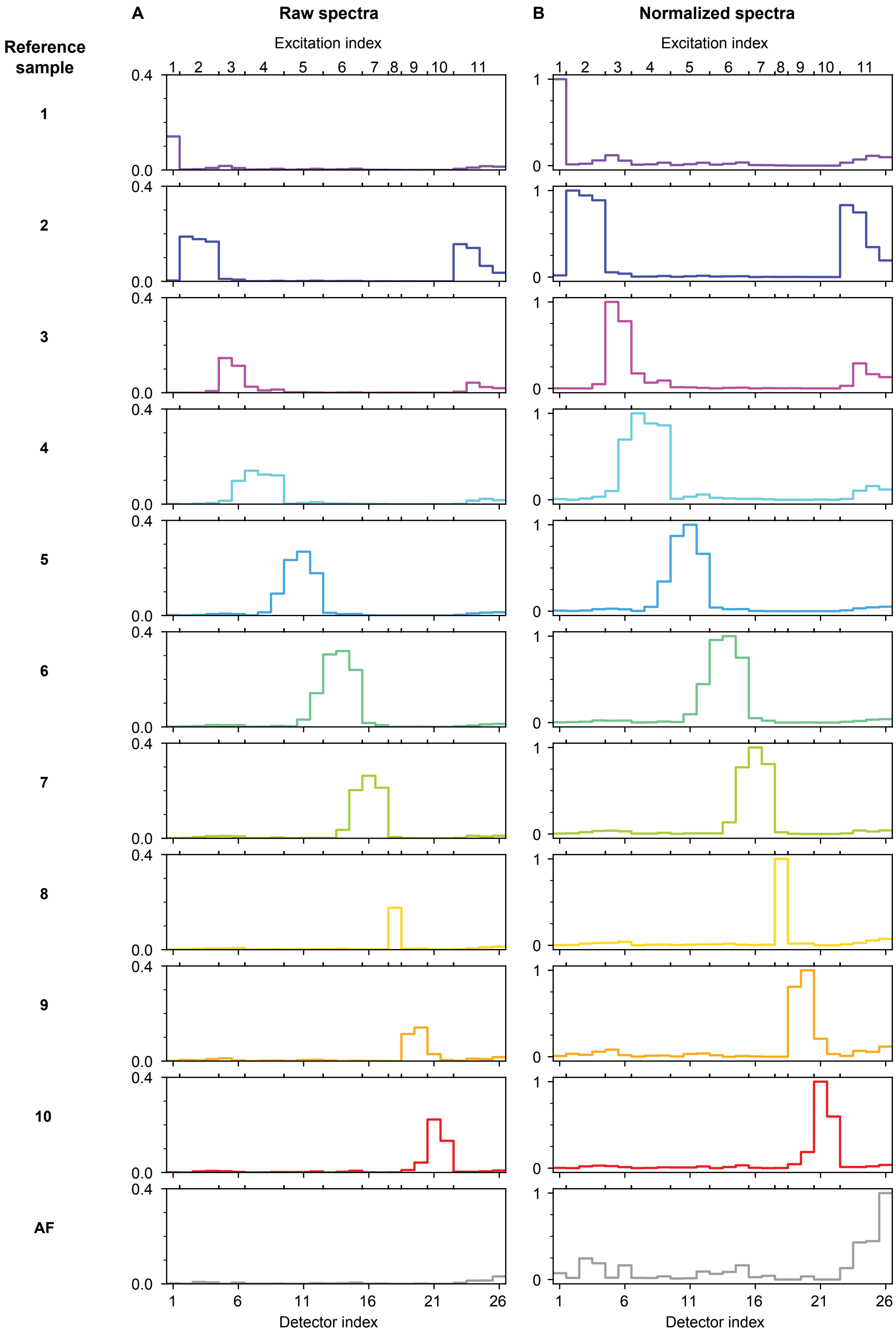

**Figure S1. 11 reference spectra in whole-mount zebrafish embryos (1 per fluorophore and 1 for autofluorescence (AF); cf. Figure 1C).** (A) Raw reference spectra. (B) Normalized reference spectra (each spectrum scaled to have a maximum value of 1). Protocol: Section S2.1. Sample: whole-mount zebrafish embryos; fixed 27 hpf.

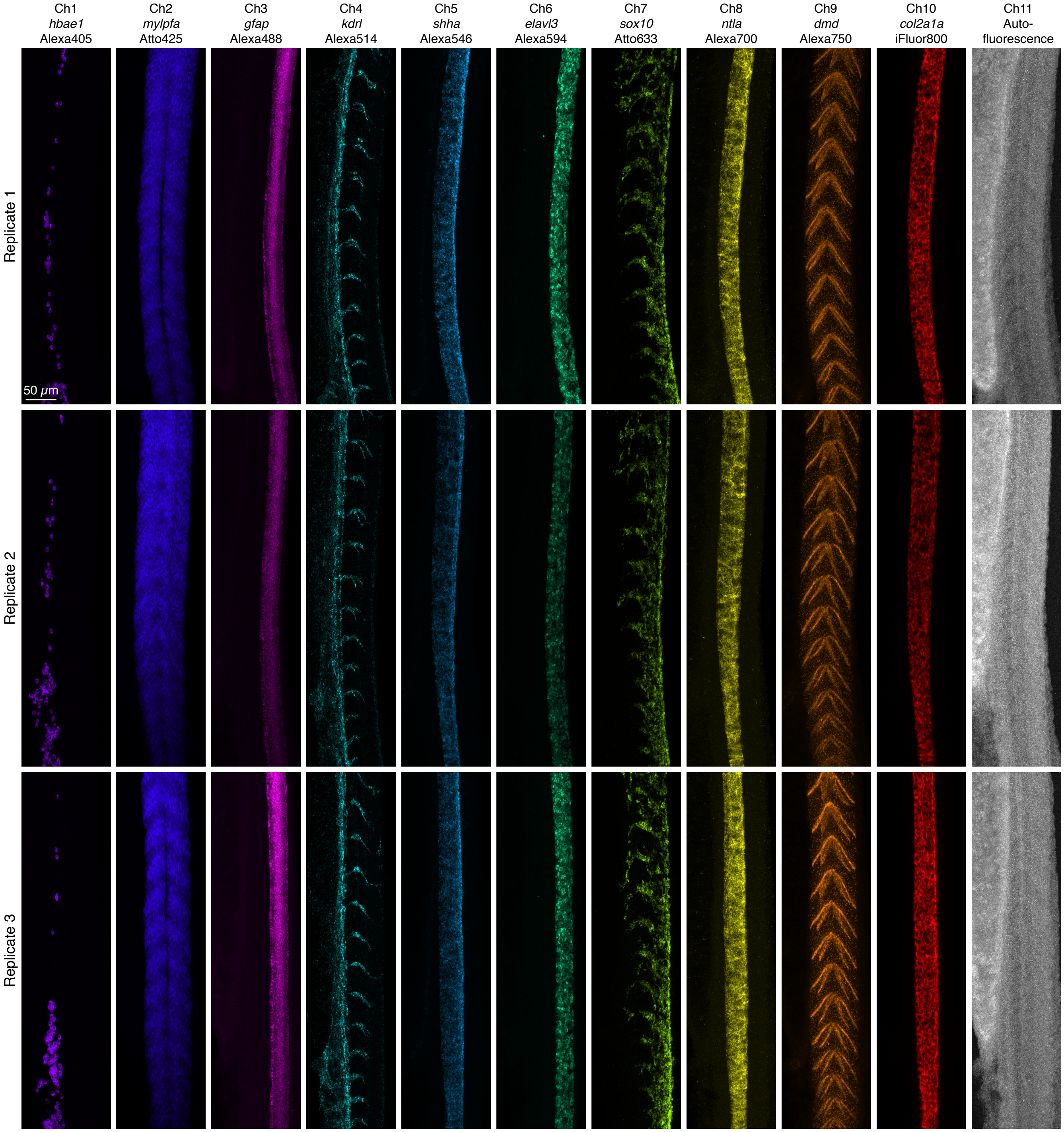

**Figure S2. Replicates for 10-plex RNA imaging using HCR RNA-FISH in whole-mount zebrafish embryos (cf. Figure 2C).** Linearly unmixed fluorescence channels using Leica LAS X software. Maximum-intensity z-projection for each channel; 0.57×0.57×4.0 μm pixels. Replicate 1: 64 optical sections (88.4 μm total). Replicate 2: 55 optical sections (76.3 μm total). Replicate 3: 77 optical sections (105.8 μm total). Ch1: *hbae1* (Alexa405). Ch2: *mylpfa* (Atto425). Ch3: *gfap* (Alexa488). Ch4: *kdrl* (Alexa514). Ch5: *shha* (Alexa546). Ch6: *elavl3* (Alexa594). Ch7: *sox10* (Atto633). Ch8: *ntla* (Alexa700). Ch9: *dmd* (Alexa750). Ch10: *col2a1a* (iFluor800). Sample: whole-mount zebrafish embryos; fixed 27 hpf.

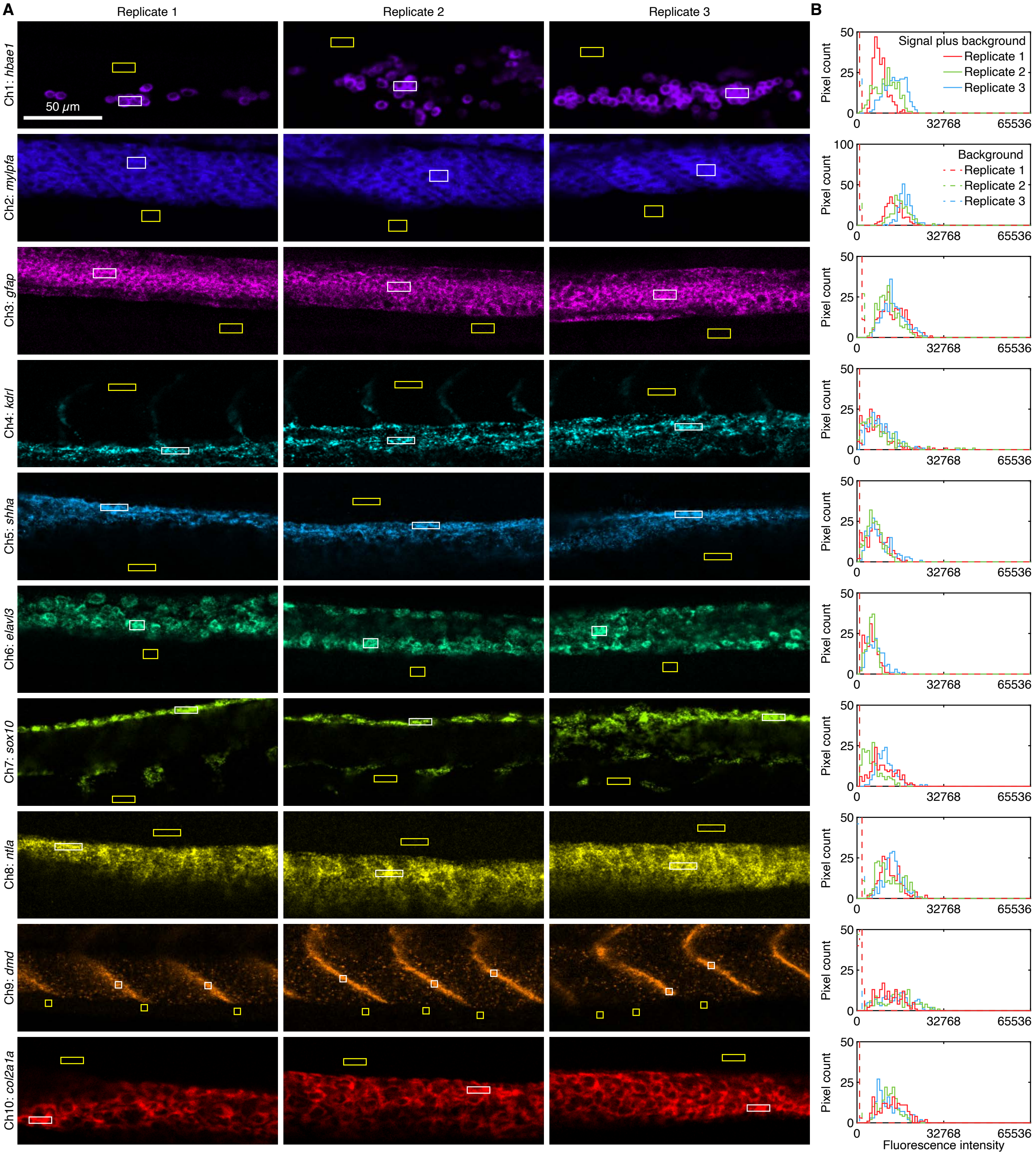

**Figure S3. Measurement of signal and background for 10-plex RNA imaging using HCR RNA-FISH in whole-mount zebrafish embryos (cf. Figure 2C).** (A) Linearly unmixed fluorescence channels using Leica LAS X software. For each of three replicate embryos, a representative single optical section was selected for each channel based on the expression of the corresponding target RNA. (B) Pixel intensity histograms for signal plus background (pixels within solid white boundary in panel A) and background (pixels within solid yellow boundary in panel A). Confocal images collected with the microscope laser intensity and detector gain optimized to avoid saturating SIG + BACK pixels. Ch1: *hbae1* (Alexa405). Ch2: *mylpfa* (Atto425). Ch3: *gfap* (Alexa488). Ch4: *kdrl* (Alexa514). Ch5: *shha* (Alexa546). Ch6: *elavl3* (Alexa594). Ch7: *sox10* (Atto633). Ch8: *ntla* (Alexa700). Ch9: *dmd* (Alexa750). Ch10: *col2a1a* (iFluor800). Sample: whole-mount zebrafish embryos; fixed 27 hpf.

| Target RNA | SIG+BACK | SIG | BACK | SIG/BACK |
|---|---|---|---|---|
| *hbae1* | $11\,000 \pm 2000$ | $11\,000 \pm 2000$ | $197 \pm 5$ | $54 \pm 8$ |
| *mylpfa* | $16\,000 \pm 1000$ | $16\,000 \pm 1000$ | $249 \pm 5$ | $62 \pm 5$ |
| *gfap* | $12\,800 \pm 700$ | $12\,100 \pm 700$ | $650 \pm 50$ | $19 \pm 2$ |
| *kdrl* | $7800 \pm 400$ | $7600 \pm 400$ | $130 \pm 10$ | $58 \pm 7$ |
| *shha* | $6900 \pm 500$ | $6700 \pm 500$ | $170 \pm 20$ | $39 \pm 6$ |
| *elavl3* | $5500 \pm 500$ | $5300 \pm 500$ | $160 \pm 20$ | $33 \pm 5$ |
| *sox10* | $8000 \pm 2000$ | $8000 \pm 2000$ | $110 \pm 10$ | $70 \pm 20$ |
| *ntla* | $12\,300 \pm 400$ | $11\,600 \pm 400$ | $700 \pm 100$ | $17 \pm 3$ |
| *dmd* | $14\,000 \pm 1000$ | $13\,000 \pm 1000$ | $700 \pm 80$ | $18 \pm 3$ |
| *col2a1a* | $12\,000 \pm 400$ | $11\,800 \pm 400$ | $120 \pm 20$ | $100 \pm 20$ |

**Table S8. Estimated signal-to-background for 10-plex RNA imaging using HCR RNA-FISH in whole-mount zebrafish embryos (cf. Figure 2C).** Mean $\pm$ estimated standard error of the mean via uncertainty propagation for $N = 3$ replicate embryos. Analysis based on rectangular regions depicted in Figure S3 using methods of Section S1.4.2.

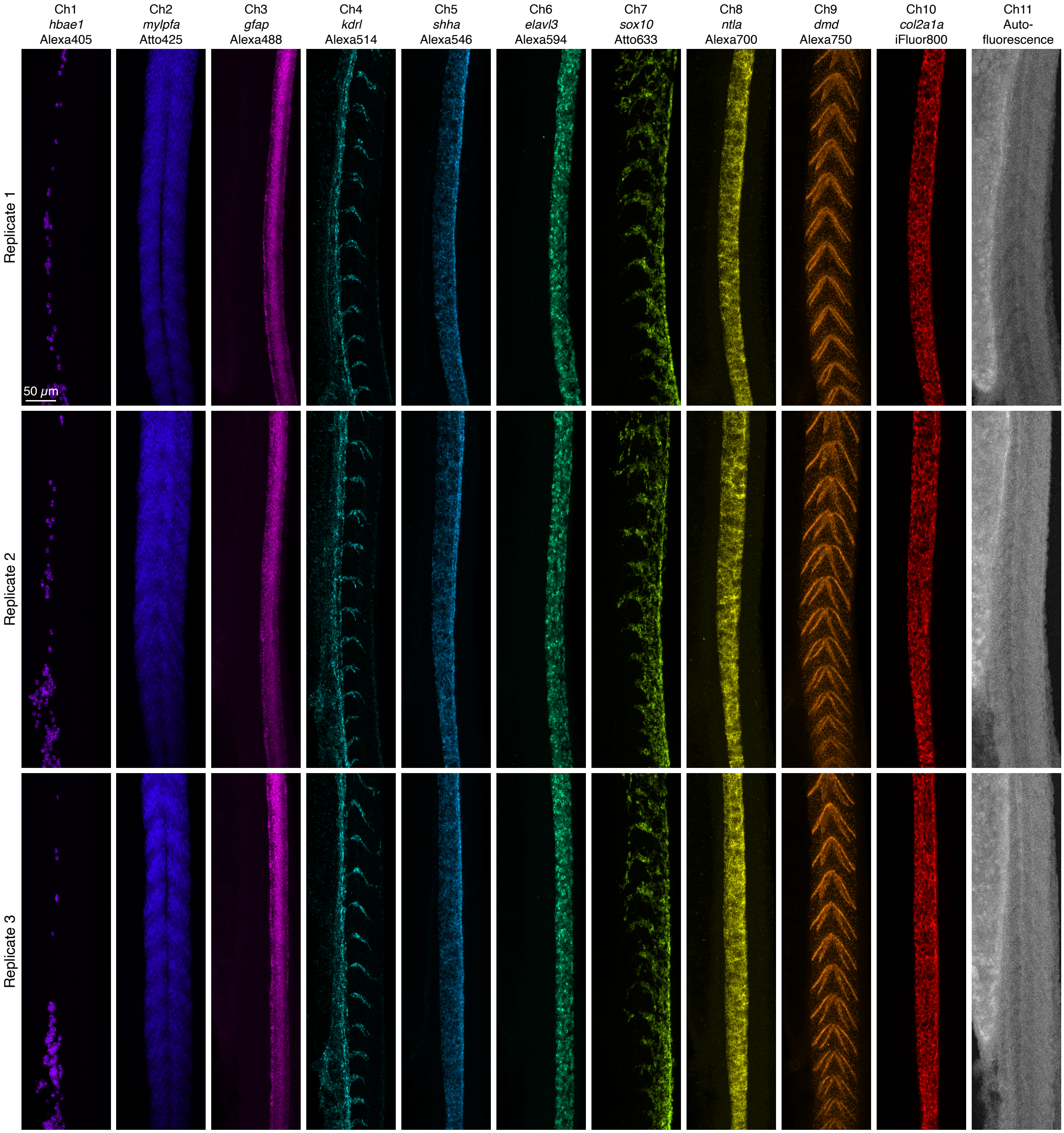

**Figure S4. Replicates for 10-plex RNA imaging using HCR RNA-FISH in whole-mount zebrafish embryos linearly unmixed using the Unmix 1.0 software package (cf. Figure S2 linearly unmixed using the Leica LAS X software).** Linearly unmixed fluorescence channels. Maximum-intensity z-projection for each channel; 0.57×0.57×4.0 μm pixels. Replicate 1: 64 optical sections (88.4 $\mu$m total). Replicate 2: 55 optical sections (76.3 $\mu$m total). Replicate 3: 77 optical sections (105.8 $\mu$m total). Ch1: *hbae1* (Alexa405). Ch2: *mylpfa* (Atto425). Ch3: *gfap* (Alexa488). Ch4: *kdrl* (Alexa514). Ch5: *shha* (Alexa546). Ch6: *elavl3* (Alexa594). Ch7: *sox10* (Atto633). Ch8: *ntla* (Alexa700). Ch9: *dmd* (Alexa750). Ch10: *col2a1a* (iFluor800). Sample: whole-mount zebrafish embryos; fixed 27 hpf.

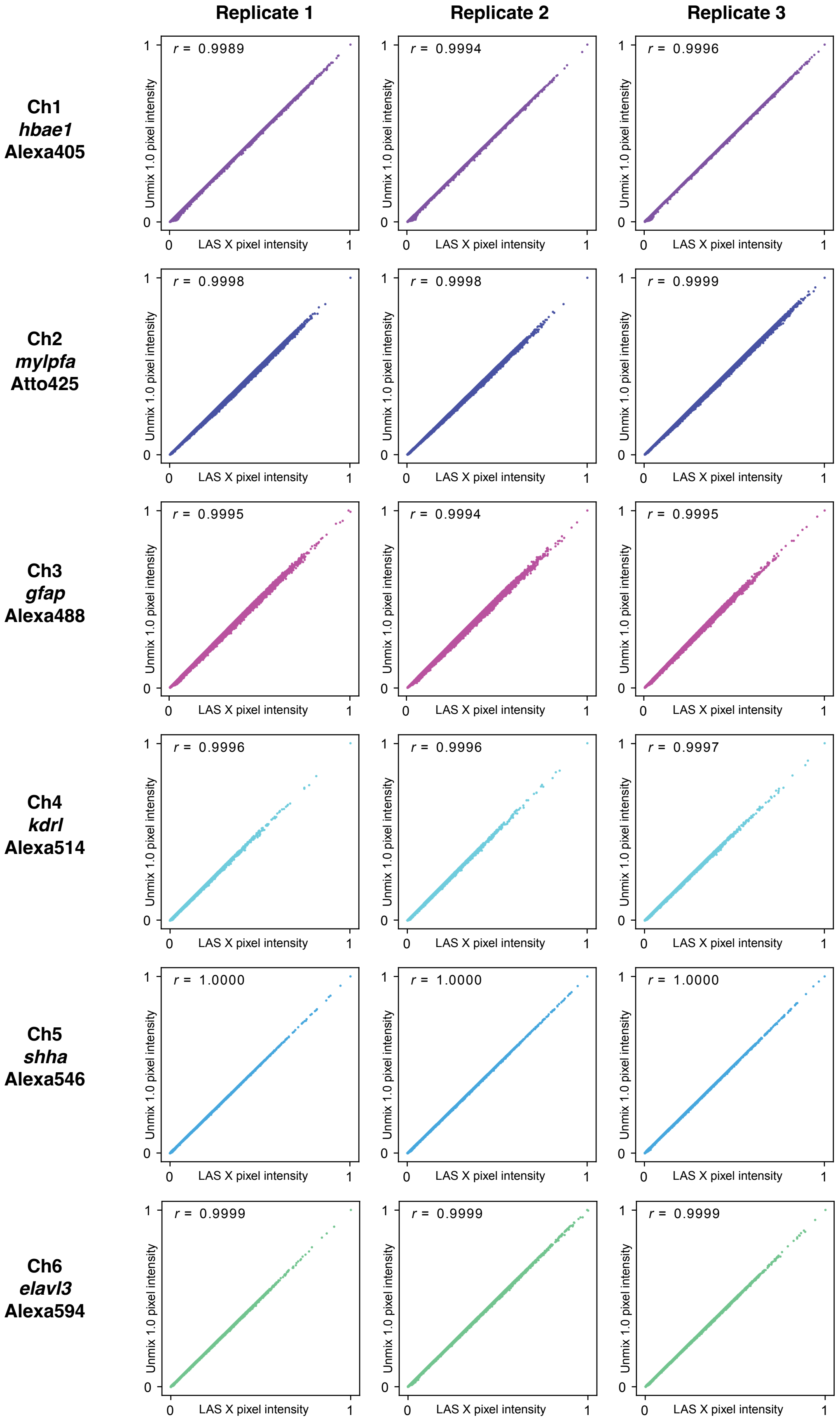
Replicate 1
Replicate 2
Replicate 3
Ch1
hbae1
Alexa405
r = 0.9989
r = 0.9994
r = 0.9996
Ch2
mylpfa
Atto425
r = 0.9998
r = 0.9998
r = 0.9999
Ch3
gfap
Alexa488
r = 0.9995
r = 0.9994
r = 0.9995
Ch4
kdrl
Alexa514
r = 0.9996
r = 0.9996
r = 0.9997
Ch5
shha
Alexa546
r = 1.0000
r = 1.0000
r = 1.0000
Ch6
elavl3
Alexa594
r = 0.9999
r = 0.9999
r = 0.9999
Unmix 1.0 pixel intensity
LAS X pixel intensity
0
1

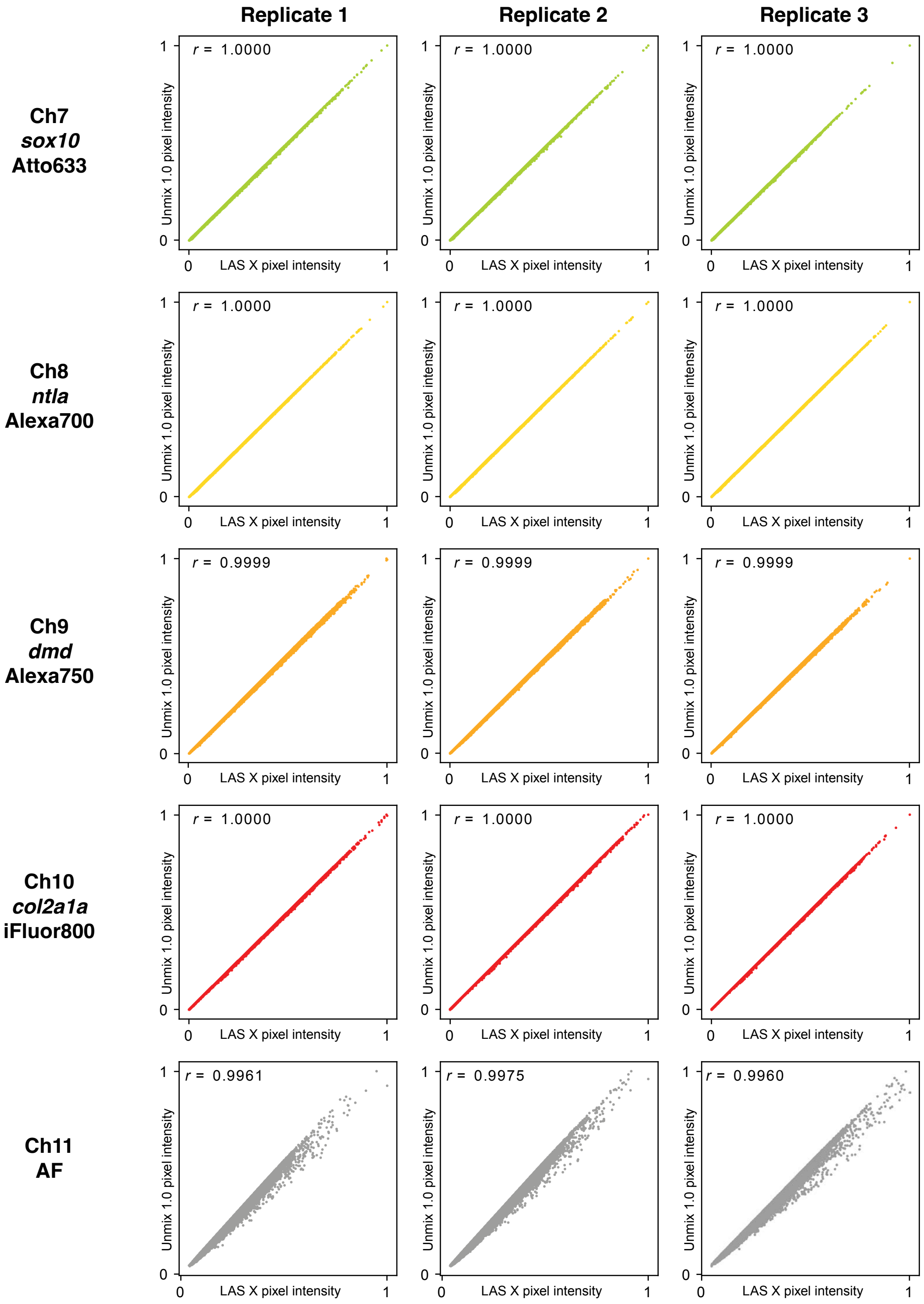

**Figure S5. Comparison of pixel intensities for linear unmixing using Leica LAS X software vs Unmix 1.0 software for 10-plex RNA imaging in whole-mount zebrafish embryos.** Pixel intensity scatter plots for replicate images of Figure S2 (Leica LAS X software) and Figure S4 (Unmix 1.0 software). Maximum-intensity z-projection for each channel; 0.57×0.57×4.0 μm pixels. Each scatter plot contains 262,144 dots corresponding to 1024×256 pixels. Pixel intensities normalized so that the maximum intensity for each axis is 1. Pearson correlation coefficient, $r$.

##### S3.3 10-plex qHCR imaging: RNA relative quantitation with subcellular resolution in an anatomical context (cf. Figure 3)

Additional studies are presented as follows:

- Figures S6–S10 display 2-channel images and 2-channel voxel intensity scatter plots for each of 5 target RNAs in $N = 3$ replicate embryos. To address chromatic aberration resulting from the wide range of wavelengths used in spectral imaging, Huygens Software was used to apply a chromatic aberration correction across the 10 channels.
- Table S9 displays values used for signal normalization in Figures S6–S10.
- Figures S11–S15 display 2-channel images and 2-channel voxel intensity scatter plots for each of 5 target RNAs in $N = 3$ replicate embryos without chromatic aberration correction (cf. Figures S6–S10). For a 2-channel redundant detection experiment, a scatter plot of voxel intensities compares two intensities for a given voxel (1 per channel) and that voxel is supposed to represent the identical physical volume within the sample for both channels. If there is a chromatic aberration across the 10 channels, a given voxel can represent different volumes within the sample for different channels, thus artificially degrading the correlation in the scatter plot. For example, for 2-channel redundant detection of *col2a1a* using Ch1 (Alexa405) and Ch10 (iFluor800), chromatic aberration due to the large wavelength gap between the two channels artificially reduces the correlation between voxel intensities (Figure S11B); correction of the chromatic aberration allows comparison of voxel intensities representing the same physical volume within the sample and significantly improves the correlation between voxel intensities for this pair of channels (Figure S6B).
- Table S10 displays values used for signal normalization in Figures S11–S15.
- Figure S16 display representative regions used for measurement of signal and background for the 10 channels. Images used for signal and background calculations were not corrected for chromatic aberration as each channel is analyzed separately.
- Table S11 displays estimated values for signal, background, and signal-to-background for 10 channels.

**Protocol:** Spectral imaging and linear unmixing protocol of Section S2.1 using the HCR RNA-FISH protocol of Section S2.2.
**Sample:** Whole-mount zebrafish embryos; fixed 27 hpf.
**Reagents:** Table S1.
**Microscope settings:** Table S3.

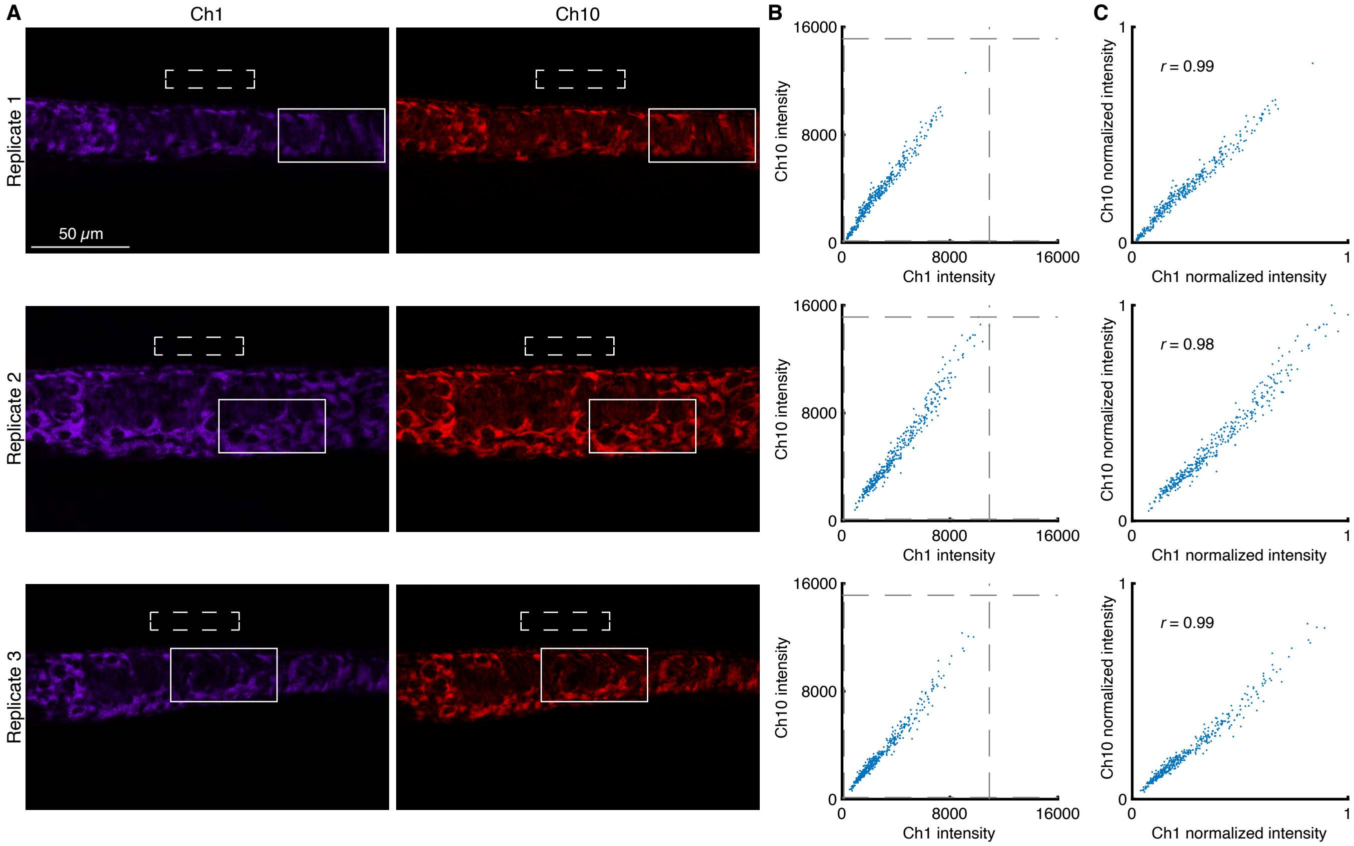

**Figure S6. 2-channel redundant detection of target mRNA *col2a1a* in the context of a 10-plex experiment in whole-mount zebrafish embryos (cf. Figure 3).** (A) Confocal images: individual channels; single optical section. Chromatic aberration correction applied with Huygens Software. Solid boundary denotes region of variable expression; dashed boundary denotes region of no/low expression. Pixel size: 0.180×0.180×1.2 μm. Sample: Whole-mount zebrafish embryos; fixed 27 hpf. (B) Raw voxel intensity scatter plots representing signal plus background for voxels within solid boundary of panel A. Voxel size: 2.0×2.0×1.2 μm. Dashed lines represent BOT and TOP values (Table S9) used to normalize data for panel C using methods of Section S1.4.3. (C) Normalized voxel intensity scatter plots representing estimated normalized signal (Pearson correlation coefficient, $r$). Ch1: *col2a1a* (Alexa405). Ch10: *col2a1a* (iFluor800). Sample: Whole-mount zebrafish embryos; fixed 27 hpf.

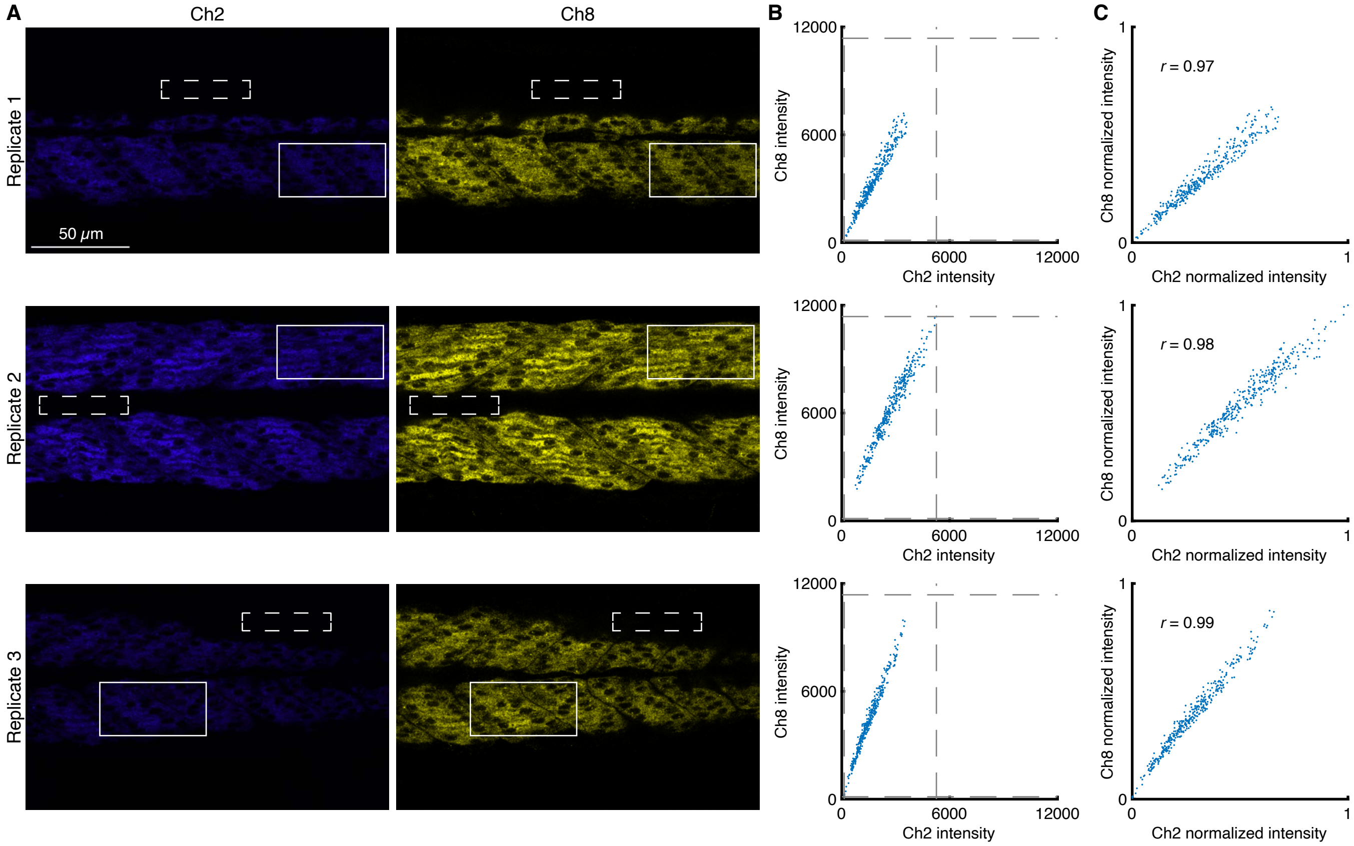

**Figure S7. 2-channel redundant detection of target mRNA *mylpfa* in the context of a 10-plex experiment in whole-mount zebrafish embryos (cf. Figure 3).** (A) Confocal images: individual channels; single optical section. Chromatic aberration correction applied with Huygens Software. Solid boundary denotes region of variable expression; dashed boundary denotes region of no/low expression. Pixel size: $0.180 \times 0.180 \times 1.2$ $\mu$m. Sample: Whole-mount zebrafish embryos; fixed 27 hpf. (B) Raw voxel intensity scatter plots representing signal plus background for voxels within solid boundary of panel A. Voxel size: $2.0 \times 2.0 \times 1.2$ $\mu$m. Dashed lines represent BOT and TOP values (Table S9) used to normalize data for panel C using methods of Section S1.4.3. (C) Normalized voxel intensity scatter plots representing estimated normalized signal (Pearson correlation coefficient, $r$). Ch2: *mylpfa* (Atto425). Ch8: *mylpfa* (Alexa700). Sample: Whole-mount zebrafish embryos; fixed 27 hpf.

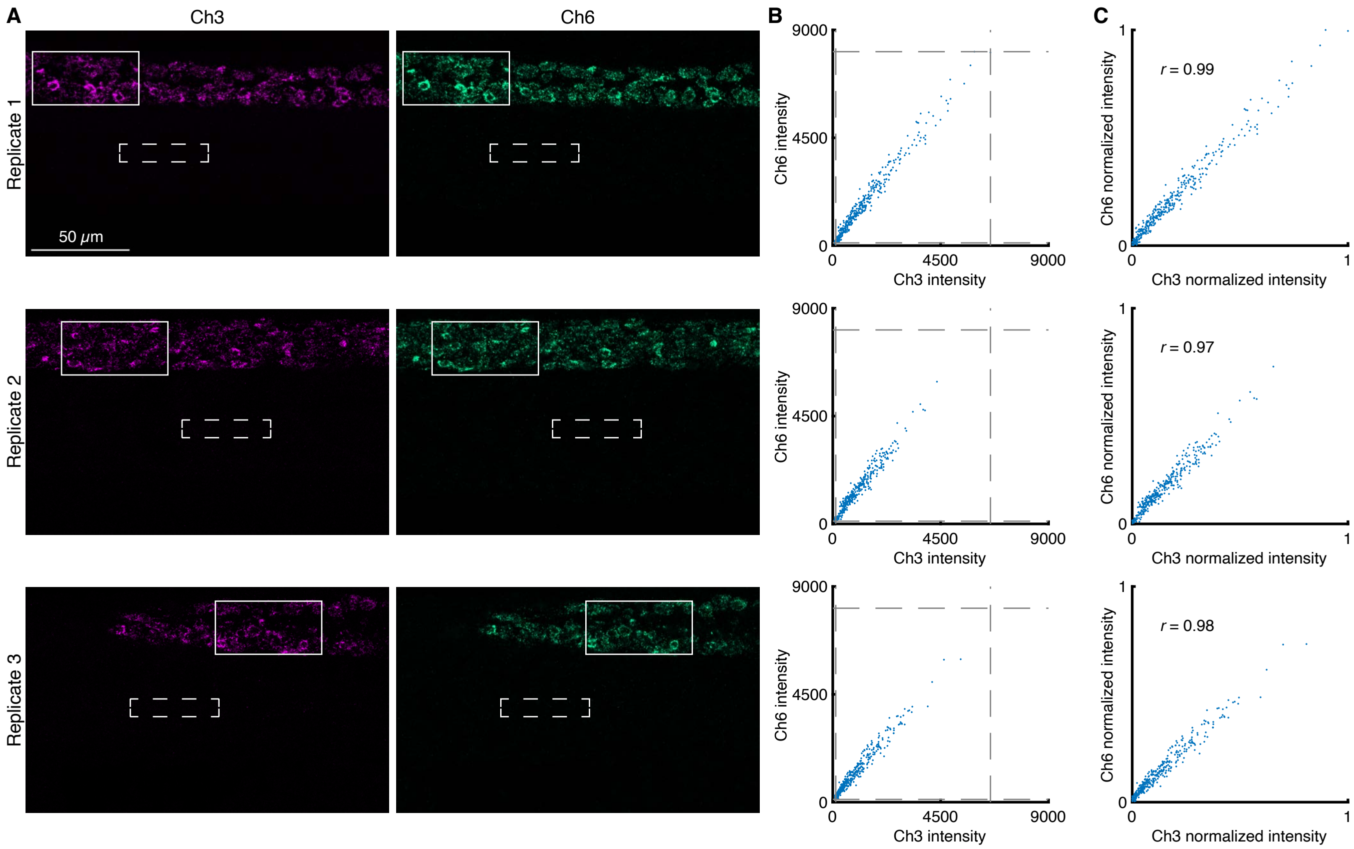

**Figure S8. 2-channel redundant detection of target mRNA *elavl3* in the context of a 10-plex experiment in whole-mount zebrafish embryos (cf. Figure 3).** (A) Confocal images: individual channels; single optical section. Chromatic aberration correction applied with Huygens Software. Solid boundary denotes region of variable expression; dashed boundary denotes region of no/low expression. Pixel size: $0.180 \times 0.180 \times 1.2$ $\mu$m. Sample: Whole-mount zebrafish embryos; fixed 27 hpf. (B) Raw voxel intensity scatter plots representing signal plus background for voxels within solid boundary of panel A. Voxel size: $2.0 \times 2.0 \times 1.2$ $\mu$m. Dashed lines represent BOT and TOP values (Table S9) used to normalize data for panel C using methods of Section S1.4.3. (C) Normalized voxel intensity scatter plots representing estimated normalized signal (Pearson correlation coefficient, $r$). Ch3: *elavl3* (Alexa488). Ch6: *elavl3* (Alexa594). Sample: Whole-mount zebrafish embryos; fixed 27 hpf.

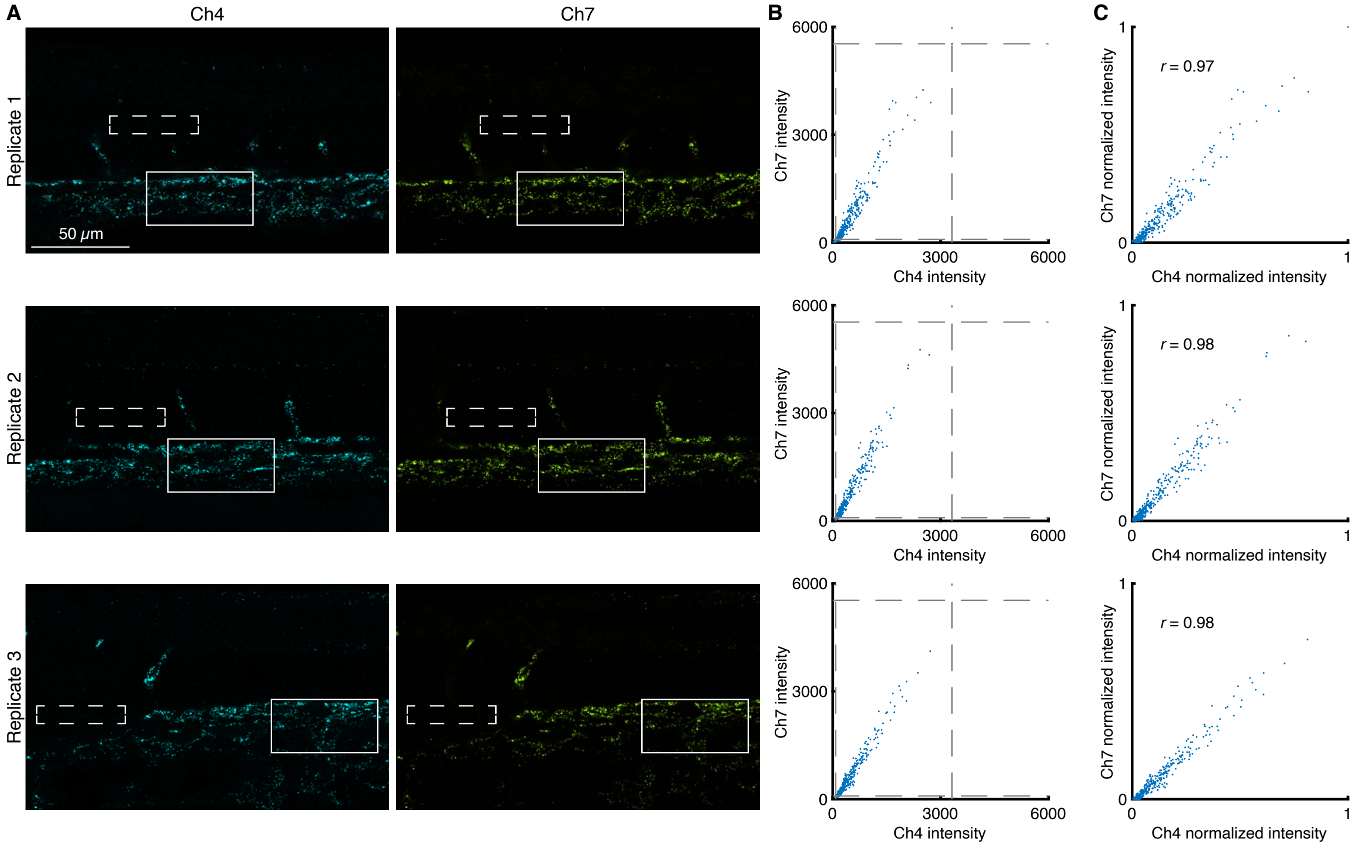

**Figure S9. 2-channel redundant detection of target mRNA *kdrl* in the context of a 10-plex experiment in whole-mount zebrafish embryos (cf. Figure 3).** (A) Confocal images: individual channels; single optical section. Chromatic aberration correction applied with Huygens Software. Solid boundary denotes region of variable expression; dashed boundary denotes region of no/low expression. Pixel size: $0.180\times0.180\times1.2$ $\mu$m Sample: Whole-mount zebrafish embryos; fixed 27 hpf. (B) Raw voxel intensity scatter plots representing signal plus background for voxels within solid boundary of panel A. Voxel size: $2.0\times2.0\times1.2$ $\mu$m. Dashed lines represent BOT and TOP values (Table S9) used to normalize data for panel C using methods of Section S1.4.3. (C) Normalized voxel intensity scatter plots representing estimated normalized signal (Pearson correlation coefficient, $r$). Ch4: *kdrl* (Alexa514). Ch7: *kdrl* (Atto633). Sample: Whole-mount zebrafish embryos; fixed 27 hpf.

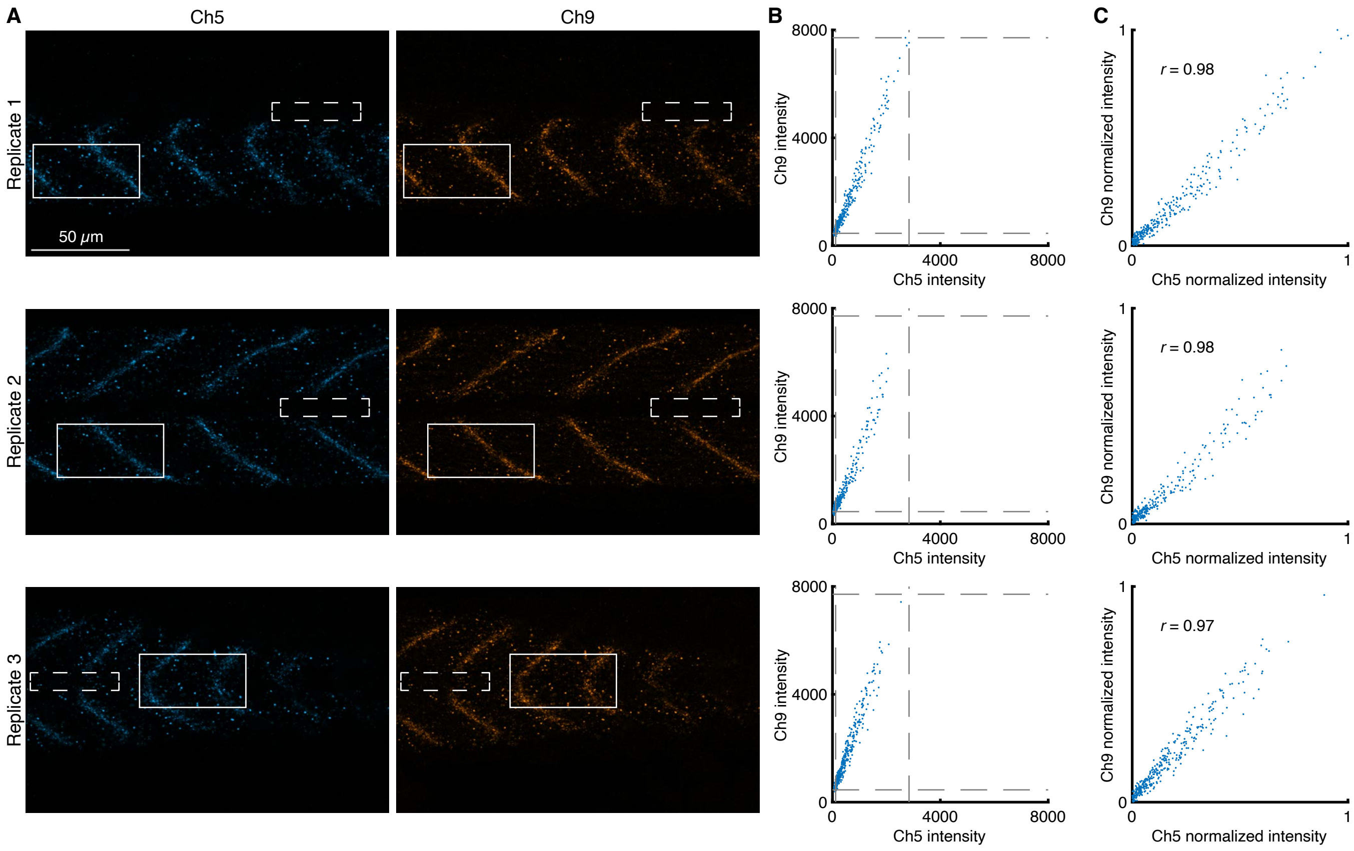

**Figure S10. 2-channel redundant detection of target mRNA *dmd* in the context of a 10-plex experiment in whole-mount zebrafish embryos (cf. Figure 3).** (A) Confocal images: individual channels; single optical section. Chromatic aberration correction applied with Huygens Software. Solid boundary denotes region of variable expression; dashed boundary denotes region of no/low expression. Pixel size: $0.180 \times 0.180 \times 1.2$ $\mu$m. Sample: Whole-mount zebrafish embryos; fixed 27 hpf. (B) Raw voxel intensity scatter plots representing signal plus background for voxels within solid boundary of panel A. Voxel size: $2.0 \times 2.0 \times 1.2$ $\mu$m. Dashed lines represent BOT and TOP values (Table S9) used to normalize data for panel C using methods of Section S1.4.3. (C) Normalized voxel intensity scatter plots representing estimated normalized signal (Pearson correlation coefficient, $r$). Ch5: *dmd* (Alexa546). Ch9: *dmd* (Alexa750). Sample: Whole-mount zebrafish embryos; fixed 27 hpf.

| Channel | Target RNA | Fluorophore | BOT | TOP |
|---|---|---|---|---|
| Ch1 | *col2a1a* | Alexa405 | 140 | 10923 |
| Ch2 | *mylpfa* | Atto425 | 136 | 5263 |
| Ch3 | *elavl3* | Alexa488 | 126 | 6577 |
| Ch4 | *kdrl* | Alexa514 | 83 | 3316 |
| Ch5 | *dmd* | Alexa546 | 70 | 2832 |
| Ch6 | *elavl3* | Alexa594 | 115 | 8094 |
| Ch7 | *kdrl* | Atto633 | 90 | 5532 |
| Ch8 | *mylpfa* | Alexa700 | 133 | 11371 |
| Ch9 | *dmd* | Alexa750 | 293 | 7750 |
| Ch10 | *col2a1a* | iFluor800 | 122 | 15125 |

**Table S9. BOT and TOP values used to calculate normalized voxel intensities for scatter plots of Figures 3C and S6C–S10C.** Analysis based on rectangular regions depicted in Figures S6A–S10A using the methods of Section S1.4.3.

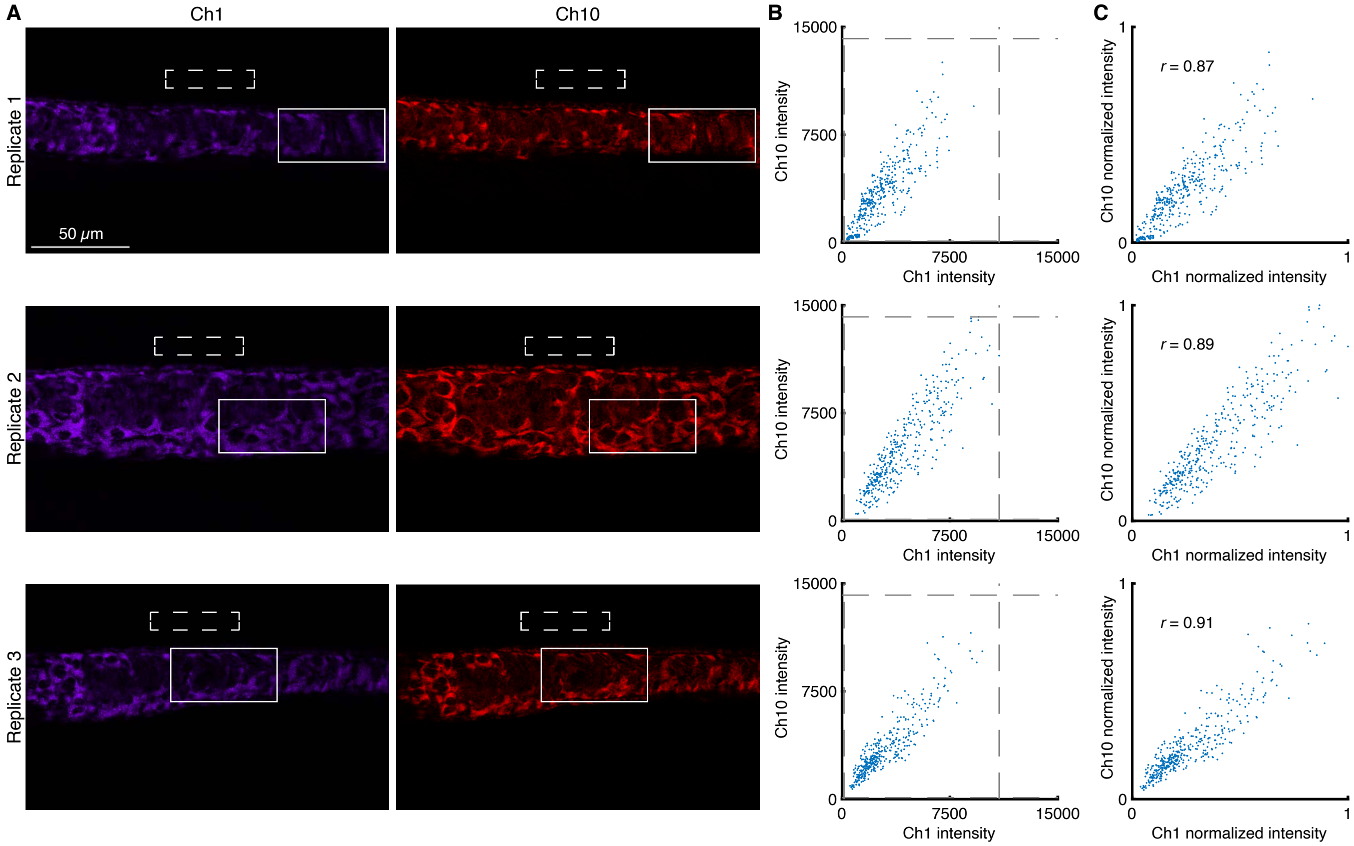

**Figure S11. 2-channel redundant detection of target mRNA *col2a1a* in the context of a 10-plex experiment in whole-mount zebrafish embryos without chromatic aberration correction (cf. Figure S6).** (A) Confocal images: individual channels; single optical section. Images not corrected for chromatic aberration. Solid boundary denotes region of variable expression; dashed boundary denotes region of no/low expression. Pixel size: 0.180×0.180×1.2 $\mu$m. Sample: Whole-mount zebrafish embryos; fixed 27 hpf. (B) Raw voxel intensity scatter plots representing signal plus background for voxels within solid boundary of panel A. Voxel size: 2.0×2.0×1.2 $\mu$m. Dashed lines represent BOT and TOP values (Table S10) used to normalize data for panel C using methods of Section S1.4.3. (C) Normalized voxel intensity scatter plots representing estimated normalized signal (Pearson correlation coefficient, $r$). Ch1: *col2a1a* (Alexa405). Ch10: *col2a1a* (iFluor800). Sample: Whole-mount zebrafish embryos; fixed 27 hpf.

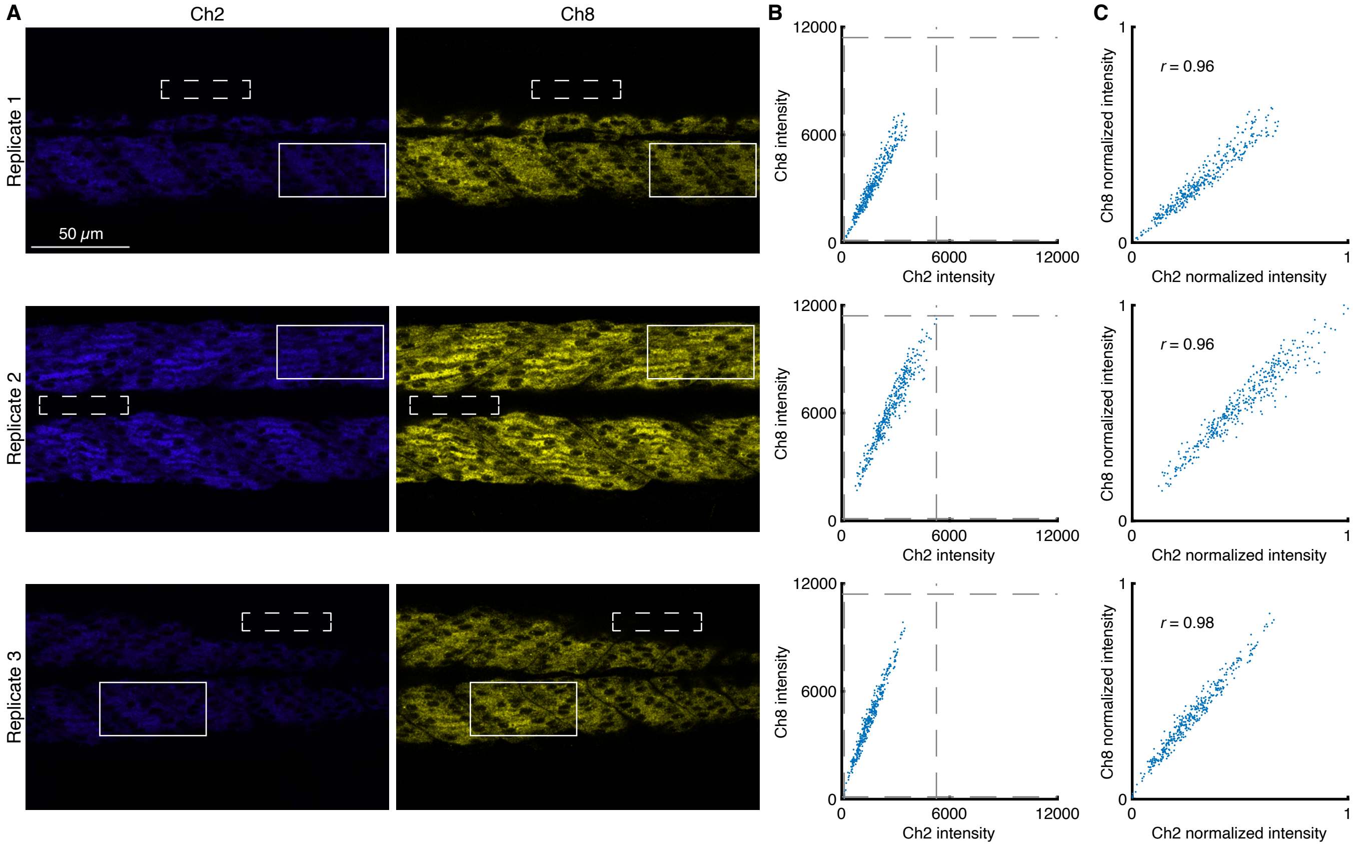

**Figure S12. 2-channel redundant detection of target mRNA *mylpfa* in the context of a 10-plex experiment in whole-mount zebrafish embryos without chromatic aberration correction (cf. Figure S7).** (A) Confocal images: individual channels; single optical section. Images not corrected for chromatic aberration. Solid boundary denotes region of variable expression; dashed boundary denotes region of no/low expression. Pixel size: $0.180 \times 0.180 \times 1.2$ $\mu$m. Sample: Whole-mount zebrafish embryos; fixed 27 hpf. (B) Raw voxel intensity scatter plots representing signal plus background for voxels within solid boundary of panel A. Voxel size: $2.0 \times 2.0 \times 1.2$ $\mu$m. Dashed lines represent BOT and TOP values (Table S10) used to normalize data for panel C using methods of Section S1.4.3. (C) Normalized voxel intensity scatter plots representing estimated normalized signal (Pearson correlation coefficient, $r$). Ch2: *mylpfa* (Atto425). Ch8: *mylpfa* (Alexa700). Sample: Whole-mount zebrafish embryos; fixed 27 hpf.

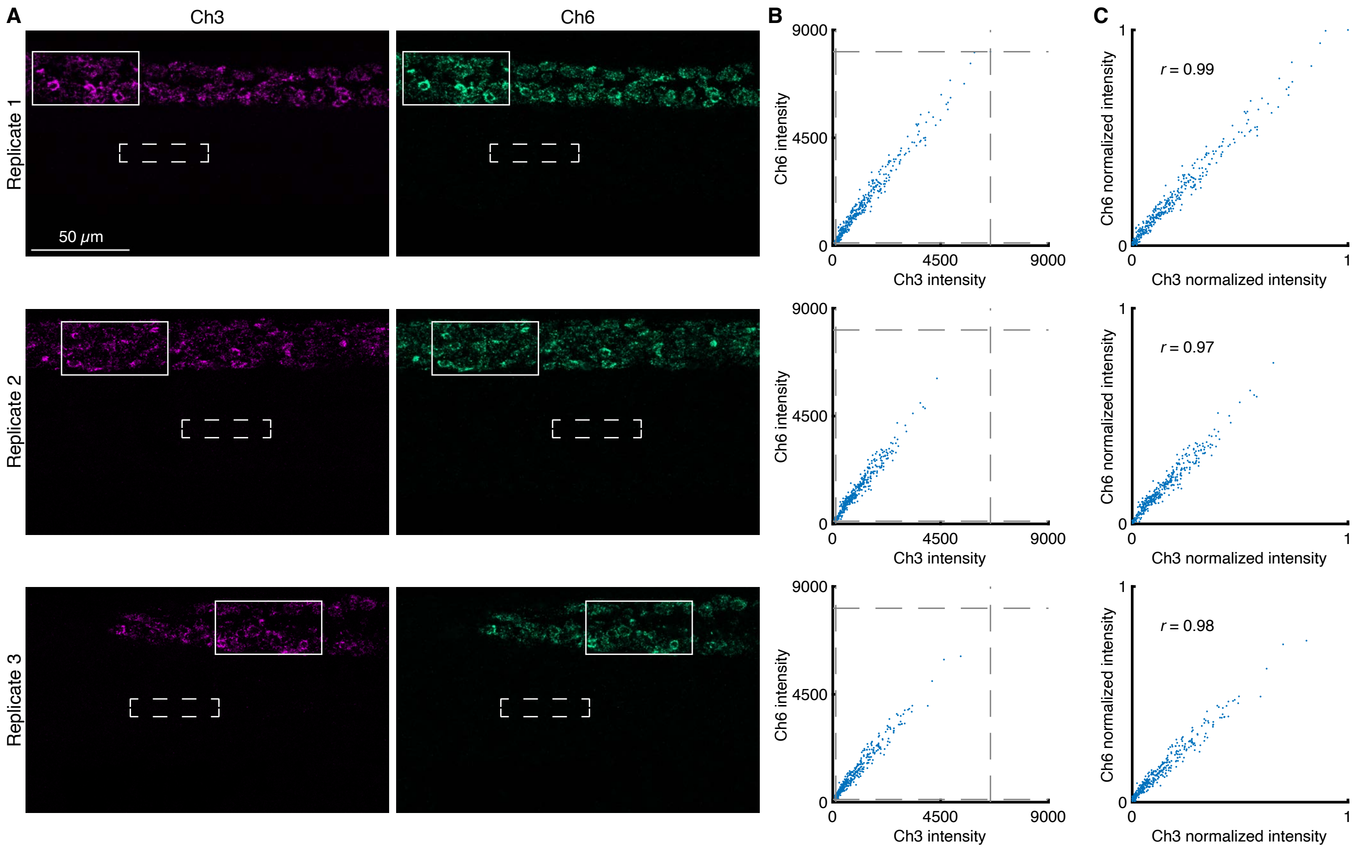

**Figure S13. 2-channel redundant detection of target mRNA *elavl3* in the context of a 10-plex experiment in whole-mount zebrafish embryos without chromatic aberration correction (cf. Figure S8).** (A) Confocal images: individual channels; single optical section. Images not corrected for chromatic aberration. Solid boundary denotes region of variable expression; dashed boundary denotes region of no/low expression. Pixel size: 0.180×0.180×1.2 μm. Sample: Whole-mount zebrafish embryos; fixed 27 hpf. (B) Raw voxel intensity scatter plots representing signal plus background for voxels within solid boundary of panel A. Voxel size: 2.0×2.0×1.2 μm. Dashed lines represent BOT and TOP values (Table S10) used to normalize data for panel C using methods of Section S1.4.3. (C) Normalized voxel intensity scatter plots representing estimated normalized signal (Pearson correlation coefficient, $r$). Ch3: *elavl3* (Alexa488). Ch6: *elavl3* (Alexa594). Sample: Whole-mount zebrafish embryos; fixed 27 hpf.

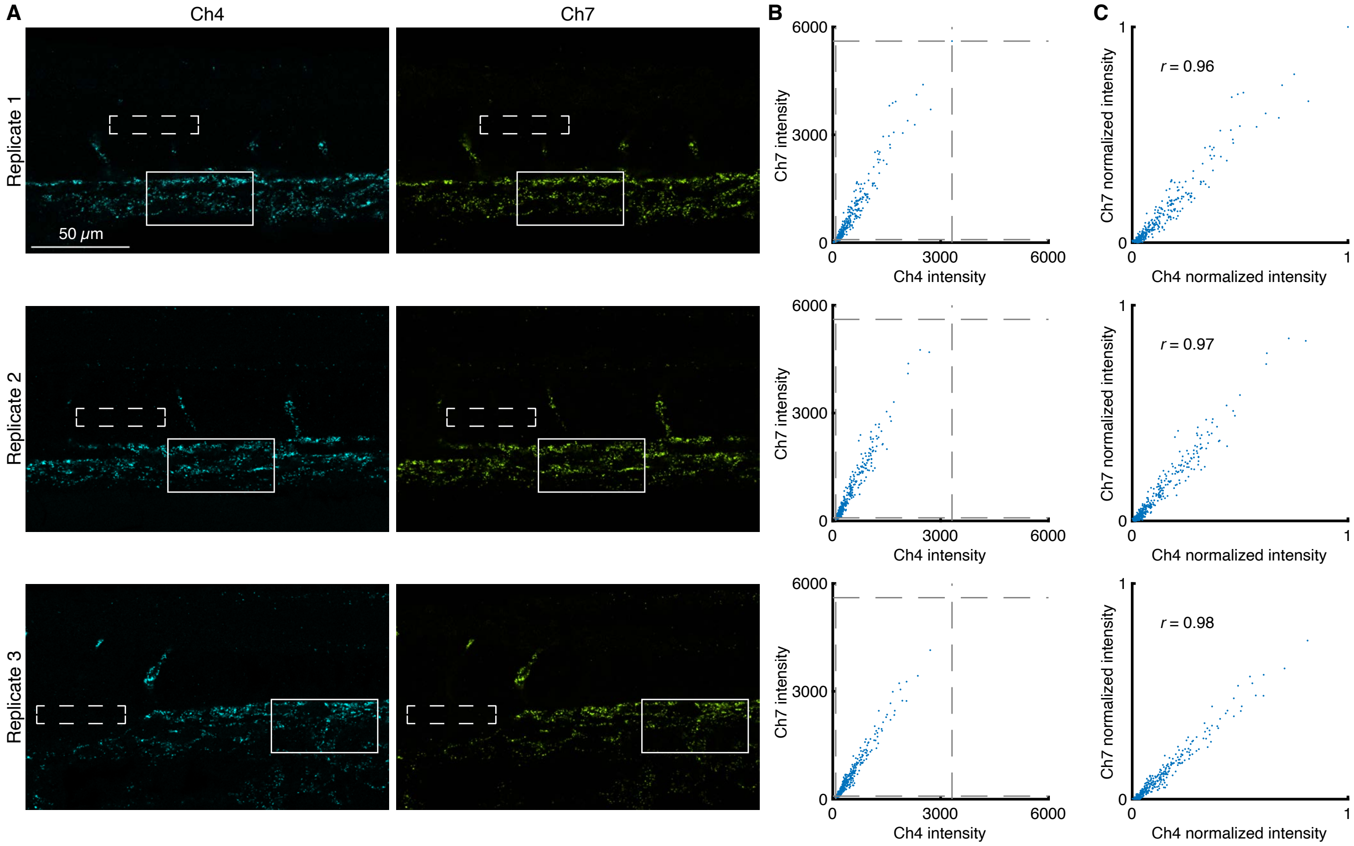

**Figure S14. 2-channel redundant detection of target mRNA *kdrl* in the context of a 10-plex experiment in whole-mount zebrafish embryos without chromatic aberration correction (cf. Figure S9).** (A) Confocal images: individual channels; single optical section. Images not corrected for chromatic aberration. Solid boundary denotes region of variable expression; dashed boundary denotes region of no/low expression. Pixel size: $0.180 \times 0.180 \times 1.2$ $\mu$m Sample: Whole-mount zebrafish embryos; fixed 27 hpf. (B) Raw voxel intensity scatter plots representing signal plus background for voxels within solid boundary of panel A. Voxel size: $2.0 \times 2.0 \times 1.2$ $\mu$m. Dashed lines represent BOT and TOP values (Table S10) used to normalize data for panel C using methods of Section S1.4.3. (C) Normalized voxel intensity scatter plots representing estimated normalized signal (Pearson correlation coefficient, $r$). Ch4: *kdrl* (Alexa514). Ch7: *kdrl* (Atto633). Sample: Whole-mount zebrafish embryos; fixed 27 hpf.

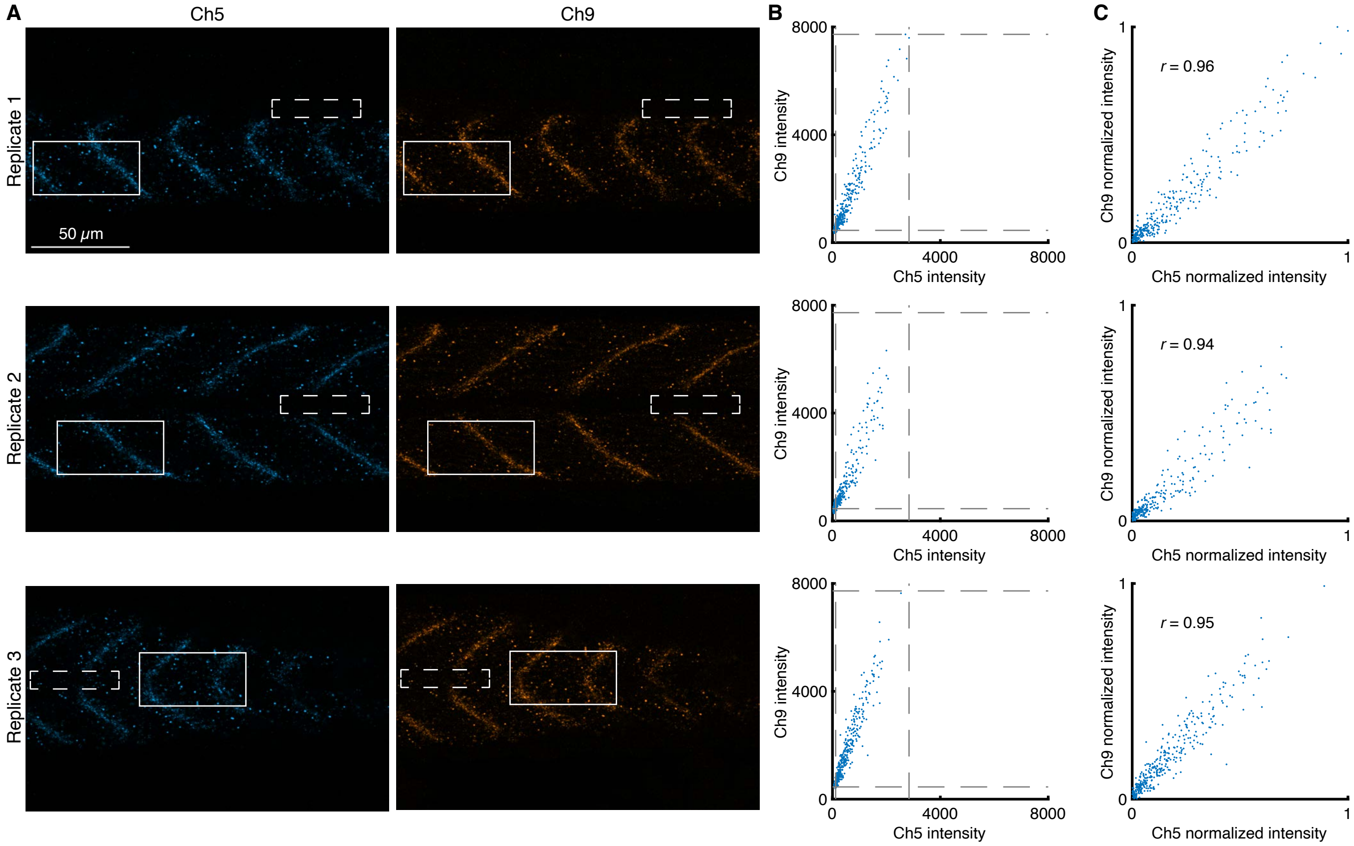

**Figure S15. 2-channel redundant detection of target mRNA *dmd* in the context of a 10-plex experiment in whole-mount zebrafish embryos without chromatic aberration correction (cf. Figure S10).** (A) Confocal images: individual channels; single optical section. Images not corrected for chromatic aberration. Solid boundary denotes region of variable expression; dashed boundary denotes region of no/low expression. Pixel size: $0.180 \times 0.180 \times 1.2$ $\mu$m. Sample: Whole-mount zebrafish embryos; fixed 27 hpf. (B) Raw voxel intensity scatter plots representing signal plus background for voxels within solid boundary of panel A. Voxel size: $2.0 \times 2.0 \times 1.2$ $\mu$m. Dashed lines represent BOT and TOP values (Table S10) used to normalize data for panel C using methods of Section S1.4.3. (C) Normalized voxel intensity scatter plots representing estimated normalized signal (Pearson correlation coefficient, $r$). Ch5: *dmd* (Alexa546). Ch9: *dmd* (Alexa750). Sample: Whole-mount zebrafish embryos; fixed 27 hpf.

| Channel | Target RNA | Fluorophore | BOT | TOP |
|---|---|---|---|---|
| Ch1 | *col2a1a* | Alexa405 | 140 | 10923 |
| Ch2 | *mylpfa* | Atto425 | 136 | 5263 |
| Ch3 | *elavl3* | Alexa488 | 126 | 6577 |
| Ch4 | *kdrl* | Alexa514 | 83 | 3316 |
| Ch5 | *dmd* | Alexa546 | 70 | 2832 |
| Ch6 | *elavl3* | Alexa594 | 112 | 8092 |
| Ch7 | *kdrl* | Atto633 | 85 | 5606 |
| Ch8 | *mylpfa* | Alexa700 | 127 | 11410 |
| Ch9 | *dmd* | Alexa750 | 286 | 7506 |
| Ch10 | *col2a1a* | iFluor800 | 108 | 14190 |

**Table S10. BOT and TOP values used to calculate normalized voxel intensities for scatter plots of Figures S11C–S15C without chromatic aberration correction.** Analysis based on rectangular regions depicted in Figures S11A–S15A using the methods of Section S1.4.3.

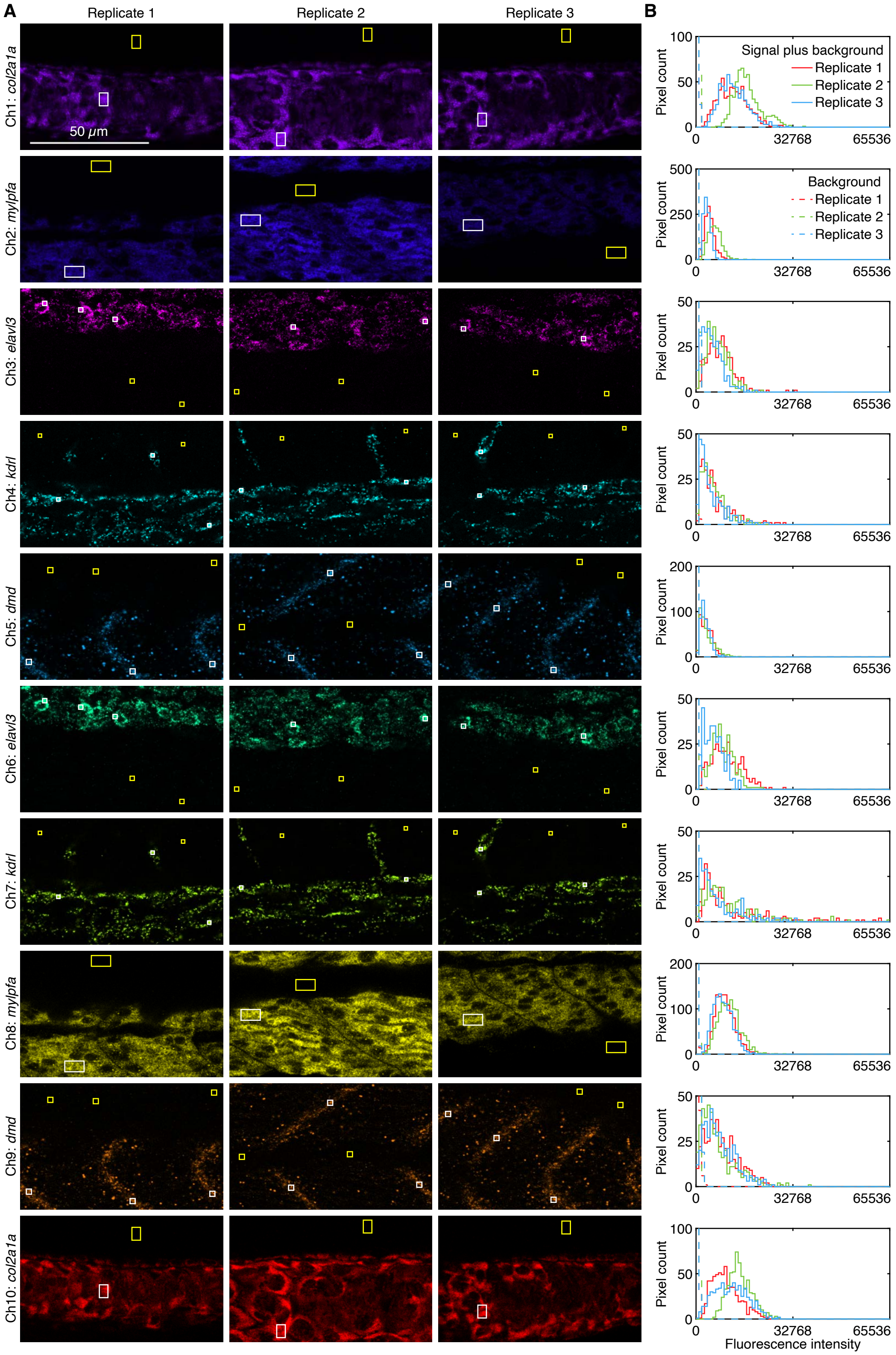

**Figure S16. Measurement of signal and background for 2-channel redundant detection of 5 target mRNAs in whole-mount zebrafish embryos (cf. Figure 3).** (A) Linearly unmixed fluorescence channels using Leica LAS X software; single optical section. For each of three replicate embryos, a representative single optical section was selected for each channel based on the expression of the corresponding target mRNA. Images not corrected for chromatic aberration. Solid white boundaries denote representative regions of high expression; solid yellow boundaries denote representative regions of no/low expression. (B) Pixel intensity histograms for signal plus background (pixels within solid white boundary in panel A) and background (pixels within solid yellow boundary in panel A). Ch1: *col2a1a* (Alexa405). Ch2: *mylpfa* (Atto425). Ch3: *elavl3* (Alexa488). Ch4: *kdrl* (Alexa514). Ch5: *dmd* (Alexa546). Ch6: *elavl3* (Alexa594). Ch7: *kdrl* (Atto633). Ch8: *mylpfa* (Alexa700). Ch9: *dmd* (Alexa750). Ch10: *col2a1a* (iFluor800). Sample: Whole-mount zebrafish embryos; fixed 27 hpf.

| Channel | Target RNA | Fluorophore | SIG+BACK | SIG | BACK | SIG/BACK |
|---|---|---|---|---|---|---|
| Ch1 | *col2a1a* | Alexa405 | $14\,000 \pm 2000$ | $14\,000 \pm 2000$ | $160 \pm 30$ | $80 \pm 20$ |
| Ch2 | *mylpfa* | Atto425 | $4400 \pm 800$ | $4300 \pm 800$ | $123 \pm 5$ | $35 \pm 7$ |
| Ch3 | *elavl3* | Alexa488 | $6500 \pm 900$ | $6300 \pm 900$ | $130 \pm 20$ | $47 \pm 9$ |
| Ch4 | *kdrl* | Alexa514 | $5300 \pm 400$ | $5200 \pm 400$ | $100 \pm 10$ | $50 \pm 8$ |
| Ch5 | *dmd* | Alexa546 | $3000 \pm 100$ | $2900 \pm 100$ | $70 \pm 10$ | $40 \pm 6$ |
| Ch6 | *elavl3* | Alexa594 | $8000 \pm 1000$ | $8000 \pm 1000$ | $110 \pm 30$ | $70 \pm 20$ |
| Ch7 | *kdrl* | Atto633 | $10\,000 \pm 1000$ | $10\,000 \pm 1000$ | $90 \pm 10$ | $110 \pm 20$ |
| Ch8 | *mylpfa* | Alexa700 | $9600 \pm 600$ | $9500 \pm 600$ | $70 \pm 10$ | $130 \pm 20$ |
| Ch9 | *dmd* | Alexa750 | $7600 \pm 400$ | $7300 \pm 400$ | $270 \pm 60$ | $27 \pm 6$ |
| Ch10 | *col2a1a* | iFluor800 | $12\,000 \pm 1000$ | $12\,000 \pm 1000$ | $110 \pm 20$ | $100 \pm 20$ |

**Table S11. Estimated signal-to-background for 2-channel redundant detection of 5 target mRNAs in whole-mount zebrafish embryos (cf. Figure 3).** Mean ± standard error of the mean (SEM), $N = 3$ replicate embryos. Analysis based on rectangular regions depicted in Figure S16 using methods of Section S1.4.2.

##### S3.4 dHCR imaging: RNA absolute quantitation in an anatomical context (cf. Figure 4)

Additional studies are presented as follows:

- Figure S17 displays 2-channel redundant detection of target mRNA *kdrl* for $N = 3$ replicate embryos. Images are annotated with single-molecule dots identified using the Dot Detection 2.0 software package.
- Table S12 displays the number of dots detected per channel and the colocalization fraction for $N = 3$ replicate embryos.
- Figure S18 displays 10-plex images for $N = 3$ replicate embryos linearly unmixed using the Leica LAS X software (cf. Ch4 and Ch7 displayed with single-molecule dots annotated in Figure S17).

**Protocol:** Spectral imaging and linear unmixing protocol of Section S2.1 using the HCR RNA-FISH protocol of Section S2.2.
**Sample:** Whole-mount zebrafish embryos; fixed 27 hpf.
**Reagents:** Table S1.
**Microscope settings:** Table S3.
**Dot detection and colocalization:** Section S1.4.4.

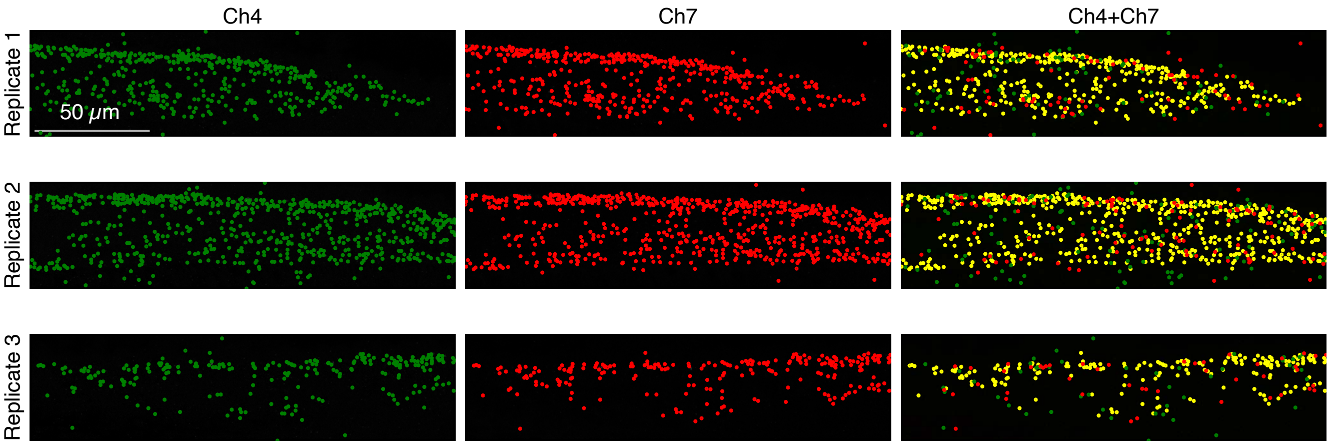

**Figure S17. dHCR imaging: digital RNA absolute quantitation in an anatomical context using 10-plex spectral imaging and linear unmixing (cf. Figure 4).** 2-channel redundant detection target mRNA *kdrl* alongside 2-channel redundant detection of 4 other target mRNAs in whole-mount zebrafish embryos. Linearly unmixed channels from a 10-plex confocal image in the dorsal posterior tail (Figure S18), where *kdrl* is expressed as single-molecule punctae. Maximum intensity z-projection for 5 optical sections (2.7 $\mu$m total); 0.18×0.18×1.2 µm pixels. Left: Ch4 (*kdrl*). Middle: Ch7 (*kdrl*). Right: Ch4+Ch7 merge. Green circles: dots detected in Ch4. Red circles: dots detected in Ch7. Yellow circles: dots detected in both channels. See Table S12 for colocalization rates. Sample: whole-mount zebrafish embryos; fixed 27 hpf.

| | Dots | | Colocalized dots | Colocalization fractions | |
|---|---|---|---|---|---|
| | $N_4$ | $N_7$ | $N_{47}$ | $C_4$ | $C_7$ |
| Replicate 1 | 453 | 445 | 364 | 0.804 | 0.818 |
| Replicate 2 | 666 | 644 | 538 | 0.808 | 0.835 |
| Replicate 3 | 235 | 232 | 187 | 0.796 | 0.806 |
| Mean | | | | $0.803 \pm 0.004$ | $0.820 \pm 0.009$ |

**Table S12. Dot colocalization fractions for redundant 2-channel detection of single *kdrl* mRNAs in whole-mount zebrafish embryos in the context of 10-plex spectral imaging with linear unmixing (cf. Figure 4).** The *kdrl* target RNA was detected in Ch4 and Ch7 as part of a 10-plex experiment. Colocalization rate indicates the fraction of dots in each channel that are detected in both channels (mean $\pm$ SEM for $N = 3$ replicate embryos). Analysis based on the images in Figure S17 using the methods of section S1.4.4.

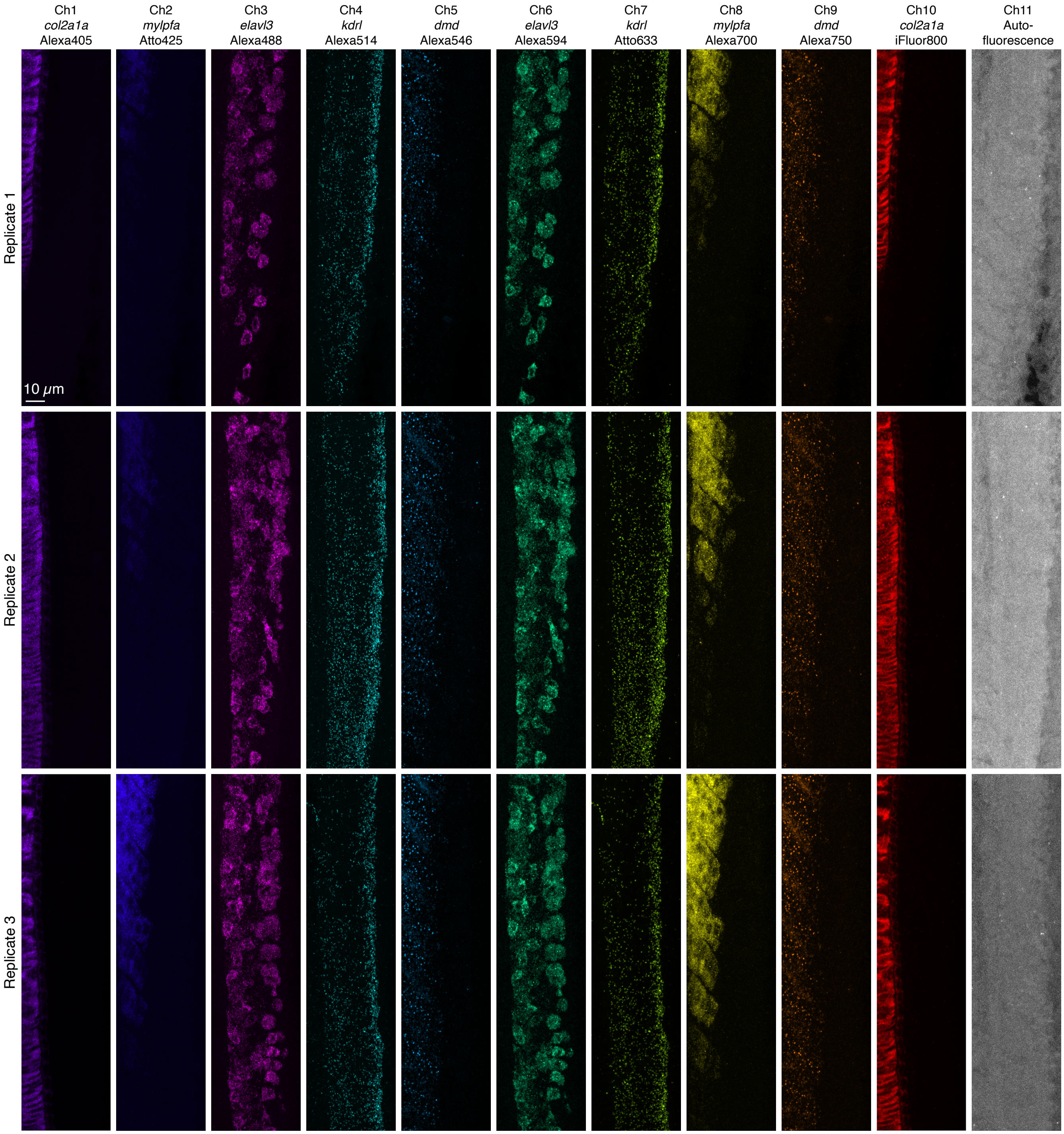

**Figure S18. Replicates for dHCR single-molecule RNA imaging in the context of 10-plex spectral imaging with linear unmixing in whole-mount zebrafish embryos (cf. Figure 4).** Linearly unmixed fluorescence channels using Leica LAS X software. Maximum-intensity z-projection for each channel. Replicate 1: 40 optical sections (16.1 $\mu$m total). Replicate 2: 45 optical sections (18.1 $\mu$m total). Replicate 3: 43 optical sections (17.3 $\mu$m total). $0.180 \times 0.180 \times 1.2$ $\mu$m pixels. Ch1: *col2a1a* (Alexa405). Ch2: *mylpfa* (Atto425). Ch3: *elavl3* (Alexa488). Ch4: *kdrl* (Alexa514). Ch5: *dmd* (Alexa546). Ch6: *elavl3* (Alexa594). Ch7: *kdrl* (Atto633). Ch8: *mylpfa* (Alexa700). Ch9: *dmd* (Alexa750). Ch10: *col2a1a* (iFluor800). Sample: whole-mount zebrafish embryos; fixed 27 hpf.

#### S3.5 Reference spectra, replicates, signal, and background for 10-plex RNA and protein imaging with high signal-to-background in fresh-frozen mouse brain sections (cf. Figure 5)

Additional studies are presented as follows:

- Figure S19 displays 11 raw and normalized reference spectra (1 per fluorophore and 1 for autofluorescence).
- Figures S20–S22 display 10-plex images for $N = 3$ replicate fresh-frozen mouse brain sections unmixed using the Leica LAS X software (cf. Figure 5C).
- Figure S23 displays representative regions of individual channels used for measurement of signal and background for each target.
- Table S13 displays estimated values for signal, background, and signal-to-background for each target.
- Figures S24–S26 display 10-plex images linearly unmixed using the Unmix 1.0 software package (cf. Figures S20–S22), providing an alternative for researchers that do not have access to the LAS X software.
- Figure S27 compares the pixel intensities for linear unmixing using the Leica LAS X software vs the Unmix 1.0 software package. For each channel and replicate, the pixel intensities are highly correlated between the two methods (Pearson correlation coefficient $r > 0.995$ for each fluorophore and $r > 0.986$ for autofluorescence).

**Protocol:** spectral imaging and linear unmixing protocol of Section S2.1 using the HCR RNA-FISH/IF protocol of Section S2.3.
**Sample:** fresh-frozen mouse brain section (coronal); 5 µm thickness; interaural region 0.88 mm ± 0.20 mm; 8 weeks old.
**Reagents:** Tables S1 and S2.
**Microscope settings:** Table S3.

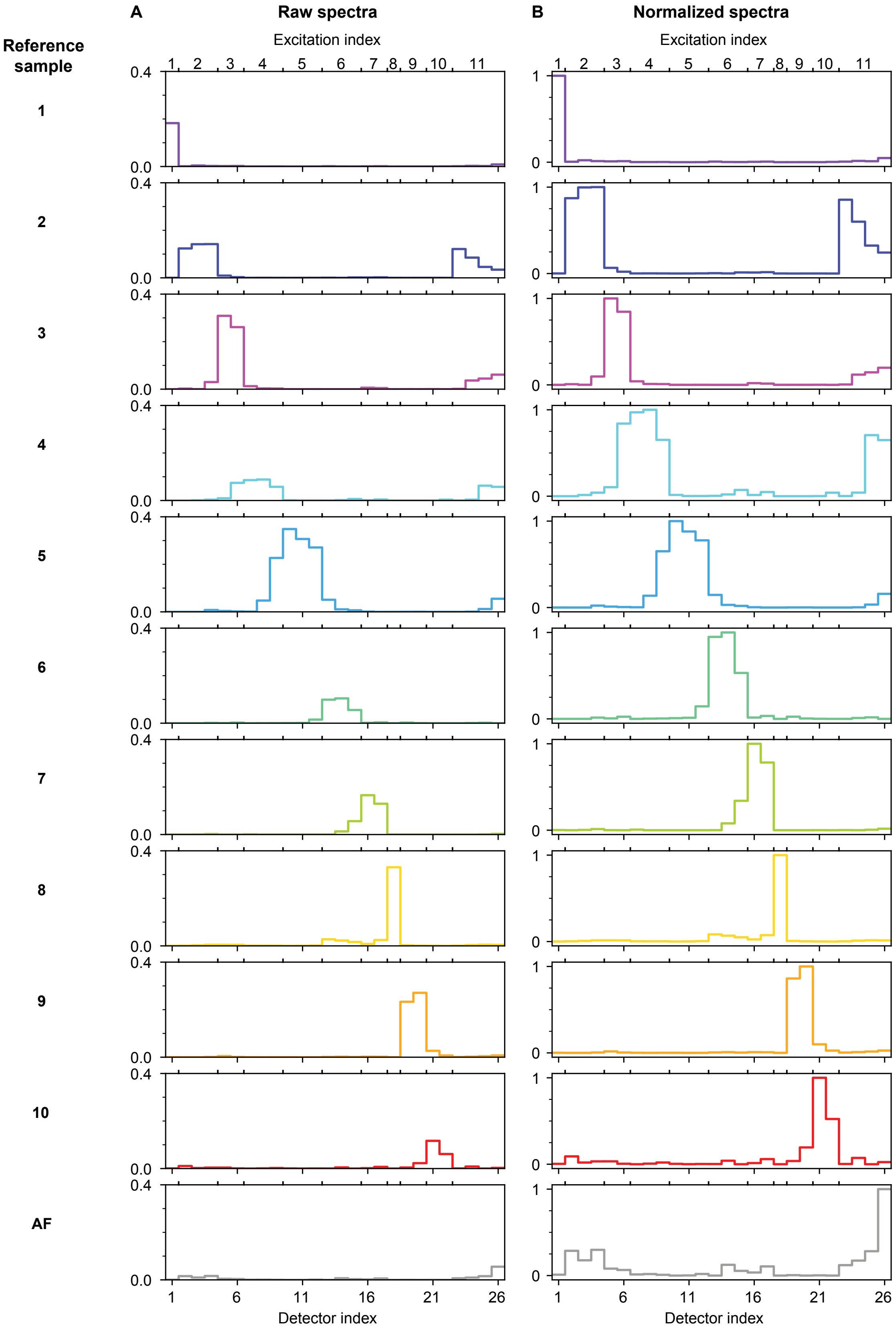

**Figure S19. 11 reference spectra in fresh-frozen mouse brain sections (1 per fluorophore and 1 for autofluorescence (AF); cf. Figure 1C for whole-mount zebrafish embryos).** (A) Raw reference spectra. (B) Normalized reference spectra (each spectrum scaled to have a maximum value of 1). Protocol: Section S2.1. Sample: fresh-frozen mouse brain section (coronal); 5 μm thickness; interaural region 0.88 mm ± 0.20 mm; 8 weeks old.

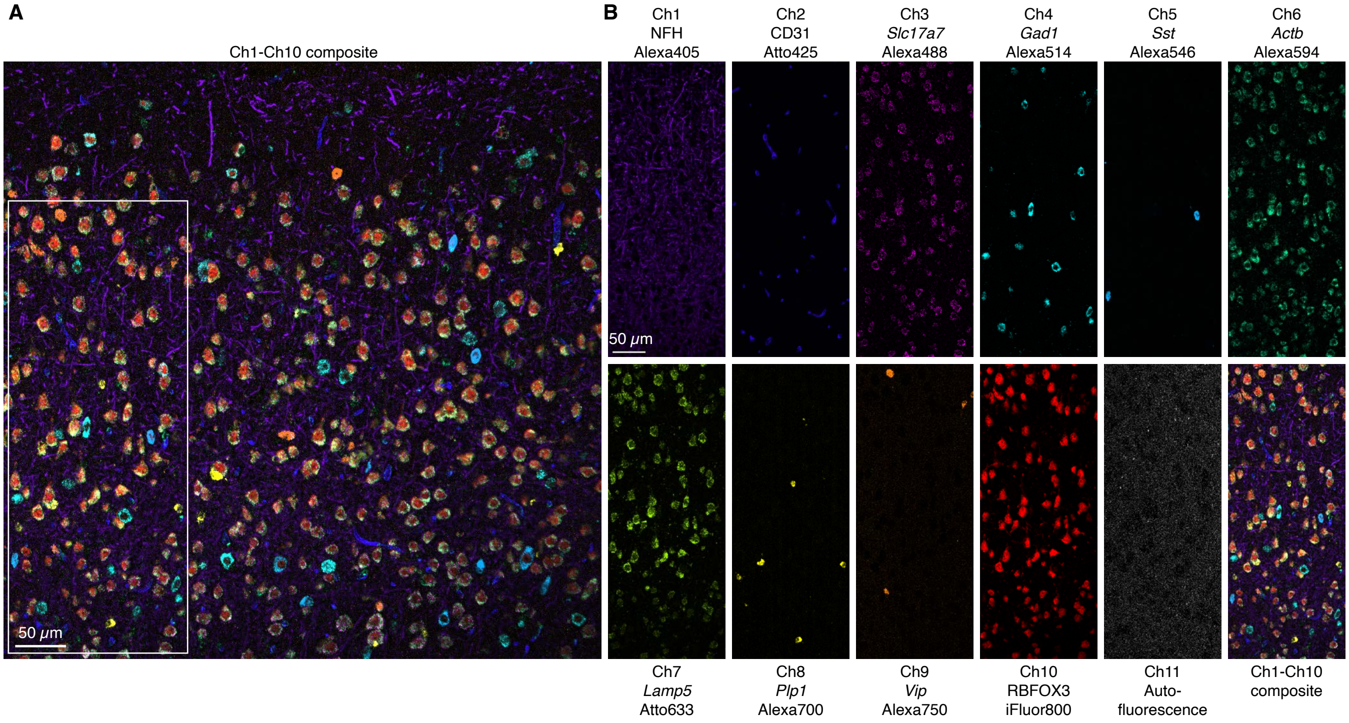

**Figure S20. Replicate 1 for 10-plex simultaneous RNA and protein imaging using HCR RNA-FISH/IF in a fresh-frozen mouse brain section (cf. Figure 5C).** Linearly unmixed fluorescence channels using Leica LAS X software. (A) Composite image of linearly unmixed Ch1-Ch10 for 3 protein targets and 7 mRNA targets in the cerebral cortex of a fresh-frozen mouse brain section; single optical section; $0.57 \times 0.57 \times 4.0$ µm pixels. (B) Right: single fluorescence channels for the depicted region in panel A (including autofluorescence as an 11th channel). Ch1: target protein NFH (Alexa405). Ch2: target protein CD31 (Atto425). Ch3: target RNA *Slc17a7* (Alexa488). Ch4: target RNA *Gad1* (Alexa514). Ch5: target RNA *Sst* (Alexa546). Ch6: target RNA *Actb* (Alexa594). Ch7: target RNA *Lamp5* (Atto633). Ch8: target RNA *Plp1* (Alexa700). Ch9: target RNA *Vip* (Alexa750). Ch10: target protein RBFOX3 (iFluor800). Sample: fresh-frozen mouse brain section (coronal); 5 µm thickness; interaural region 0.88 mm $\pm$ 0.20 mm; 8 weeks old.

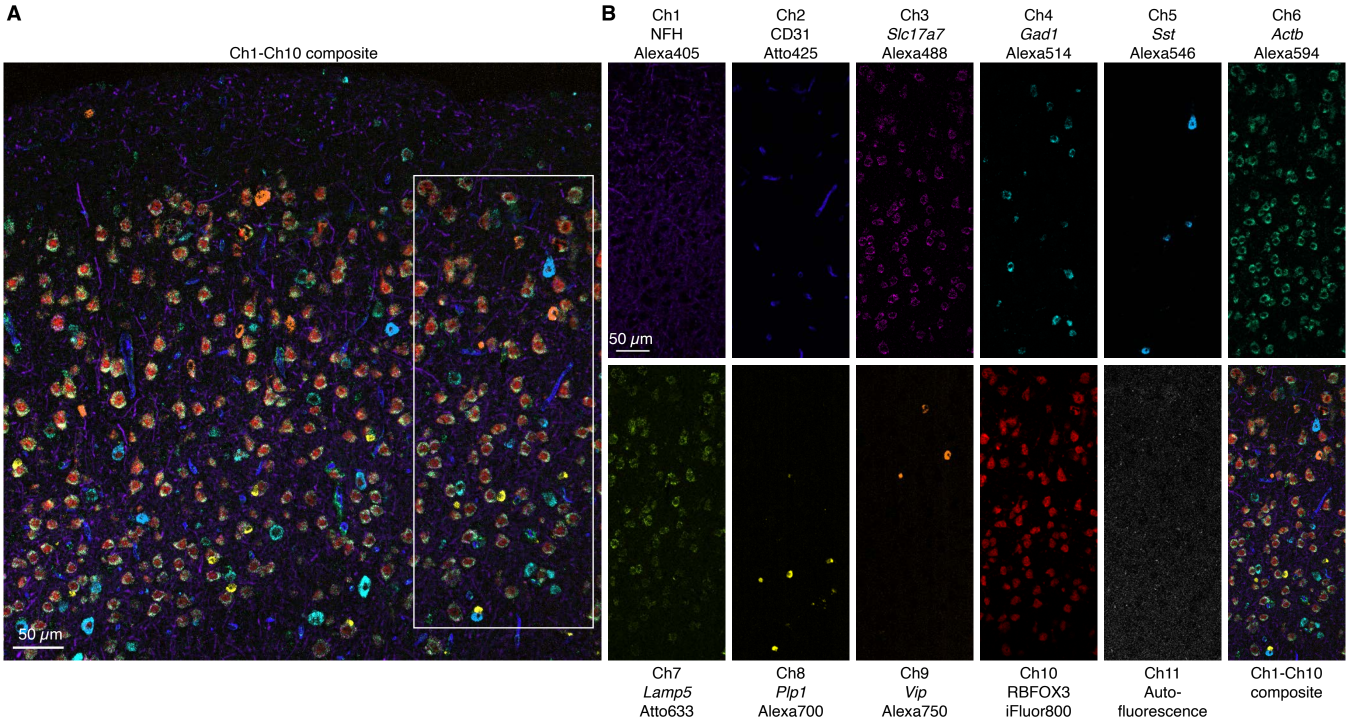

**Figure S21. Replicate 2 for 10-plex simultaneous RNA and protein imaging using HCR RNA-FISH/IF in a fresh-frozen mouse brain section (cf. Figure 5C).** Linearly unmixed fluorescence channels using Leica LAS X software. (A) Composite image of linearly unmixed Ch1-Ch10 for 3 protein targets and 7 mRNA targets in the cerebral cortex of a fresh-frozen mouse brain section; single optical section; 0.57×0.57×4.0 μm pixels. (B) Right: single fluorescence channels for the depicted region in panel A (including autofluorescence as an 11th channel). Ch1: target protein NFH (Alexa405). Ch2: target protein CD31 (Atto425). Ch3: target RNA *Slc17a7* (Alexa488). Ch4: target RNA *Gad1* (Alexa514). Ch5: target RNA *Sst* (Alexa546). Ch6: target RNA *Actb* (Alexa594). Ch7: target RNA *Lamp5* (Atto633). Ch8: target RNA *Plp1* (Alexa700). Ch9: target RNA *Vip* (Alexa750). Ch10: target protein RBFOX3 (iFluor800). Sample: fresh-frozen mouse brain section (coronal); 5 μm thickness; interaural region 0.88 mm ± 0.20 mm; 8 weeks old.

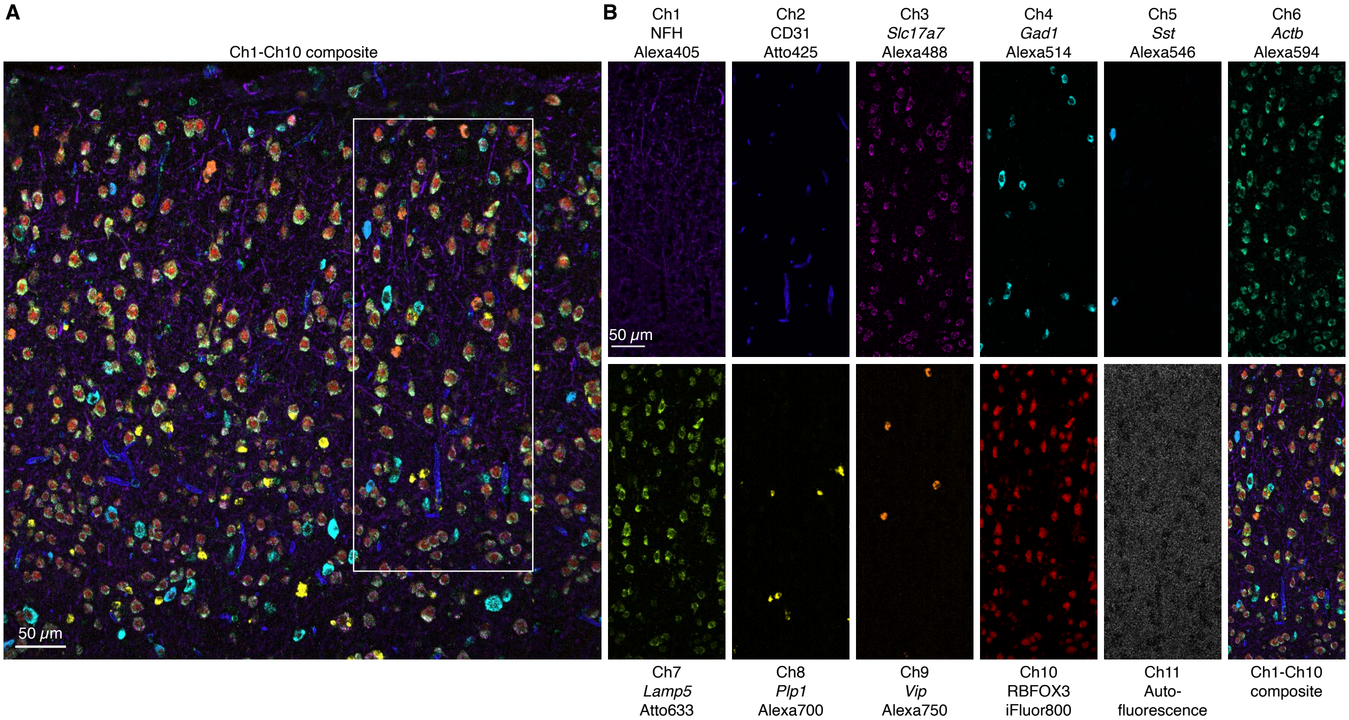

**Figure S22. Replicate 3 for 10-plex simultaneous RNA and protein imaging using HCR RNA-FISH/IF in a fresh-frozen mouse brain section (cf. Figure 5C).** Linearly unmixed fluorescence channels using Leica LAS X software. (A) Composite image of linearly unmixed Ch1-Ch10 for 3 protein targets and 7 mRNA targets in the cerebral cortex of a fresh-frozen mouse brain section; single optical section; $0.57\times0.57\times4.0$ µm pixels. (B) Right: single fluorescence channels for the depicted region in panel A (including autofluorescence as an 11th channel). Ch1: target protein NFH (Alexa405). Ch2: target protein CD31 (Atto425). Ch3: target RNA *Slc17a7* (Alexa488). Ch4: target RNA *Gad1* (Alexa514). Ch5: target RNA *Sst* (Alexa546). Ch6: target RNA *Actb* (Alexa594). Ch7: target RNA *Lamp5* (Atto633). Ch8: target RNA *Plp1* (Alexa700). Ch9: target RNA *Vip* (Alexa750). Ch10: target protein RBFOX3 (iFluor800). Sample: fresh-frozen mouse brain section (coronal); 5 µm thickness; interaural region 0.88 mm $\pm$ 0.20 mm; 8 weeks old.

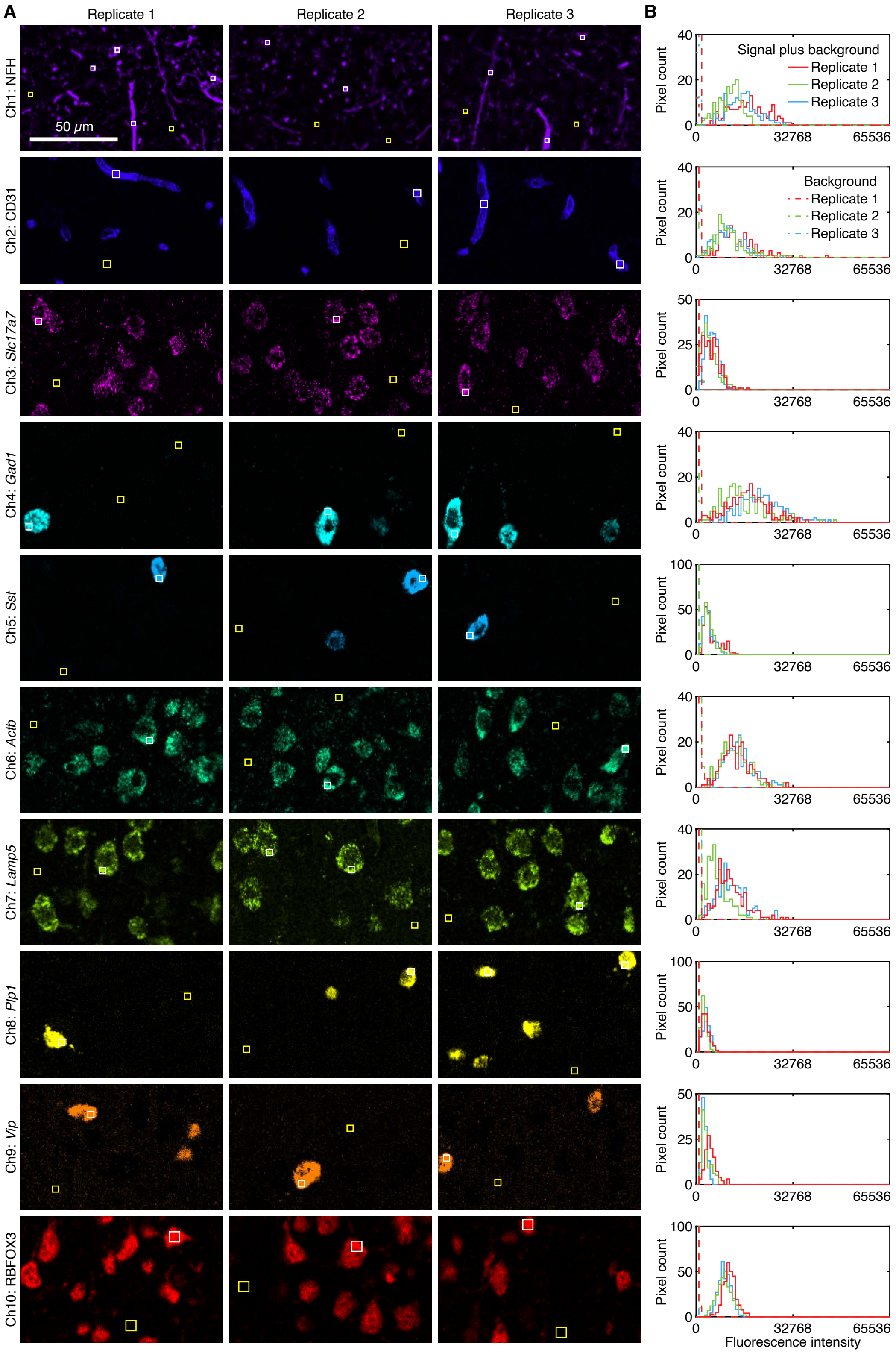

**Figure S23. Measurement of signal and background for 10-plex simultaneous RNA and protein imaging using HCR RNA-FISH/IF in fresh-frozen mouse brain sections (cf. Figure 5C).** (A) Linearly unmixed fluorescence channels using Leica LAS X software; single optical section; 0.57×0.57×4.0 μm pixels. (B) Pixel intensity histograms for signal plus background (pixels within solid white boundary in panel A) and background (pixels within solid yellow boundary in panel A). Confocal images collected with the microscope laser intensity and detector gain optimized to avoid saturating SIG + BACK pixels. Ch1: protein NFH (Alexa405). Ch2: protein CD31 (Atto425). Ch3: RNA *Slc17a7* (Alexa488). Ch4: RNA *Gad1* (Alexa514). Ch5: RNA *Sst* (Alexa546). Ch6: RNA *Actb* (Alexa594). Ch7: RNA *Lamp5* (Atto633). Ch8: RNA *Plp1* (Alexa700). Ch9: RNA *Vip* (Alexa750). Ch10: protein RBFOX3 (iFluor800). Sample: fresh-frozen mouse brain section (coronal); 5 μm thickness; interaural region 0.88 mm ± 0.20 mm; 8 weeks old.

| Channel | Target | Type | Fluorophore | SIG+BACK | SIG | BACK | SIG/BACK |
|---|---|---|---|---|---|---|---|
| Ch1 | NFH | protein | Alexa405 | $14\,000 \pm 2000$ | $14\,000 \pm 2000$ | $500 \pm 100$ | $26 \pm 7$ |
| Ch2 | CD31 | protein | Atto425 | $11\,500 \pm 900$ | $11\,300 \pm 900$ | $200 \pm 30$ | $55 \pm 9$ |
| Ch3 | *Slc17a7* | RNA | Alexa488 | $4500 \pm 300$ | $4300 \pm 300$ | $170 \pm 30$ | $25 \pm 4$ |
| Ch4 | *Gad1* | RNA | Alexa514 | $18\,000 \pm 2000$ | $18\,000 \pm 2000$ | $170 \pm 20$ | $110 \pm 20$ |
| Ch5 | *Sst* | RNA | Alexa546 | $4100 \pm 300$ | $4100 \pm 300$ | $28 \pm 2$ | $140 \pm 20$ |
| Ch6 | *Actb* | RNA | Alexa594 | $13\,200 \pm 400$ | $12\,900 \pm 400$ | $380 \pm 50$ | $34 \pm 4$ |
| Ch7 | *Lamp5* | RNA | Atto633 | $10\,000 \pm 2000$ | $10\,000 \pm 2000$ | $310 \pm 80$ | $30 \pm 9$ |
| Ch8 | *Plp1* | RNA | Alexa700 | $2700 \pm 300$ | $2700 \pm 300$ | $27 \pm 4$ | $100 \pm 20$ |
| Ch9 | *Vip* | RNA | Alexa750 | $3300 \pm 600$ | $3200 \pm 600$ | $39 \pm 8$ | $80 \pm 20$ |
| Ch10 | RBFOX3 | protein | iFluor800 | $9800 \pm 600$ | $9700 \pm 600$ | $101 \pm 7$ | $96 \pm 8$ |

**Table S13. Estimated signal-to-background for 10-plex simultaneous RNA and protein imaging using HCR RNA-FISH/IF in fresh-frozen mouse brain sections (cf. Figure 5C).** Mean $\pm$ estimated standard error of the mean via uncertainty propagation for $N = 3$ replicate brain sections. Analysis based on representative rectangular regions depicted in Figure S23 using methods of Section S1.4.2.

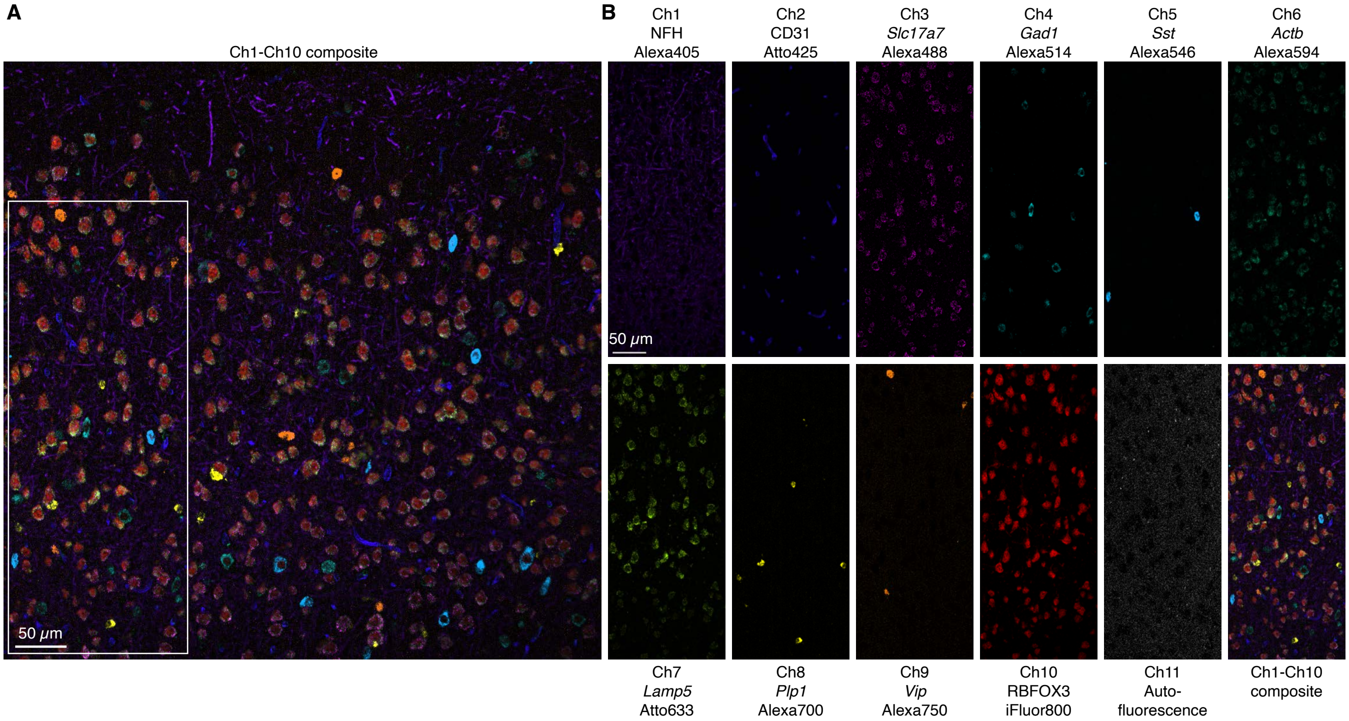

**Figure S24. Replicate 1 for 10-plex simultaneous RNA and protein imaging using HCR RNA-FISH/IF in a fresh-frozen mouse brain section linearly unmixed using the Unmix 1.0 software package (cf. Figure S20 linearly unmixed using the Leica LAS X software).** (A) Composite image of linearly unmixed Ch1-Ch10 for 3 protein targets and 7 mRNA targets in the cerebral cortex of a fresh-frozen mouse brain section; single optical section; 0.57×0.57×4.0 μm pixels. (B) Right: single fluorescence channels for the depicted region in panel A (including autofluorescence as an 11th channel). Ch1: target protein NFH (Alexa405). Ch2: target protein CD31 (Atto425). Ch3: target RNA *Slc17a7* (Alexa488). Ch4: target RNA *Gad1* (Alexa514). Ch5: target RNA *Sst* (Alexa546). Ch6: target RNA *Actb* (Alexa594). Ch7: target RNA *Lamp5* (Atto633). Ch8: target RNA *Plp1* (Alexa700). Ch9: target RNA *Vip* (Alexa750). Ch10: target protein RBFOX3 (iFluor800). Sample: fresh-frozen mouse brain section (coronal); 5 μm thickness; interaural region 0.88 mm ± 0.20 mm; 8 weeks old.

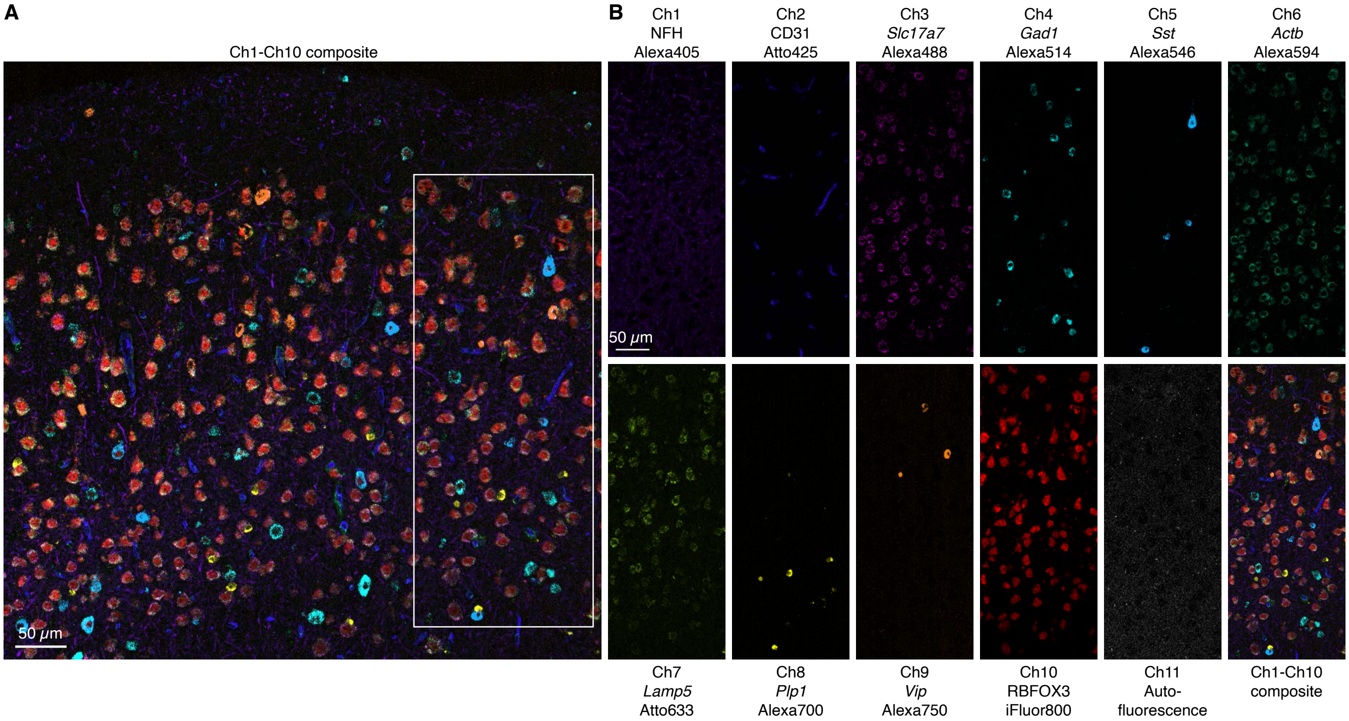

**Figure S25. Replicate 2 for 10-plex simultaneous RNA and protein imaging using HCR RNA-FISH/IF in a fresh-frozen mouse brain section linearly unmixed using the Unmix 1.0 software package (cf. Figure S21 linearly unmixed using the Leica LAS X software).** (A) Composite image of linearly unmixed Ch1-Ch10 for 3 protein targets and 7 mRNA targets in the cerebral cortex of a fresh-frozen mouse brain section; single optical section; 0.57×0.57×4.0 μm pixels. (B) Right: single fluorescence channels for the depicted region in panel A (including autofluorescence as an 11th channel). Ch1: target protein NFH (Alexa405). Ch2: target protein CD31 (Atto425). Ch3: target RNA *Slc17a7* (Alexa488). Ch4: target RNA *Gad1* (Alexa514). Ch5: target RNA *Sst* (Alexa546). Ch6: target RNA *Actb* (Alexa594). Ch7: target RNA *Lamp5* (Atto633). Ch8: target RNA *Plp1* (Alexa700). Ch9: target RNA *Vip* (Alexa750). Ch10: target protein RBFOX3 (iFluor800). Sample: fresh-frozen mouse brain section (coronal); 5 μm thickness; interaural region 0.88 mm ± 0.20 mm; 8 weeks old.

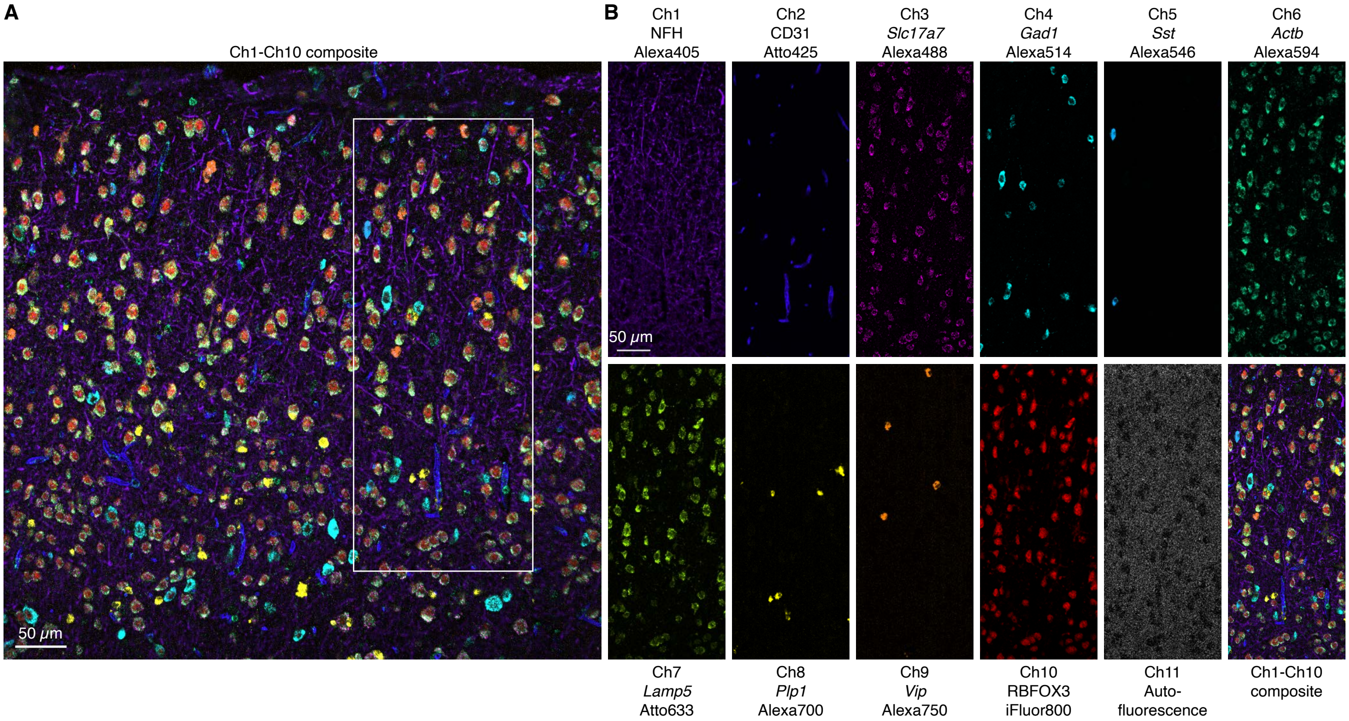

**Figure S26. Replicate 3 for 10-plex simultaneous RNA and protein imaging using HCR RNA-FISH/IF in a fresh-frozen mouse brain section linearly unmixed using the Unmix 1.0 software package (cf. Figure S22 linearly unmixed using the Leica LAS X software).** (A) Composite image of linearly unmixed Ch1-Ch10 for 3 protein targets and 7 mRNA targets in the cerebral cortex of a fresh-frozen mouse brain section; single optical section; 0.57×0.57×4.0 μm pixels. (B) Right: single fluorescence channels for the depicted region in panel A (including autofluorescence as an 11th channel). Ch1: target protein NFH (Alexa405). Ch2: target protein CD31 (Atto425). Ch3: target RNA *Slc17a7* (Alexa488). Ch4: target RNA *Gad1* (Alexa514). Ch5: target RNA *Sst* (Alexa546). Ch6: target RNA *Actb* (Alexa594). Ch7: target RNA *Lamp5* (Atto633). Ch8: target RNA *Plp1* (Alexa700). Ch9: target RNA *Vip* (Alexa750). Ch10: target protein RBFOX3 (iFluor800). Sample: fresh-frozen mouse brain section (coronal); 5 μm thickness; interaural region 0.88 mm ± 0.20 mm; 8 weeks old.

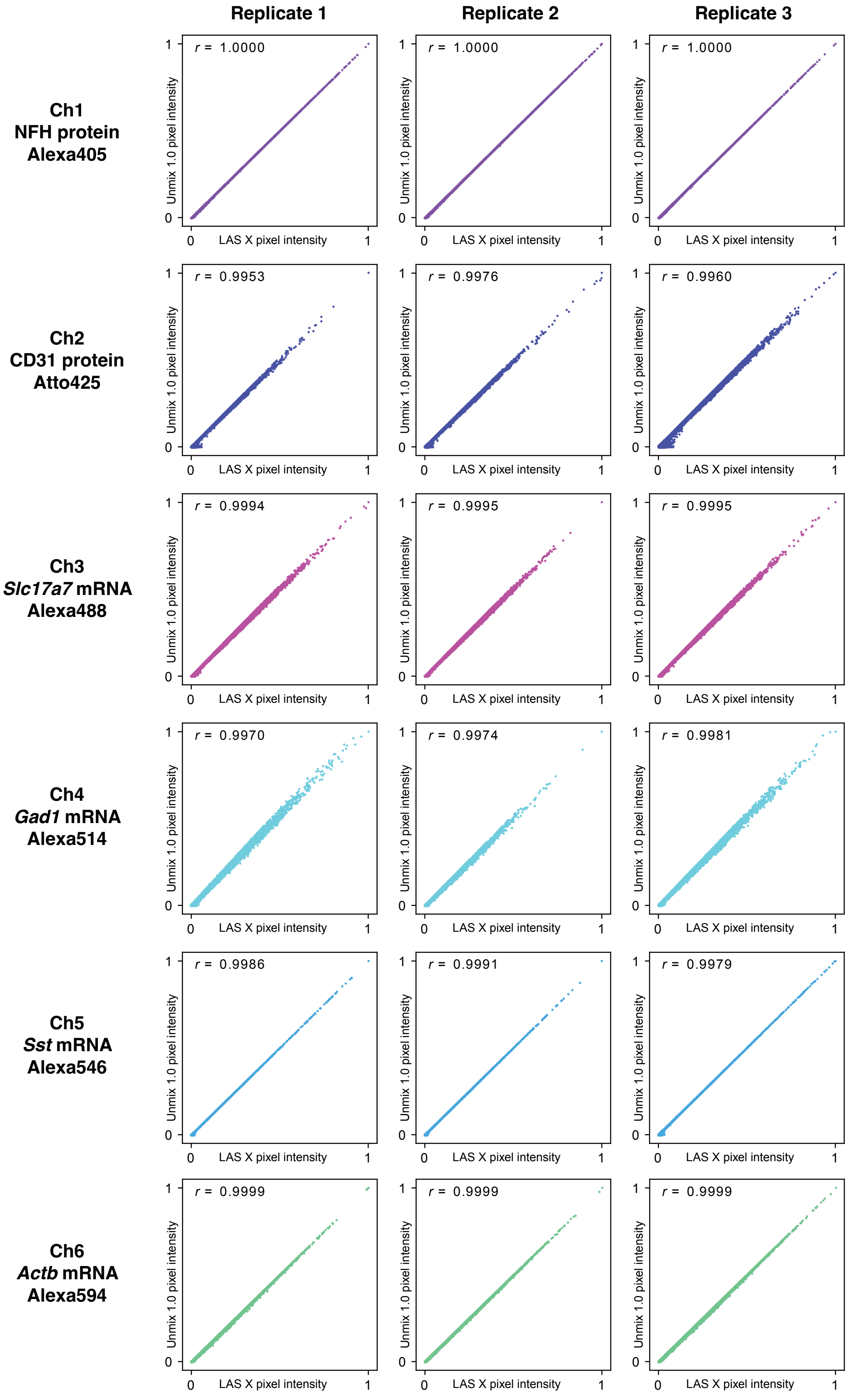
Replicate 1
Replicate 2
Replicate 3
Ch1
NFH protein
Alexa405
Ch2
CD31 protein
Atto425
Ch3
Slc17a7 mRNA
Alexa488
Ch4
Gad1 mRNA
Alexa514
Ch5
Sst mRNA
Alexa546
Ch6
Actb mRNA
Alexa594
r = 1.0000
r = 1.0000
r = 1.0000
r = 0.9953
r = 0.9976
r = 0.9960
r = 0.9994
r = 0.9995
r = 0.9995
r = 0.9970
r = 0.9974
r = 0.9981
r = 0.9986
r = 0.9991
r = 0.9979
r = 0.9999
r = 0.9999
r = 0.9999
Unmix 1.0 pixel intensity
LAS X pixel intensity
0
1

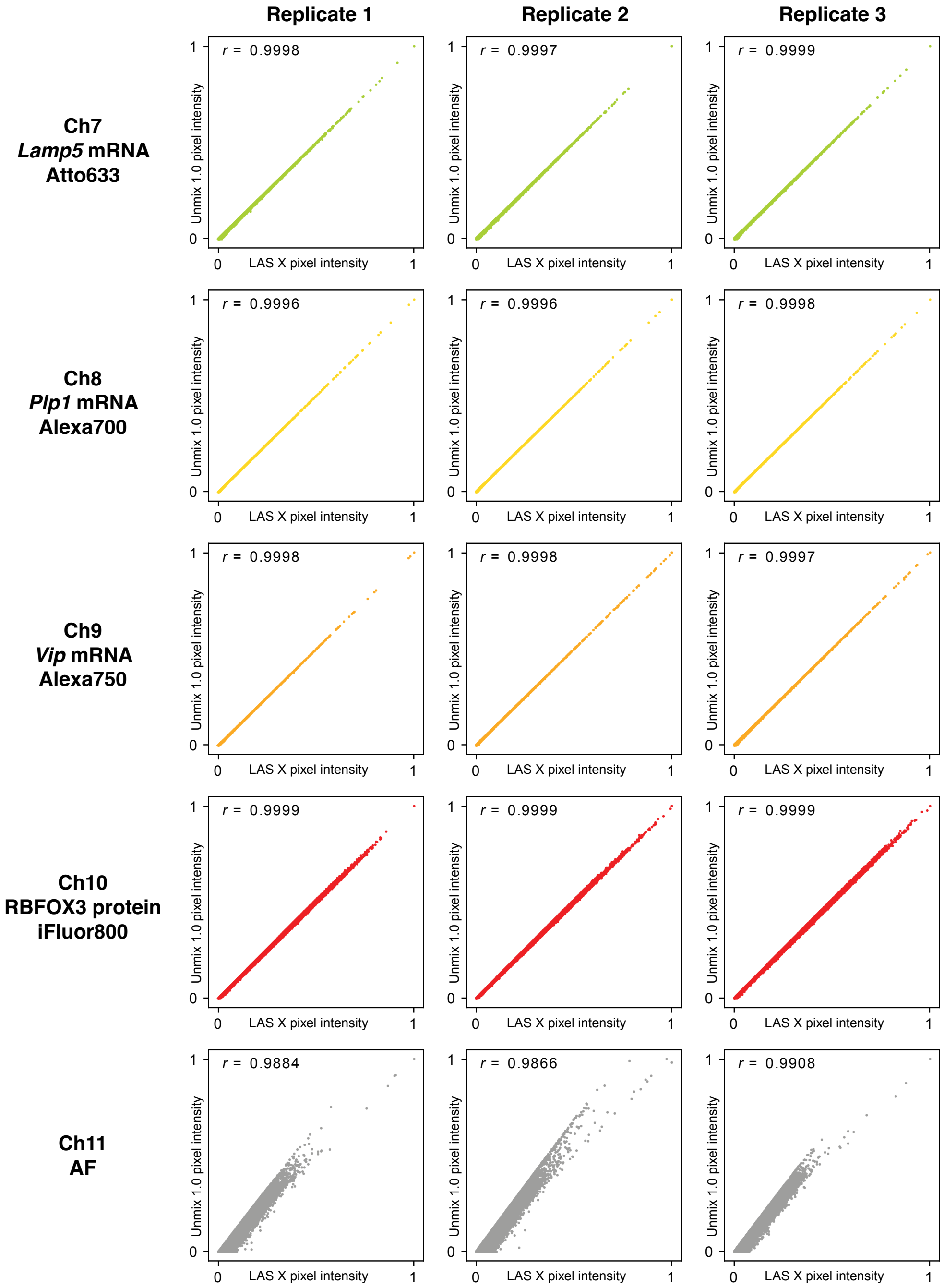

**Figure S27. Comparison of pixel intensities for linear unmixing using Leica LAS X software vs Unmix 1.0 software for 10-plex RNA and protein imaging in fresh-frozen mouse brain sections.** Pixel intensity scatter plots for replicate images of Figures S20A–S22A (Leica LAS X software) and Figures S24A–S26A (Unmix 1.0 software). Single optical section; $0.57 \times 0.57 \times 4.0$ µm pixels. Each scatter plot contains 1,048,576 dots corresponding to $1024 \times 1024$ pixels. Pixel intensities normalized so that the maximum intensity for each axis is 1. Pearson correlation coefficient, $r$.
